# *De novo* synthesis of fatty acids in Archaea via an archaeal fatty acid synthase complex

**DOI:** 10.1101/2024.07.05.601840

**Authors:** Christian Schmerling, Xiaoxiao Zhou, Paul E. Görs, Stephan Köstlbacher, Till Kessenbrock, David Podlesainski, David Sybers, Kun Wang, Ann-Christin Lindås, Jacky L. Snoep, Eveline Peeters, Markus Kaiser, Thijs J.G. Ettema, Sven W. Meckelmann, Christopher Bräsen, Bettina Siebers

**Author notes:** Address correspondence to: Christopher Bräsen. Christian Schmerling and Xiaoxiao Zhou contributed equally to this work.

## Abstract

Archaea synthesize membranes using isoprenoid-based ether lipids, whereas Bacteria and Eukarya use *fatty* acid-based ester lipids. While the factors responsible for this “lipid divide” remain unclear, this has important implications for understanding the evolutionary history of eukaryotes, which likely originated from within the Archaea and therefore changed membrane composition from isoprenoid-based to fatty acid-based lipids. Here, using ^13^C labelling studies, we demonstrate that the archaeal model organisms *Sulfolobus acidocaldarius* and *Haloferax volcanii* are capable of *de novo* fatty acid synthesis. Biochemical characterization and *in vitro* pathway reconstitution identify the key enzymes of a newly proposed fatty acid synthesis pathway in *S. acidocaldarius* and show that ketothiolase, ketoacyl-CoA reductase, and hydroxyacyl-CoA dehydratase form a stable assembly mediated by a DUF35 domain protein, which represents the first characterization of an archaeal fatty acid synthase complex. The final step is catalysed by an NADPH-dependent enoyl-CoA reductase. Deletion of the enoyl-CoA reductase demonstrate that this pathway operates *in vivo* in S*. acidocaldarius*. The presented results including phylogenetic analysis reveal that the potential to synthesize fatty acids is widespread across archaeal lineages. Collectively, our findings demonstrate that archaea are capable of synthesizing fatty acids, elucidate the molecular mechanisms involved in this process and provide additional insights into the evolutionary histories of fatty acid synthesis in archaea.

## Main text

One of the most striking differences between Archaea and the other domains of life (Bacteria and Eukarya) is the composition of their lipid membranes, which has been referred to as the “lipid divide” (*1–3*). Archaea have characteristic membrane lipids comprised of isoprenoid chains ether-linked to glycerol 1-phosphate (G1P) forming lipid mono-or bilayers, which are fundamentally different from those found in Bacteria and Eukaryotes which are composed of fatty acids ester-linked to glycerol 3-phosphate (G3P) forming membrane bilayers. Interest in this “lipid divide” has been revived by recent studies that support that eukaryotes likely originated from within the Archaea, i.e. from the Asgardarcheota (*4*), which suggests that eukaryogenesis involved a transition in the membrane composition of cells from archaeal (isoprenoid-based) to bacterial (fatty acid-based) lipids. However, the evolutionary history behind the “lipid divide” and the transition during the evolution of eukaryotic cells remains unclear (*3, 5, 6*).

Despite these differences, some studies have suggested that fatty acids are present in a few Archaea, such as *Sulfolobus* spp. (*5, 7*). However, no studies to date have been able to conclusively demonstrate fatty acid biosynthesis in archaea or define the pathways that would enable their production, raising questions on whether the presence of these molecules in archaeal cells results from *de novo* synthesis, importation and/or contamination. Notably, archaea do not seem to encode neither a complete classical type II fatty acid synthase (FAS II) system (present in bacteria) nor a type I fatty acid synthase (FAS I) system (present in animals and fungi) that could support fatty acid synthesis (*5*). Furthermore, the acyl carrier protein (ACP) and the ACP synthase, which are essential for fatty acid biosynthesis in Bacteria and Eukarya, are absent from nearly all Archaea (*8*). Thus, if some archaea are able to synthesize fatty acids, their synthesis machinery must be fundamentally different from what is found in other organisms, including by being ACP-independent (*5, 8*).

One hypothesis that has been proposed to potentially enable fatty acid synthesis in Archaea is the existence of a reversed β oxidation pathway. Under this scenario, the same enzymes that are used for fatty acid degradation in Archaea may also be able to catalyze the reverse reactions that would enable fatty acid assembly. This hypothesis is supported by studies showing that some archaea (such as *Natronomonas pharaonis*, *Archaeoglobus fulgidus* and *S. acidocaldarius*) can use fatty acids as carbon and energy source, and the observation that several archaeal genomes encode complete sets of bacterial-like β oxidation homologues (*5, 9, 10*). However, so far no studies have been able to demonstrate that β oxidation is functional in Archaea, either to degrade or synthesize fatty acids. Furthermore, although some reactions of the β oxidation pathway may operate reversibly under certain conditions, a complete reversal of the pathway has never been demonstrated in any organism, and various regulatory and thermodynamic constrains are thought to contribute to separate both processes (see “Supplementary Text”, Fig. S1). Therefore, whether and how Archaea can synthesize fatty acids, and how they degrade these molecules, remains unclear.

Here, we address these questions by investigating fatty acid metabolism in the archaeal model organisms *S. acidocaldarius* and *H. volcanii.* First, using ^13^C-labelled glycerol combined with a newly established methodology for fatty acid detection (*11*), we show that these distantly related Archaea are both capable of *de novo* fatty acid synthesis. Then, we characterize the function of the selected enzymes involved in β oxidation in *S. acidocaldarius*, demonstrating that while they mediate fatty acid degradation, the pathway is likely irreversible and does not support fatty acid synthesis. We then identify and functionally characterize the four key enzymes of a proposed novel ACP-independent fatty acid synthesis pathway, and support the *in vivo* significance of this pathway using mutant analyses. Furthermore, we demonstrate that three of the four pathway enzymes form a stable assembly mediated by a DUF35 domain protein that represents the first example of an archaeal fatty acid synthase complex. Finally, we use phylogenetic analysis to demonstrate that this fatty acid synthesis pathway is likely present in additional archaeal lineages, suggesting – in combination with the demonstrated presence of fatty acids in *S. acidocaldarius* and *H. volcanii*-that the ability to make fatty acids is widespread within this domain.

## Results

### Synthesis of fatty acids in *S. acidocaldarius* and *H. volcanii*

To elucidate the capability of *de-novo* fatty acid synthesis in Archaea, we tested our ability to detect ^13^C-labelled fatty acids in archaeal cells upon growth on a ^13^C-labelled substrate as sole carbon source, to exclude the possibility that the fatty acids detected in these cells are imported from other sources and/or contaminants. The archaeal model organisms *S. acidocaldarius* and *H. volcanii* were grown on ^13^C-labeled glycerol for three passages. In *S. acidocaldarius*, fatty acid detection (*11*) revealed mainly ^13^C8:0 and ^13^C10:0 fatty acids (both 170-180 fmol mg^-1^ cell wet weight), and to a lesser extend ^13^C12:0 fatty acids. In *H. volcanii*, we mainly detected ^13^C16:0 fatty acids (700 fmol mg^-1^ cell wet weight), with trace amounts of ^13^C10:0, ^13^C12:0, ^13^C14:0, and ^13^C18:0 fatty acids (Fig. 1, Fig. S2-5). Furthermore, in *S. acidocaldarius,* no fatty acids were detected when cells were grown on ^13^C-labelled D-xylose, indicating that fatty acid formation is dependent on the growth substrate (Fig. S6). Collectively, these data demonstrate the *de-novo* synthesis of fatty acids in organisms from distant archaeal phyla.

**Fig. 1.**
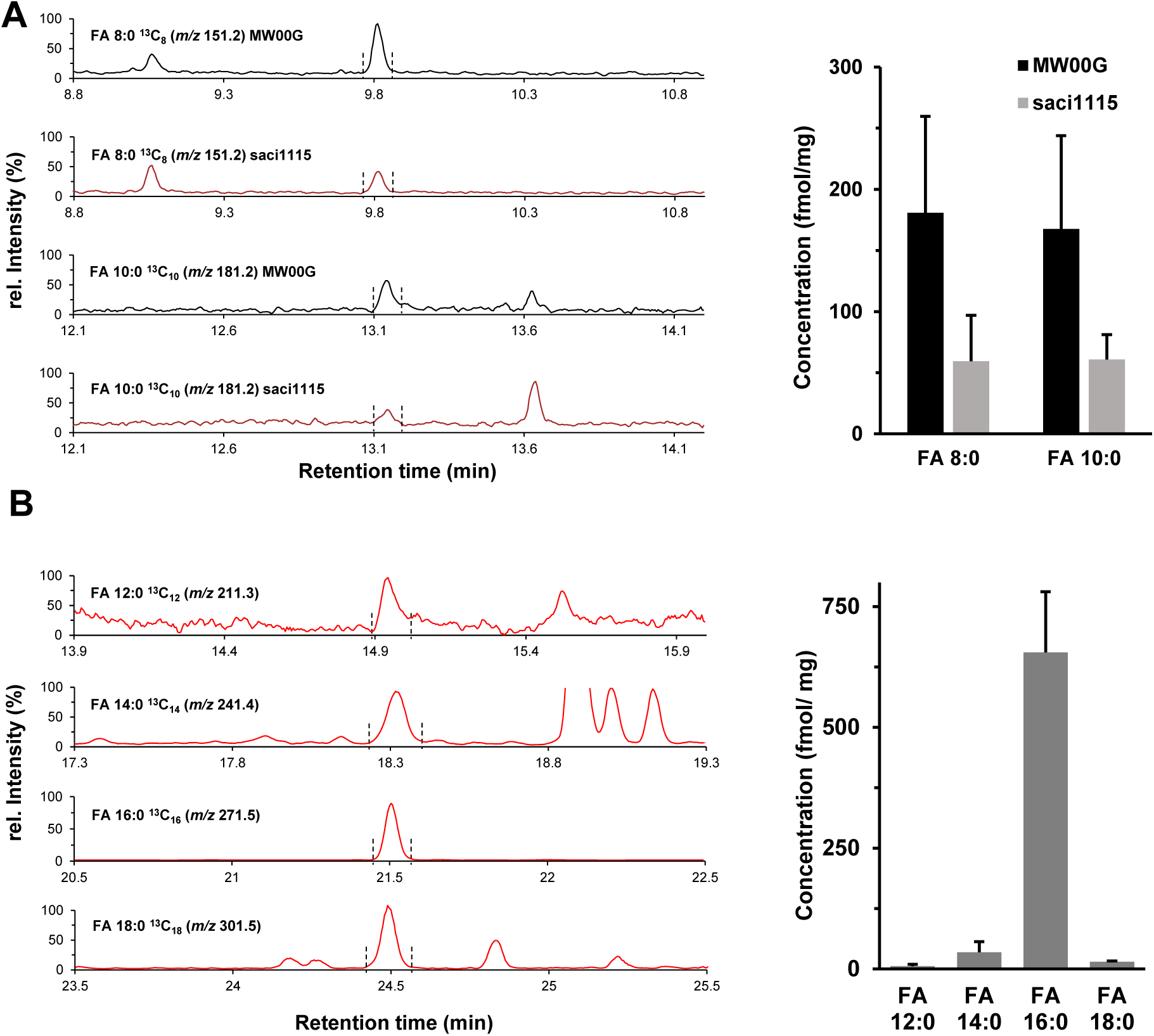
Fatty acid synthesis in *S. acidocaldarius* (MW00G parental strain and Δ*saci1115* deletion mutant) (A) and *H. volcanii* (B) as confirmed by isotopic labeling experiments and GC-MS. The *S. acidocaldarius* wildtype MW00G*, Δsaci1115* deletion mutant, and *H. volcanii* were grown on ^13^C-glycerol as only carbon source for four passages to obtain fully ^13^C labelled variants. Cells were isolated, hydrolysed, extracted and analysed for ^13^C labelled fatty acids as described in the methods section. (A) GC-MS analysis of *S. acidocaldarius* wildtype MW00G and *Δsaci1115* deletion mutant: Left panel shows the individual chromatograms for the mass-to-charge ratios (m/z) 151.2 for FA 8:0 and m/z 181.2 for FA 10:0 of the fully labelled ^13^C variants. For FA 8:0 a signal at the retention time of 9.8 min and for FA 10:0 at 13.2 min is observed (indicated by the dashed lines). All retention times were confirmed with ^12^C standards of the corresponding fatty acid. Right panel: Quantification of both fatty acids in *S. acidocaldarius* wildtype MW00G (black) and *Δsaci1115* deletion mutant (gray). (B) GC-MC analysis of *H. volcanii:* Left panel shows the individual chromatograms for the m/z that corresponds to the fully labelled FA 12:0 to FA 18:0. For FA 12:0 a signal at 15.0 min, for FA 14:0 at 18.3 min, for FA 16:0 at 21.5 min and for 18:0 at 24.5 min was observed. All retention times were confirmed by with ^12^C standards of the corresponding fatty acid. Right panel: Quantification of the individual fatty acids in *H. volcanii*.

### β oxidation enables fatty acid degradation in *S. acidocaldarius*

Previous studies have proposed that a reversible β oxidation pathway may potentially enable both fatty acid synthesis and degradation in Archaea. In *S. acidocaldarius,* the regulation of a β oxidation related gene cluster (*saci_1103-1126*) via a TetR like regulator (FadR) has been recently studied (*10*). To elucidate if the β oxidation enzymes encoded in the *saci_1103-1126* gene cluster in *S. acidocaldarius* function in fatty acid degradation and if they could additionally be involved in fatty acid synthesis, selected recombinant proteins were expressed, purified and characterized *in vitro* with respect to kinetic constants and chain length/substrate specificity (Table S1, Fig. S7-12). The conversions catalyzed by these enzymes were also confirmed via HPLC and the complete β oxidation cascade was reconstituted *in vitro* (Fig. 2; Fig. S13-19). Briefly, the acyl-CoA dehydrogenase (ACAD, Saci_1123) oxidized saturated acyl-CoAs to unsaturated 2,3-enoyl-CoAs, and used an unusual electron transfer flavoprotein ETF (Saci_0315, a fusion protein of the canonical α and β ETF subunits) as electron acceptor (Table S1, Fig. S7-9, Fig. S14, Fig. S18). The following conversions of enoyl-to the hydroxyacyl-CoA and further to the ketoacyl-CoA were catalysed by a bifunctional fusion protein (Saci_1109) (Table S1, Fig. S10-11, Fig. S15, Fig. S17-19) comprised of an N-terminal 3(S)-hydroxyacyl-CoA dehydrogenase (HCDH) and a C-terminal enoyl-CoA hydratase (ECH), which represents the inverted domain architecture of bacterial homologues (*12, 13*)(Fig. S10H). The ketoacyl-CoA was finally converted to acetyl-CoA by β-ketothiolase (KT) in presence of free CoA (Fig. 2, Fig. S12, Fig. S16-19).

**Fig. 2.**
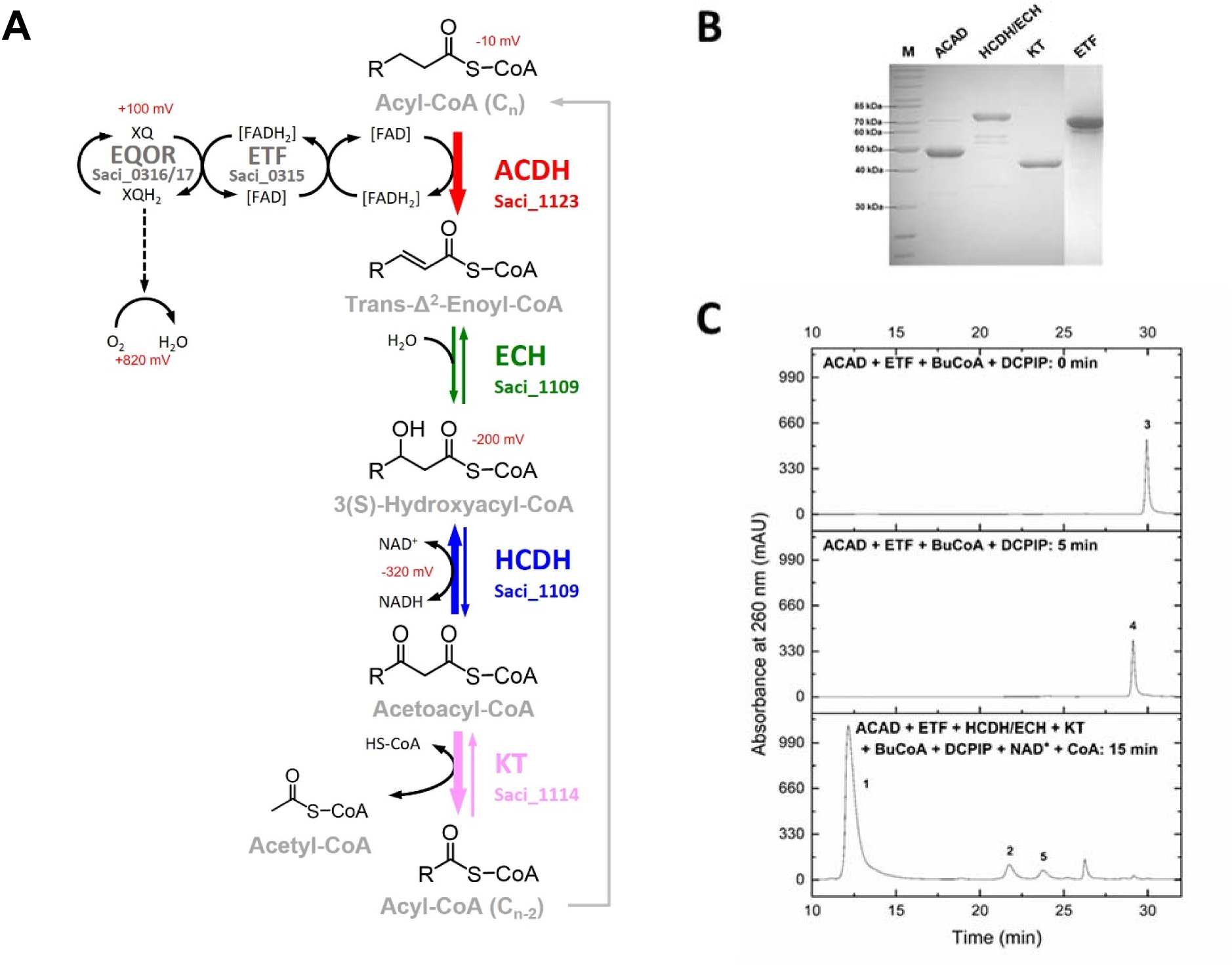
Reconstructed β oxidation pathway for fatty acid degradation in *S. acidocaldarius*. The β oxidation reactions (A), the purified recombinant proteins that catalyze these reactions (SDS-PAGE and Coomassie Blue staining) (B) as well as the HPLC chromatogram of the β oxidation enzyme cascade for butyryl-CoA conversion (C) are shown. In (A), the subscript “n” of acyl-CoA represents the number of carbon atoms in the fatty acid chain. The thickness of the arrow indicates the energetics of the respective reaction. For redox reactions the reduction potentials are given. The arrow colours (same as in Fig. 3) indicate the similar chemical conversion common to β oxidation and fatty acid synthesis. During the β oxidation enzyme cascade (C), butyryl-CoA (peak 3) was completely oxidized to crotonyl-CoA (peak 4) by ACAD transferring the electron to DCPIP through ETF in the first step. Then crotonyl-CoA was further converted to 3-hydroxybutyryl-CoA (peak 5) by HCDH/ECH and finally to acetyl-CoA (peak 2) by KT in presence of free CoA (peak 1). However, the intermediate acetoacetyl-CoA released by HCDH/ECH using NAD^+^ as cofactor was not detectable under the applied analytical conditions. The abbreviations: ACAD, acyl-CoA dehydrogenase; ETF, electron transfer flavoprotein; EQOR, ETF: quinone oxidoreductase; FAD, flavin adenine dinucleotide; NAD^+^, nicotinamide adenine dinucleotide; CoA, Coenzyme A; ECH, enoyl-CoA hydratase; HCDH, 3(S)-hydroxyacyl-CoA dehydrogenase; KT, β-ketothiolase or acetyl-CoA C acetyltransferase; BuCoA: butyryl-CoA; DCPIP: 2,6-dichlorophenolindophenol.

Together, these results show that *S. acidocaldarius* has a functional archaeal β oxidation pathway that supports fatty acid degradation. Importantly, our findings also suggest that this pathway likely does not operate in the reverse direction for mechanistic and thermodynamic reasons (see “Supplementary text”). For example, while the reaction catalyzed by ACAD (Saci_1123) ran to completion with both ETF and the artificial electron acceptor ferrocenium, it could not operate in the reductive direction (Fig. S14, Fig. S8). This is in line with the mechanisms described for ACAD enzymes which-in their reduced state-preferentially bind the product enoyl-CoA and thus kinetically promote the oxidative half-reaction, i.e. the electron transfer from the ACAD flavin to the electron acceptor (*14*). Furthermore, although the ETF (Saci_0315) could accept electrons from NADH (Fig. S7-8), it could not convey the electrons to re-reduce ACAD, further supporting that these enzymes cannot work in the reductive direction. Similarly, according to energetics, the reversal of the ECH/HCDH/KT (Saci_1109/Saci_1114) cascade is only possible to a low extent with a surplus of acetyl-CoA and NADH (Fig. S19). Furthermore, both ECH and HCDH parts of Saci_1109 were specific for the (S) stereoisomer characteristic for the β oxidation (Table S1, Fig. S10), again supporting that the reverse reaction is likely not supported by this protein. Together, these results demonstrate that *S. acidocaldarius* has a functional β oxidation pathway for fatty acid degradation that however does likely not support the reverse direction that would enable fatty acid synthesis.

### Unravelling a novel fatty acid synthesis pathway in *S. acidocaldarius*

Having ruled out that a reverse β oxidation pathway supports fatty acid synthesis in *S. acidocaldarius*, we explored alternative pathways that could support assembly of these molecules. In the absence of ACP and ACP synthase (*5, 8*), it is likely that such a pathway relies on CoA for fatty acid activation, as other CoA-dependent fatty acid conversions have been described (for example, from fatty acid elongation, from polyhydroxyalkanoate synthesis, from certain fermentations, and also from CO_2_ fixation pathways in Crenarchaeota and Thaumarchaeota). Another important question is if the pathway involves an initial reaction in which a KT homologue catalyzes the highly endergonic condensation of acyl-CoA substrates or whether this reaction is bypassed. Bypassing the reaction by using malonyl-CoA as an extender unit and coupling the Claisen condensation to decarboxylation appears unlikely, because although acetyl-CoA carboxylase is present in *S. acidocaldarius* (*saci_0260-0262*), no decarboxylating KTs could be identified and for the Saci_1114 KT malonyl-CoA dependent activity was disproved (see “Supplementary information”) (Fig. S21). Another possibility is to drive the KT-mediated reaction in the condensation direction by substrate channelling facilitated by complex formation, as recently described for isoprenoid synthesis in *Methanothermococcus thermolithotrophicus.* In this organisms, KT and the hydroxymethylglutaryl-CoA synthase that catalyses the subsequent reaction form a complex facilitated by a third protein with a DUF35 domain, with all three proteins being encoded in a single operon (*15*). Notably, in addition to the KT (Saci_1114) that is involved in β oxidation, the *saci_1103-1126* gene cluster encodes a second thiolase, Saci_1121, which forms an operon with a downstream gene encoding a DUF35 domain protein (Saci_1120), suggesting that these enzymes may form a complex involved in fatty acid synthesis. Indeed, homologous co-expression of both genes in *S. acidocaldarius* resulted in production of soluble and active protein whereas expression of the single proteins failed. Purification and size exclusion chromatography showed that both proteins form a stable complex (Fig. S22). Of note, the catalytic efficiency of the reversibly operating Saci_1121/1120 complex in the direction of acetoac(et)yl-CoA formation was 60-fold higher (the *K_M_* for acetyl-CoA 150-fold lower) than for the stand-alone KT Saci_1114 (see above) whereas the efficiency in the cleavage direction was similar (Fig. S23, Fig. S12, Table S1 and S2).

The next steps in fatty acid synthesis are the reduction of the ketoacyl-CoA followed by dehydration of the hydroxyacyl-CoA to the enoyl-CoA. The *saci_1103-1126* gene cluster includes one gene, *saci_1104,* annotated as *fabG,* a homologue of the canonical ketoacyl-ACP reductases belonging to the short chain dehydrogenase/reductase (SDR) superfamily (*16*) of which in total eleven representatives are present in the *S. acidocaldarius* genome. We expressed and purified the encoded enzyme, which was able to catalyse the reversible ketoacyl-CoA reduction with a strict specificity for NADPH and (R)-hydroxyacyl-CoA stereoisomer and a chain length preference for C8 over C4 (Table S2, Fig: S24-S25). Thus, this aceto(keto)acyl-CoA reductase (ACR, Saci_1104) has the opposite cosubstrate and stereospecificity compared to the bifunctional fusion protein Saci_1109 that is involved in β oxidation (see above), explaining how fatty acid synthesis and degradation are kept separate in *S. acidocaldarus*.

The presence of an NADPH dependent ACR with (R) specificity raised the questions whether *S. acidocaldarius* also harbor an (R)-specific hydroxyacyl thioester dehydratase. In Bacteria and Eukarya this reaction is carried out by dehydratases from the hotdog fold superfamily (*16*), which were not present in archaeal genomes (*17*) including *S. acidocaldarius*. Only distantly related, putative MaoC-like hydroxyacyl-CoA dehydratases (MaoC-HCD), which also belong to the hotdog fold superfamily, were identified, particularly *saci_1070* and *saci_1085*. Such MaoC-like dehydratases are known from some fatty acid related pathways in Bacteria, such as in mycobacteria (*18*). We expressed and purified these proteins and Saci_1085 indeed showed reversible 3-hydroxyacyl-CoA dehydratase (HCD) activity forming 2,3-enoyl-CoAs with a pronounced preference for the (R)-stereoisomer and preference for C8 enoyl-CoA over C4 and C6 (Table S2, Fig. S25, Fig. S26). Thus, both the ketoacyl-CoA reduction and dehydration reactions allow for the separation of fatty acid synthesis from degradation in *S. acidocaldarius*.

The final step in fatty acid synthesis is the enoyl-CoA reduction yielding the saturated acyl-CoA, which in the canonical pathways usually depends on NADPH as electron donor rendering this reaction highly exergonic (ΔG^0’^-56.2 kJ mol^-1^) (*16, 19*). The canonical ACP-dependent enoyl thioester reductases belong either to the SDR (FAS II Bacteria, plants, protists) or to the medium chain dehydrogenase/reductase (MDR) superfamily (mitochondrial FAS II, mammalian FAS I) (*19*). In addition to the ACR Saci_1104 (see above), one further gene (*saci_1115*) in the *saci_1103-1126* gene cluster is annotated as a MDR dehydrogenase, suggesting that the enzyme encoded by this gene is a potential candidate to catalyse this reaction. Therefore, we expressed and purified Saci_1115 and demonstrate that the protein is indeed an enoyl-CoA reductase (ECR) capable of catalysing the NADPH specific reduction of enoyl-CoAs of various chain lengths from C4-C10 (C8 preferred), with some minor residual activity also with C16 (Table S2, Fig. S27-S28).

Thus, all four enzymes required for fatty acid synthesis were identified in *S. acidocaldarius*. To demonstrate the functionality of this pathway, the whole cascade comprising the purified Saci_1121/1120 KT/DUF35 complex, the Saci_1104 KCR, the Saci_1085 MaoC-HCD, and the Saci_1115 ECR was stepwise reconstituted *in vitro* and the respective conversions were observed via HPLC (Fig. 3, Fig. S29, Fig. S26). First, we confirmed the conversion of 3(R)-hydroxybutyryl-CoA and acetoacetyl-CoA to butyryl-CoA via MaoC-HCDand ECR as well as ACR, MaoC-HCD, and ECR, respectively (Fig. S26, Fig. S29). Then, all four enzymes were combined in equal activities *in vitro* with acetyl-CoA and NADPH, resulting in the detection of increasing amounts of hexanoyl-CoA within a time frame from 0 h to 21 h (Fig. 3). The rather slow formation of butyryl-CoA and hexanoyl-CoA can likely be attributed to the comparably low catalytic efficiency of the ECR and feedback inhibition by product and other pathway intermediates (*20*). Interestingly, other pathway intermediates were hardly detectable via HPLC, which may be explained by substrate channelling. Collectively, these data delineate a novel pathway capable of supporting fatty acid synthesis in archaea.

**Fig. 3.**
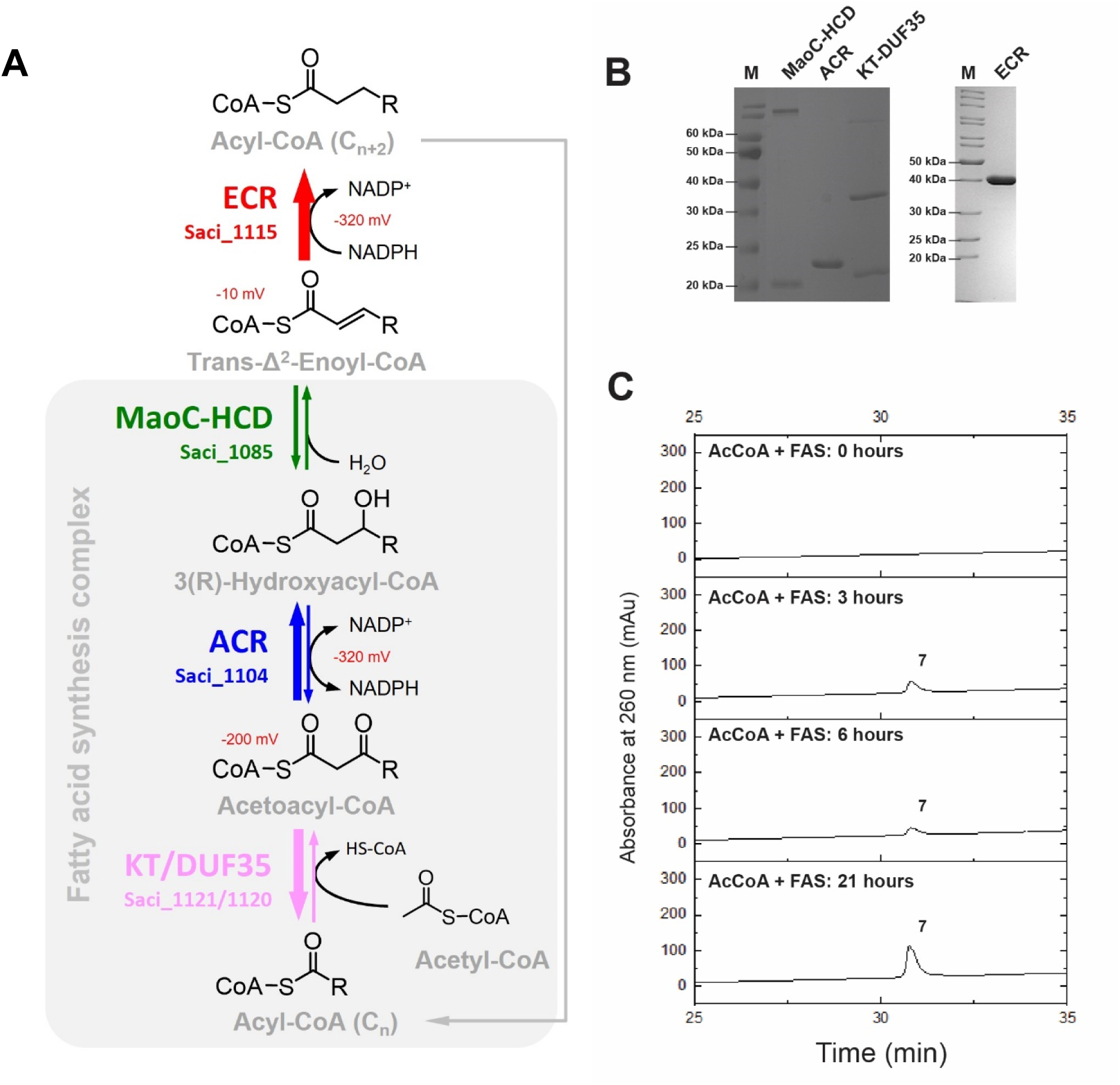
Reconstructed novel pathway for fatty acid synthesis in *S. acidocaldarius*. The fatty acid synthesis reactions (A), the purified recombinant proteins that catalyze these reactions (SDS-PAGE and Coomassie Blue staining) (B) as well as the HPLC chromatogram of the fatty acid synthesis enzyme cascade for acetyl-CoA conversion to hexanoyl-CoA (C) are shown. In (A), the subscript “n” of acyl-CoA represents the number of carbon atoms in the fatty acid chain. The thickness of the arrows indicates the energetics of the respective reaction. For redox reactions the reduction potentials are given. The arrow colours (same as in Fig. 3) indicate the similar chemical conversion common to β oxidation and fatty acid synthesis. Gray shading indicates the complex formation between KT/DUF35, ACR, and MaoC-HCD likely driving especially the endergonic Claisen condensation (for details see text and Fig. 4). The abbreviations: KT, β-ketothiolase; DUF35, domain of unknown function 35 protein; KT/DUF35, subcomplex composed of KT and DUF35; ACR, aceto(keto)acyl-CoA reductase; MaoC-HCD, MaoC-like (R)-hydroxyacyl-CoA dehydratase; ECR, enoyl-CoA reductase; AcCoA, acetyl-CoA; NADP^+^, nicotinamide adenine dinucleotide phosphate; CoA, Coenzyme A; FAS, fatty acid synthesis enzyme cascade comprised of KT/DUF35, ACR, MaoC-HCD, ECR).

To further support the *in vivo* significance of this fatty acid synthesis pathway, we generated an in-frame deletion mutant of the ECR-encoding gene *saci_1115* in *S. acidocaldarius* and analyzed the growth phenotype of the mutant strain and its fatty acid content. Whereas the growth of the Δ*saci_1115* mutant was unaffected (Fig. S30), its content in C8:0 and C10:0 fatty acids was substantially reduced (by 60%) upon growth on ^13^C glycerol (Fig. 1A), supporting that the proposed pathway is involved in fatty acid synthesis in *S. acidocaldarius in vivo*. However, the residual fatty acid content detected in the Δ*saci_1115* mutant suggests that another ECR is also capable of supporting fatty acid synthesis.

### KT, ACR and HCD form a stable fatty acid synthase complex mediated by a DUF35 domain protein

As described above, in isoprenoid synthesis KT/DUF35 forms a complex with the next enzyme in the pathway, HMG synthase. Therefore, we tested whether the KT/DUF35 pair (Saci_1121/Saci_1120) involved in fatty acid synthesis also forms a similar complex with other enzymes in the pathway. For this, we performed *in vitro* reconstitution and size exclusion chromatography (SEC) experiments with his-tagged Saci_1104 ACR, twin-strep-tagged Saci_1121/20 KT/DUF35 complex and twin-strep-tagged Saci_1085 (MaoC-HCD). Upon *in vitro* reconstitution, all four proteins Saci_1121/1120/1104/1085 ran in a single peak at 171 kDa (Fig. 4A). The Saci_1121/1120 complex alone eluted at 90 kDa (between a αβ heterodimer and a α2β2 heterotetramer), while native mass spectrometry (nMS) experiments revealed a molecular mass of 130 kDa supporting the α2β2 heterotetrameric structure (Fig. S31). Each, the Saci_1121/1120/1104 (Fig. 4B) and the Saci_1121/1120/1085 (Fig. 4C) mixtures also showed a single peak with molecular masses of approximately 155 kDa in SEC indicating that both Saci_1104 and Saci_1085 directly interact with the Saci_1121/1120 heterodimer, which would fit to either α2β2γ and α2β2δ or α2β2γ2 and α2β2δ2 heterotrimers (due to inaccuracy of the SEC). Binding of Saci_1115 ECR to the complex could not be demonstrated (Fig. 4D).

**Fig. 4.**
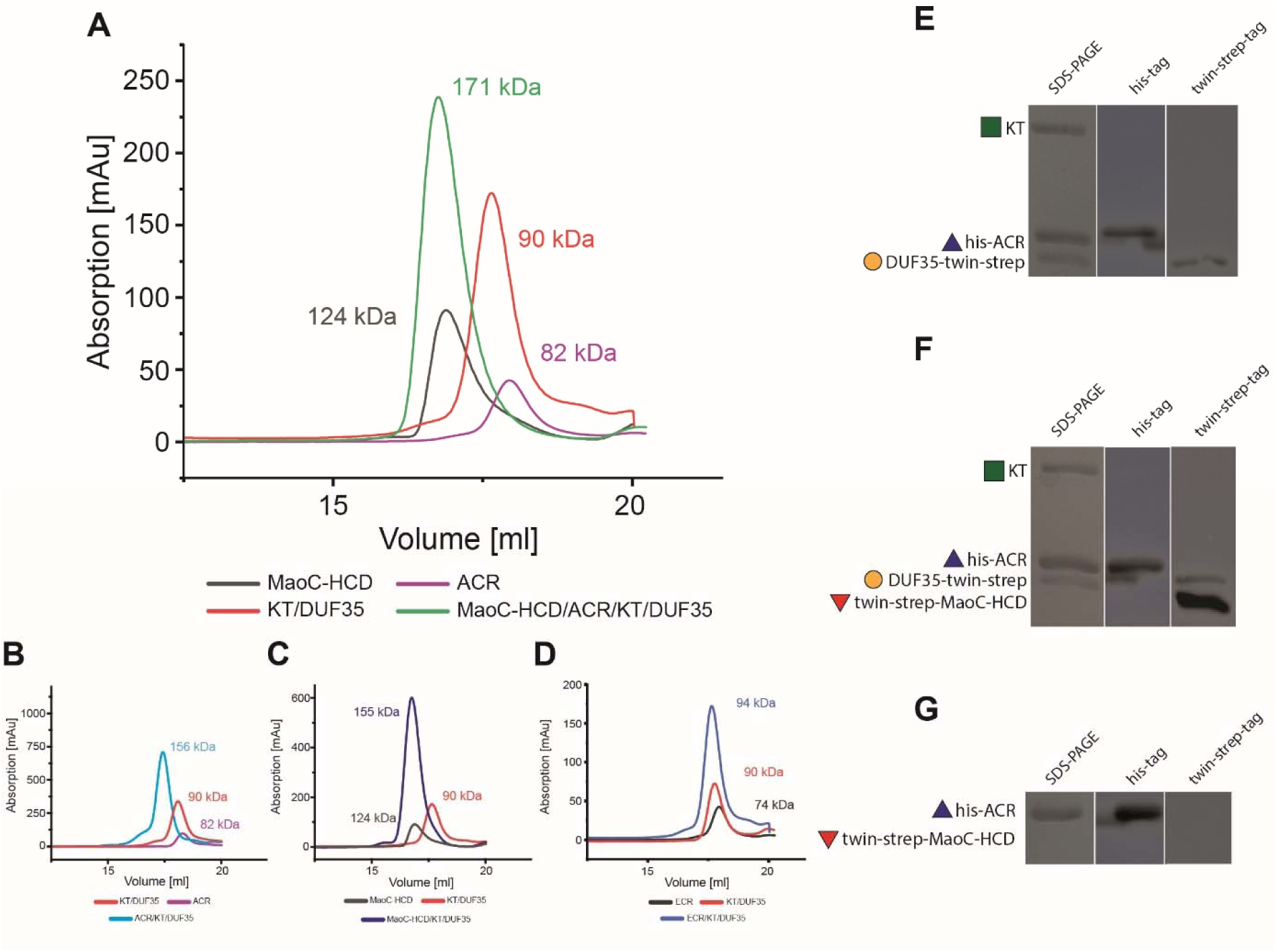
Fatty acid synthesis enzymes from *S. acidocaldarius* form a stable complex mediated by a DUF35 protein. (A) Size exclusion chromatography of the Saci_1121/1120 KT/DUF35 complex, the Saci_1104 ACR, and the Saci_1085 MaoC-HCD alone as well as of all four proteins reconstituted *in vitro* (the native molecular masses derived from calibration are indicated). (B) and (C) show the subcomplex formation with a shift to higher molecular masses of Saci_1121/1120 KT/DUF35 in presence of Saci_1104 ACR and Saci_1085 MaoC-HCD, respectively. (D) Saci_1115 ECR does not interact with the Saci_1121/1120 KT/DUF35 subcomplex. For comparison, the elution profiles of the single proteins are shown in each panel (B-D). (E-G) Affinity chromatography based coelution experiments analyzed by SDS-PAGE and Western blot analyses. In (E), N-terminally his-tagged ACR Saci_1104 (his-ACR) is bound to Ni-NTA and the KT/DUF35 (Saci_1121/Saci_1120) subcomplex C-terminally twin-strep-tagged on DUF35 (Saci_1120) (DUF35-twin-strep) was applied, after elution one his-tagged protein (his-ACR, Saci_1104), one twin-strep-tagged protein (DUF35-twin-strep, Saci_1120), and un-tagged KT Saci_1121 (45 kDa, SDS-PAGE) are observed indicating protein-protein interaction. In (F), his-ACR was first bound to the Ni-NTA column followed by applying KT/DUF35-twin-strep (Saci_1121/Saci_1120) subcomplex and the N-terminally twin-strep-tagged MaoC-HCD (twin-strep-MaoC-HCD). After washing, all four proteins coeluted as confirmed by the presence of his-ACR, DUF35-twin-strep, twin-strep-MaoC-HCD, and the un-tagged Saci_1121 KT (45 kDa, SDS-PAGE). This coelution confirms protein interaction and thus stable complex formation of all four proteins. (G) shows that twin-strep-MaoC-HCD (Saci_1085) does not bind to the Ni-NTA column bound his-ACR (Saci_1104) as indicated by the missing detection with the anti-strep antibody, and suggesting a central role of the KT/DUF35 Saci_1121/1120 assembly in complex formation.

We further confirmed the interactions using coelution experiments in which the N-terminally his-tagged Saci_1104 was bound to a Ni-NTA column. Then Saci_1121/1120 complex C-terminally twin-strep-tagged on Saci_1120 and/or N-terminally twin-strep-tagged Saci_1085 were consecutively applied, which could only bind to the column when interacting with Saci_1104. In case of interaction with Saci_1104, coelution of the proteins should occur with imidazole. Indeed, all four proteins coeluted as indicated by SDS-PAGE and Western blot detection (Fig. 4E). Also, Saci_1104 and Saci_1121/1120 (in the absence of Saci_1085) (Fig. 4F) coeluted, whereas Saci_1104 and Saci_1085 did not show an interaction (Fig. 4G). This confirmed the results from SEC and suggests that both Saci_1104 and Saci_1085 interact with the Saci_1121/1120 heterodimer but not with each other.

Collectively, these results strongly indicate that three enzymes (KT, ACR and MaoC--HCD) catalyzing ac(et)yl-CoA to enoyl-CoA_(Cn+2)_ conversion in fatty acid synthesis of *S. acidocaldarius* form a fatty acid synthase complex. This complex, whose formation is at least partly mediated by a DUF35 domain protein, drives the initial Claisen condensation reaction but also the further conversions to the respective enoyl-CoAs, likely by facilitating substrate channelling from KT to the ACR and MaoC-HCD.

### Fatty acid biosynthesis pathways are present in diverse archaeal lineages

The identification of a novel pathway that enables *de novo* fatty acid synthesis in *S. acidocaldarius* prompted us to explore whether similar pathways are present in other Archaea. For this, we analyzed the distribution and phylogenetic relationships of the genes encoding the fatty acid synthesis pathway in *S. acidocaldarius* across archaeal genomes. This analysis showed that homologues of the *S. acidocaldarius* fatty acid synthesis genes are commonly found in monophyletic groups with other Crenarchaeota (Fig. 5, Fig. S32-S36), often sister to Thaumarchaeota and Thermoplasmatota sequences. These archaeal clades are commonly found as sister groups to bacterial groups like Actinobacteria, Chloroflexi, Firmicutes and Proteobacteria, suggesting frequent horizontal gene transfer (HGT) events (Fig. 5A). In other archaea, such as Thermococcales or Halobacteriales (although we clearly demonstrated fatty acids in *H. volcanii*), we could not identify closely related homologues of the proposed fatty acid synthesis pathway identified in *S. acidocaldarius*. However, the proteins involved in this pathway belong to large superfamilies with dynamic evolutionary histories marked by multiple HGT events, which complicates functional assignment based solely on sequence similarities. Therefore, similar functions might be occupied by distant homologues in archaea which might be dynamically replaced in genomes of different clades indicating a versatile fatty acid synthesis pathway, reminiscent of the diversity in sugar degradation routes in Archaea (*21*).

**Fig. 5.**
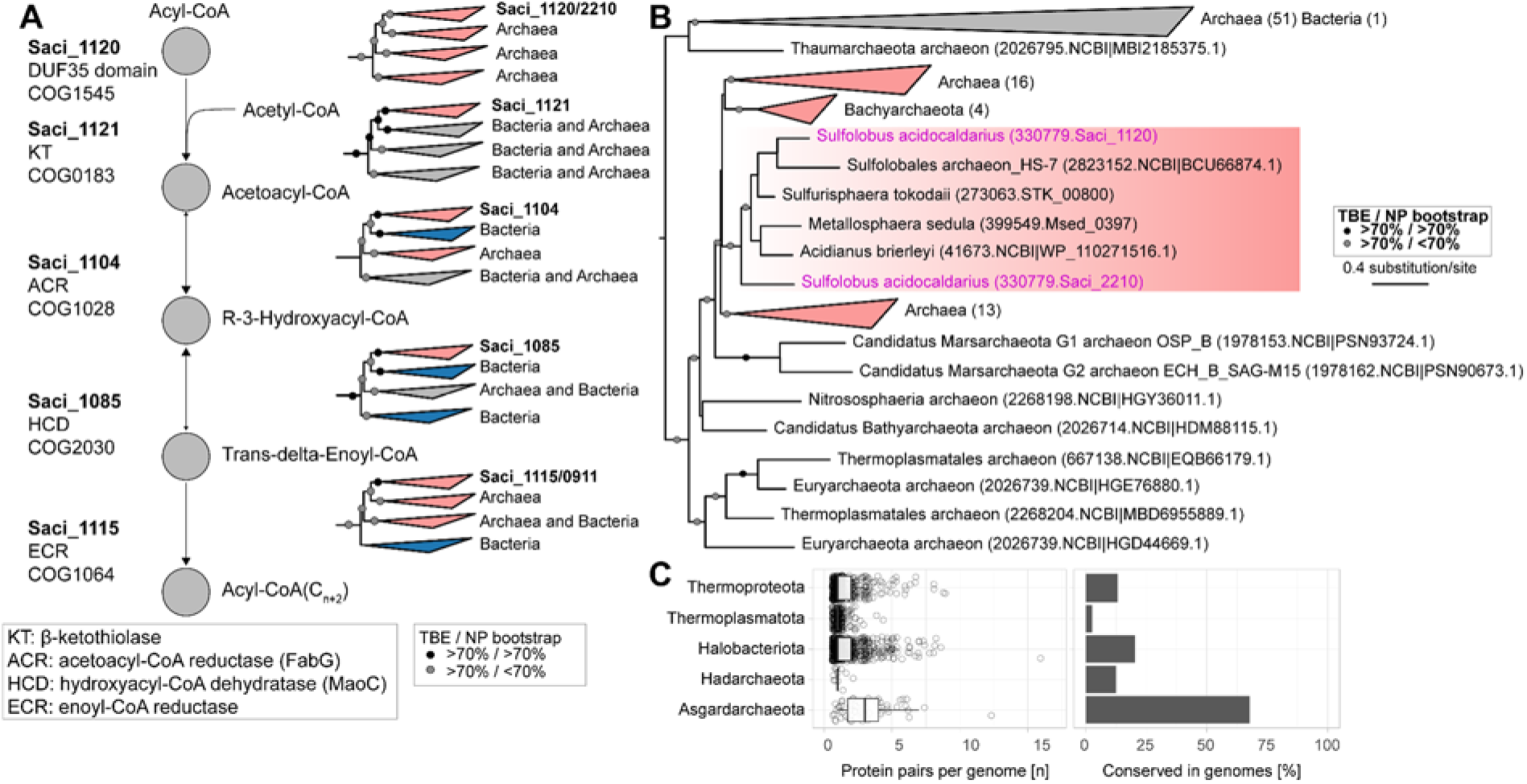
Phylogenies of the *S. acidocaldarius* proteins involved in fatty acid synthesis. (A) Simplified representations of maximum likelihood (ML) phylogenies for proteins proposed to be involved in fatty acid synthesis in *S. acidocaldarius*. The workflow and detailed phylogenetic trees are available in Fig. S32-36. (B) Maximum likelihood phylogenetic tree displaying the archaeal heritage of the DUF35 domain protein Saci_1120. Phylogenetic trees were rooted according to the initial COG gene tree sister clades. *S. acidocaldarius* proteins are highlighted in pink. Transfer bootstrap expectation, TBE; Non-parametric bootstrap, NP bootstrap. (C) Number of ketothiolase/DUF35 pairs per genome of phyla that contain more than one copy and per phylum-rank conservation of more than one pair per genome.

While homologues of most of the fatty acid synthesis genes identified in *S. acidocaldarius* including DUF35 domain proteins seem to be present in both Bacteria and Archaea, homologues of the specific DUF35 protein Saci_1120 seem to be restricted to archaea (Fig. 5B). Furthermore, genes encoding KT/DUF35 protein pairs are highly conserved genomic neighbors across various archaeal phyla, where KT/DUF35 form a complex with HMG-CoA synthase involved in isoprenoid and membrane lipid biosynthesis (*15*). However, we respectively find two and seven KT/DUF35 domain protein pairs in the genomes of *H. volcanii* and *S. acidocaldarius,* suggesting that different pairs may be involved in the metabolism of different substrates. Notably, we observe the occurrence of multiple KT/DUF35 protein pairs per genome in several archaeal phyla (Fig. 5C), most prominently conserved in 67% of screened Asgardarchaeota genomes (n=62), which carry an average of 3.8 pairs per genome. While additional experiments are needed to confirm that these genes identified in other Archaea comprise functional pathways, these observations suggest that the ability to synthesize fatty acids may be widespread in Archaea.

## Discussion

Our study represents a comprehensive analysis of fatty acid metabolism in Archaea. First, using ^13^C labelling experiments with glycerol as a carbon source, we unequivocally show that diverse Archaea contain fatty acids and are able to synthesize these compounds *de novo*, as exemplified by the model organisms *S. acidocaldarius* and *H. volcanii* (Fig. 1). In *S. acidocaldarius,* no fatty acids were detected when cells were grown on ^13^C-labelled D-xylose, indicating that fatty acid formation is dependent on the growth substrate (Fig. S6). Notably, our finding that *H. volcanii* primarily contains palmitic acid (C16:0) is in agreement with previous reports that also found this fatty acid bound to the chloride pump halorhodopsin from *Halobacterium salinarum* (*22*). By contrast, we found that the most abundant species in *S. acidocaldarius* are C8:0 and C10:0, whereas in *Saccharolobus solfataricus*, a close relative of *S. acidocaldarius,* C16:0/C16:1 and C18:0/C18:1 fatty acids were previously reported (*7*). Furthermore, the fatty acid content observed for *S. acidocaldarius* was in the pmol mg^-1^ dry weight range, whereas previous studies have reported abundances in the nmols mg^-1^ cell dry weight for other Archaea (*23, 24*), which might be due to the use different methodologies and/or different medium composition. Nevertheless, our labelling experiments exclude the possibility that the detected fatty acids are contaminants and thus demonstrate *de novo* synthesis of fatty acids in organisms from distant archaeal phyla.

Then, using biochemical approaches, we show that the β oxidation homologues present in *S. acidocaldarius* function in fatty acid degradation, in a pathway that involves the sequential activities of ACAD (Saci_1123), an ECH/HCDH fusion protein (Saci_1109) and KT (Saci_1114) (Fig. 2). Importantly, our detailed characterization of these enzymes also supports that several mechanistic, specificity and thermodynamic constraints prevent this pathway from functioning in the reverse direction to support fatty acid synthesis (see “Supplementary text”). These constraints likely also apply for other Archaea harbouring complete sets of β oxidation related enzymes, which have been indicated to be of bacterial origin (*5*).

We then identified a previously uncharacterized, ACP-independent, fatty acid synthesis pathway in *S. acidocaldarius* that meets the constraints of thermodynamics and pathway separation from β oxidation (Fig. 3 and “Supplementary text”). Furthermore, we show that three of the enzymes in this pathway – the Saci_1121 KT, the Saci_1104 ACR, and the Saci_1085 MaoC-HCD– form a stable assembly mediated by the DUF35 domain protein Saci_1120, which represents an archaeal fatty acid synthase complex in *S. acidocaldarius* (Fig. 4).

We further show that homologues of the genes involved in fatty acid synthesis are present across various archaeal lineages including Crenarchaeeota, Thermoplasmatota, and Thaumarchaeota. In combination with the demonstrated presence of fatty acids in the crenarchaeon *S. acidocaldarius* and the euryarchaeon *H. volcanii* this suggests that the capacity to synthesize fatty acids is likely widespread across the archaeal domain (Fig. 5, Fig. S32-S36). We find that several archaea carry multiple KT/DUF35 domain protein pairs, which may represent metabolic adaptations to various substrates. Notably, these pairs are particularly abundant and conserved in Asgardarchaeota genomes, from which eukaryotes are thought to have emerged, which supports previous suggestions that organisms in this clade may be capable of synthesizing fatty acids. Therefore, our study represents a substantial advance to clarify the long-lasting debate around the presence and metabolism of fatty acids in Archaea and provides further insights to resolve the conundrum of the “lipid divide”.

Finally, the demonstration that Archaea can synthesize fatty acids raises questions about the function of these molecules in these cells. While previous studies have suggested that fatty acids are integrated into membrane lipids (*23*), we were unable to specifically assess the localization of these molecules given the low abundance of fatty acids detected in *S. acidocaldarius.* However, this low fatty acid content, together with the observation that fatty acid synthesis is dependent on the growth conditions, suggests that fatty acids have more restricted, specialized functions. As fatty acids and other fatty acids-based lipids have been found in energy transducing membrane protein complexes from some eukaryotes and cyanobacteria (such as cytochrome bc1 and cytochrome C oxidases), as well as in halorhodopsin from *H. salinarum*, this suggests that fatty acid synthesis in Archaea may have been required for the evolutionary acquisition and functioning of such protein complexes. This is supported by the presence of homologues of these energy transducing membrane protein complexes in both *S. acidocaldarius* and *H. volcanii* (*5, 22*). Furthermore, fatty acids are precursors for the biosynthesis of the coenzymes biotin and lipoic acids, which are essential to the activity of enzyme like carboxylases and the glycine cleavage system that are also widespread in Archaea (*25–28*). Future studies are needed to clarify the importance of fatty acid synthesis for these processes and to explore potential additional roles including in cell signalling or molecule targeting.

## Methods

### Reagents

The CoA esters including CoA, acetyl-CoA and butyryl-CoA employed in this study were purchased from Millipore Sigma® Merck KGaA in Darmstadt, Germany. The commercial hexanoyl-CoA, octanoyl-CoA, crotonyl-CoA and mixed 3(S/R)-hydroxybutyryl-CoA were produced by Santa Cruz Biotechnology (Dallas, USA). The decenoyl-CoA, hexadecenoyl-CoA as well as the individual 3(S)-hydroxybutyryl-CoA and 3(R)-hydroxybutyryl-CoA were chemically synthesized as described before (*29*). The compound acetoacetyl-CoA was purchased either from Millipore Sigma® Merck KGaA or via chemical synthesis. The substrates hexenoyl-CoA and octenoyl-CoA were enzymatically synthesized as described in the assay for ECR (Saci_1115). All other chemicals and reagents were obtained from Sigma (Germany), Thermo Fisher Scientific (USA), Carl Roth (Germany), Abcam (United Kingdom), IBA (Germany), Sartorius (Germany), Cytavia (USA).

### Strains and growth conditions

The *E. coli* strains DH5α for cloning and Rosetta (DE3) for expression were cultivated in lysogeny broth (LB) containing appropriate antibiotics (100 μg/ml ampicillin for strains containing plasmid pET15b, pET45b, pBS-araFX-UTR-CtSS or pBS-araFX-UTR-NtSS (*30*), 50 μg/ml kanamycin for strains including plasmid pET28b as well as for *E. coli* ER1821 and 50 μg/ml chloramphenicol for Rosetta (DE3), respectively) at 37°C under aerobic conditions (all strains used in this work are listed in Table S3).

*S. acidocaldarius* strains MW001 (uracil auxotrophic) (*31*) and MW001 adapted to growth on glycerol designated herein MW00G (*32*) were cultured aerobically (Innova®44, New Brunswick, Germany) in Brock′s basal medium at 75°C, pH 3, 180 rpm (*33*) supplemented with 10 µg/ml of uracil as well as 0.1% (w/v) N-Z-amine and 0.3% (w/v) D-xylose (for induction of homologous overexpression). The Gelrite solidified Brock medium used for *S. acidocaldarius* MW001 and MW00G based cloning and transformation procedures was prepared as mentioned before (*31*).

For preparation of cell samples for fatty acid analyses a *S. acidocaldarius* MW00G was used. This strain also served as parental strain for construction of the Δ1115 deletion mutant (see below). For isotopic labelling *S. acidocaldarius* MW00G and the Δ1115 deletion mutant were pre-grown on Brock′s basal medium (see above) with ^13^G-glycerol as sole carbon source for four passages. With the fourth passage 500 mL cultures (triplicates) were inoculated to an initial OD_600_ 0.05 and grown until an OD_600_ of 0.7. Cells were cooled down using liquid nitrogen and harvest by centrifugation at 4°C, 7000 rpm for 15 min. Cell pellets were stored at-70°C until use.

*Haloferax volcanii* H26 cells were essentially cultivated in salt water minimal medium with 10 mM glycerol as sole source of carbon and energy (*34*). Isotopic labelling with 10 mM ^13^G-glycerol was performed as described for *S. acidocaldarius* MW00G. Cell pellets were stored at-70°C until use.

### Cloning, expression and purification of the recombinant proteins

Open reading frames (ORFs) *saci_1123*, *saci_0315, saci_1109*, *saci_1114* as well as *saci_1104*, *saci_1115, saci_1085, and* for coexpression *saci_1121/saci_1120* (from start codon *saci_1121* to stop of *saci_1120*) were PCR amplified using genomic DNA of *S. acidocaldarius* DSM639 as template (the employed primer pairs are listed in Table S3) and cloned into respective plasmid vectors (Table S3). For *saci_1114*, a codon optimized version cloned into the pET28b vector for expression in *E. coli* was purchased from the GeneArt synthesis service (Thermo Fisher Scientific, USA). The recombinant plasmids were transformed into *E. coli* DH5α used for propagation and into *E. coli* strain Rosetta (DE3) for heterologous overproduction induced by isopropyl-β-d-thiogalactopyranoside (IPTG) or into *S. acidocaldarius* MW001 for homologous expression induced by D-xylose (see above) (for details see Table S3-S4).

For expression cell were grown as described above. The cell pellets were harvested by centrifugation at 4°C, 7000 rpm for 15 min. Except for the cells maintaining ACAD (*saci_1123*) or ETF (*saci_0315*) protein (see below), cell pellets were resuspended in 50 mM NaH_2_PO_4_ (pH 8) containing 300 mM NaCl in a ratio of 3 ml buffer per 1 g wet cell weight (nickel ion affinity purification) or in 100 mM TRIS-HCl (pH 8.0), 150 mM NaCl, 1 mM EDTA in a ratio of 10 ml buffer per1 g wet cell weight (strep-tactin affinity purification). Cell suspensions were lysed either by sonication (0.5 cycle, 55 amplitude for 15 min) or via French press (3 times, 1200 psi). After centrifugation (4°C, 15000 rpm, 45 min) the supernatants were subjected to heat precipitation at 70°C in a water bath for 20 min (except for Saci_1085 and Saci_1121/Saci_1120) to separate thermostable proteins of interest from precipitated host proteins via centrifugation (4°C, 15000 rpm for 30 min). Afterwards, the his-tagged proteins were purified from the supernatant via Protino^®^ Ni-TED (tris-carboxymethyl ethylene diamine) affinity chromatography (Machery & Nagel, Düren, Germany) whereas the twin-strep-tagged Saci_1085 and twin-strep-tagged Saci_1121/Saci_1120 were purified by Strep-Tactin^®^XT system (IBA Solutions for Life Science, Göttingen Germany) according to the manufacturer’s instructions. An extra wash step was performed for Saci_1085 using 500 mM NaCl and 0.1% (w/v) SDS to improve removal of protein impurities. The protein samples were then applied to size exclusion chromatography (Superdex 200 prep grad, HiLoad 26/60 or 16/60, GE Healthcare Life Sciences, Freiburg, Germany) using 50 mM HEPES-KOH (pH 7.2), containing 300 mM NaCl for equilibration and elution of the proteins Saci_1109, Saci_1104 and Saci_1115, respectively. For Saci_1114, 50 mM HEPES-KOH (pH 7.2), 300 mM KCl and 3 mM DTT was used. Finally, all purified proteins were stored in 25% (v/v) glycerol at-70°C for further use. The pure Saci_1114, Saci_1085, and Saci_1121/Saci_1120 proteins were flash frozen with liquid nitrogen prior to long-term storage.

The ACAD (Saci_1123) and ETF (Saci_0315) containing cells were resuspended in 50 mM TRIS-HCl (pH 7.5) containing 150 mM KCl, 10 mM imidazole and 0.1% (v/v) Triton X-100 and were disrupted by sonication (see above). The cell debris was removed by centrifugation (4°C, 15000 rpm, 40 min) and the supernatant was supplemented with 1 mM FAD to improve the cofactor binding. Purification of ACAD and ETF proceeded *via* metal-ion affinity chromatography using an Äkta purifier (GE Healthcare) system. The crude extracts were filtered (0.45 µM polyvinylidene fluoride membrane, Carl Roth, Karlsruhe, Germany) and applied to 1 or 5 ml Nickel-IDA-Sepharose column according to the manufacturer’s instructions. The column was washed with 50 mM TRIS-HCl (pH 7.5) containing 150 mM KCl and 10 mM imidazole, and proteins were eluted with the same buffer supplemented with 400 mM imidazole. After elution, imidazole was removed using a 10 or 30 kDa cut-off centrifugal concentrator (Sartorius, Goettingen, Germany). The concentrated pure Saci_1123 and Saci_0315 proteins were stored in 25 mM PIPES-HCl (pH 6.5), 20 mM KCl, 10% (v/v) glycerol followed by flash freezing in liquid nitrogen and stored at-70°C.

The preparative SEC runs were also used to determine the native molecular masses of the single enzymes (if not stated otherwise, see below) using the Gel Filtration LMW Calibration Kit and Gel Filtration HMW Calibration Kit (Cytavia, USA) Protein concentrations were determined with TECAN reader at 450 nm and 595 nm according to the standard Bradford assay (*35*). Purification and molecular weight were visualized by SDS-polyacrylamide gel electrophoresis and the Coomassie Brilliant Blue staining. The purified proteins were employed for further enzyme characterization.

### Analysis of protein interaction between HCDH/ECH Saci_1109 and KT Saci_1114

To study the complex formation between the recombinant HCDH/ECH and KT proteins, 1090 µg of purified recombinant ECH/HCDH and 644 µg of purified recombinant KT were mixed and incubated on ice for 4 hours. Afterwards, the protein mixture was applied to the Superdex 200 prep grad HiLoad 16/60 size exclusion chromatography column using 50 mM HEPES-NaOH (pH 7.2), 300 mM NaCl for equilibration and elution.

#### Determination of the native molecular masses and analyses of complex formation of proteins involved in fatty acid synthesis

To determine the native molecular masses of the coexpressed protein complex Saci_1121/Saci_1120 and the single proteins Saci_1104, Saci_1085 and Saci_1115 size exclusion chromatography was used. To this end, pooled enzyme samples after affinity chromatography were concentrated using centrifugal concentrators (10 kDa cutoff) (Sartorius, Germany) and applied to a Superose 6/100 column (Cytavia, USA) using 50 mM TRIS-HCl (pH 8) containing 250 mM NaCl as buffer for equilibration and elution. In total 500 µl of protein samples were loaded corresponding to 300 µg of C-terminally twin-strep-tagged Saci_1121/Saci_1120, 150 µg of N-terminally his-tagged Saci_1104, 250 µg N-terminally twin-strep-tagged Saci_1085, and 400 µg of N-terminally his-tagged Saci_1115. For analyses of protein complex formation 600 µg of Saci_1121/Saci_1120 was reconstituted in the indicated combinations with 200 µg Saci_1104, 400 µg Saci_1085, and/or 200 µg of Saci_1115 by mixing the respective concentrated protein preparations after affinity chromatography and incubation on ice for 5 min followed by SEC as described above. For calibration, Gel Filtration LMW Calibration Kit and Gel Filtration HMW Calibration Kit (Cytavia, USA) were used.

### Coelution experiments for protein complex analyses

To check for protein-protein interaction of C-terminally twin-strep-tagged Saci_1121/Saci_1120 with N-terminally his-tagged Saci_1104 a co-purification assay was performed. 1 mg of twin-strep-tagged Saci_1121/Saci_1120 was loaded onto a Strep-TactinXT (IBA, Germany) column (column volume of 250 µl) followed by washing with 20 column volumes of wash buffer (100 mM TRIS-HCl (pH 8), 150 mM NaCl, 1 mM EDTA. Afterwards, 1.5 mg of N-terminally his-tagged Saci_1104 was applied to the column and unbound protein was removed by washing with 20 column volumes of wash buffer. Bound proteins were eluted with 500 µl of elution buffer (100 mM TRIS-HCl (pH 8), 150 mM NaCl, 1 mM EDTA, 50 mM biotin). In the reciprocal experiment 1 mg of N-terminally his-tagged Saci_1104 was loaded onto a Nickel-NTA (Sigma, Germany) column (column volume of 250 µl), washed with 20 column volumes of wash buffer (50 mM TRIS-HCl (pH 8), 250 mM NaCl). Then, 1 mg of C-terminally twin-strep-tagged Saci_1121/Saci_1120 was applied to the column followed by removal of unbound protein with 20 column volumes of the same wash buffer. Elution of bound protein was performed with 50 mM TRIS-HCl (pH 8), 250 mM NaCl, 300 mM Imidazol). Coelution of the proteins indicate interaction/complex formation.

To confirm the formation of a four-protein complex 1 mg of his-tagged Saci_1104 was loaded onto a 250 µl Nickel-NTA column and washed with 20 column volumes of wash buffer (same as above). Afterwards, the other proteins were consecutively applied to the column:(1) 1.5 mg C-terminally twin-strep-tagged Saci_1121/Saci_1120, followed by (2) 1.5 mg of N-terminally twin-strep-tagged Saci_1085. After washing (as before), proteins were eluted with 500 µl of elution buffer (see above). Eluted samples were analysed by SDS-PAGE and immunoblotting using antibodies against either his-tag (Anti-6x His tag; Abcam, United Kingdom) or twin-strep-tag (Strep-Tactin HRP conjugate; IBA, Germany). To exclude unspecific binding either only twin-strep-tagged Saci_1121/Saci_1120 or N-terminally twin-strep-tagged Saci_1085 were applied to the column as a control.

### Enzyme assays

The activities of all the enzymes were determined either via continuous, photometric assays or in discontinuous assays analyzed by HPLC. All the reaction mixtures were pre-incubated under the assay conditions for 2 min, and reactions were started by addition enzymes.

*Acyl-CoA dehydrogenase (ACAD)* – ACAD activity of Saci_1123 was measured at 65°C, in 50 mM HEPES-KOH (pH 6.5) containing 20 mM KCl. Continuous spectrophotometric assays were based on the acyl-CoA dependent reduction of the artificial electron acceptor ferrocenium (FcPF_6_) to its reduced species ferrocene at 300 nm (extinction coefficient: 4.3 mM^-1^ cm^-1^ (*36*)). The reaction mixture (500 µl) contained 1 mM FcPF_6_, 0.13 µg/µl Saci_1123 and 0.4 mM acyl-CoAs (butyryl-CoA, hexanoyl-CoA, octanoyl-CoA and palmitoyl-CoA). Variable concentrations of octanoyl-CoA (0-0.15 mM) were used for determination of *V*_max_ and *K_M_*. For oxidation of palmitoyl-CoA 2.5% (v/v) DMSO and 0.1% (v/v) Triton were added to increase solubility of the substrate. The discontinuous assay (50 µl) containing 50 mM MES-KOH (pH 6.5), 20 mM KCl, 0.4 mM DCPIP, 0.02 µg/µl ACAD and 0.01 µg/µl ETF with 0.4 mM of acyl-CoAs (butyryl-CoA, hexanoyl-CoA or octanoyl-CoA) were incubated for 5 min at 65°C, reactions were stopped and substrate consumption and product formation were detected via HPLC (see below). Moreover, 0.8 mM FcPF_6_ was also used as an electron acceptor instead of ETF and DCPIP.

*Electron transfer flavoprotein (ETF)* – Determination of ETF Saci_0315 activity was performed using the artificial electron acceptor DCPIP. The discontinuous assay was based on the fact that ACADs are not able to directly transfer electrons to DCPIP but instead need an intermediate electron acceptor like ETF. Thus, activity was measured as ETF dependent reduction of DCPIP by ACAD. The assay mixture (0.5 ml) containing 50 mM MES-KOH (pH 6.5), 20 mM KCl, 0.2 mM DCPIP, 0.2 mM of different acyl-CoAs (butyryl-CoA, hexanoyl-CoA or octanoyl-CoA), 0.0034 µg/µl ACAD and 0.0076 µg/µl ETF was incubated at 65°C for 5 min. Afterwards, the DCPIP spectrum in each sample was determined in the range of 400 to 800 nm and DCPIP reduction was indicated by depletion of the absorption maximum at 600 nm. Additionally, the NADH oxidizing activity of Saci_0315 was determined with iodonitrotetrazolium chloride (INT) as electron acceptor concomitantly reduced to the red formazan at 500 nm (extinction coefficient 19.3 mM^-1^ cm^-1^). The assay was performed in 50 mM HEPES-NaOH (pH 7.5) containing 100 mM NaCl, 0.2 mM INT and 0.015 µg/µl ETF protein with 0-1 mM NADH for *K_M_* and *V*_max_ determination. To measure the potential electron transfer from NADH to Crotonyl-CoA 0.0034 µg/µl ACAD and 0.0076 µg/µl ETF were incubated with 0.4 mM Crotonyl-CoA and 0.2 mM of NADH and NADH oxidation was followed at 340 nm. As a control the electron transfer from NADH to DCPIP via ETF was measured as described.

*3-Hydroxyacyl-CoA dehydrogenase/enoyl-CoA hydratase (HCDH/ECH) –* The combined activity of the bifunctional protein HCDH/ECH Saci_1109 was continuously determined at 75°C as crotonyl-CoA dependent reduction of NAD(P)^+^ at 340 nm in 100 mM TRIS-HCl (pH 7) with 0.4 mM crotonyl-CoA, 0.2 mM NAD^+^ or NADP^+^ and 0.00138 µg/µl protein (total volume 500 µl). *K_M_* values for crotonyl-CoA and NAD^+^ were determined by varying the concentrations from 0-1.6 mM and 0-0.3 mM, respectively. Temperature and pH optima were determined under the same conditions using a mixed buffer consisting of 50 mM MES, 50 mM HEPES and 50 mM TRIS adjusted with NaOH. The combined activity of HCDH/ECH as well as the ECH partial reaction was also measured discontinuously in 50 mM MES-KOH (pH 6.5 at 65°C) containing 20 mM KCl, 0.0144 µg/µl protein and 0.4 mM crotonyl-CoA with 2 mM NAD^+^ (combined HCDH/ECH activity) or without NAD^+^ (EHC activity). The reactions were incubated for 15 min, stopped and substrate consumption and product formation were detected via HPLC (see below). This assay system was also used to measure the combined conversions of the last three steps of β oxidation, i.e. the CoA and NAD^+^ dependent formation of acetyl-CoA from crotonyl-CoA via HCDH/ECH and KT (see below) by addition of 1.6 mM CoA and 0.054 µg/µl KT. The 3-HBCoA oxidizing partial reaction was measured using 0.4 mM of the commercially available racemic mixture 3(S/R)-HBCoA or the synthesized steroisomerically pure 3(S)-or 3(R)-HBCoA instead of crotonyl-CoA in the same assay. *K_M_* values for mixed 3(S/R)-HBCoA (0-0.7 mM), NAD^+^ (0-1.5 mM) and 3(S)-HBCoA (0-1.25 mM) were determined. In the reductive direction, HCDH activity of the fusion enzyme was measured as AcAcCoA dependent oxidation of NADH at 35°C due to instability of the substrate. The assay (500 µl) contained 100 mM TRIS-HCl (pH 7), 0.6 mM AcAcCoA, 0.2 mM NADH or NADPH and 0.0081 µg/µl HCDH/ECH enzyme. *K_M_* values for AcAcCoA (0-0.7 mM), NADH (0-1.5 mM) or NADPH (0-1.25 mM) were determined.

*β-Ketothiolase (also known as acetyl-CoA C-acetyltransferase) (KT)* – CoA dependent cleavage of acetoacetyl-CoA into two acetyl-CoA by KT was investigated continuously at 23°C (due to thermal instability of AcAcCoA) by monitoring the consumption of AcAcCoA-Mg^2+^ chelation complex as an decrease in absorbance at 303 nm (extinction coefficient of Mg^2+^-AcAcCoA complex: 21.4 mM^-1^ cm^-1^ (*37*)) The reaction mixture (500 µl) contained 100 mM TRIS-HCl (pH 8), 20 mM MgCl_2_, 0.2 mM CoA, 0.1 mM AcAcCoA, and 0.0054 µg/µl (Saci_1114) or 0.0036 µg/µl (Saci_1121/1120) enzyme. Variable concentrations of AcAcCoA (0-0.2 mM) and CoA (0-0.2 mM) were used for determination of *K_M_* values, respectively. KT activity was discontinuously measured in 100 µl total assay volume containing 50 mM MES-KOH (pH 6.5), 0.2 mM CoA and 0.1 mM AcAcCoA. After preincubation at 23°C for 2 min, the reaction was started by addition of 0.027 µg/µl KT and incubated for further 5 min followed by HPLC analyses of substrates and products (see below). The KT activity in the direction of the Claisen condensation of two molecules of acetyl-CoA was measured at 75°C by coupling AcAcCoA formation to NADH oxidation via the Saci_1109 HCDH/ECH as auxiliary enzyme at 340 nm. The assay mixture (500 µl) contained 100 mM MOPS-NaOH (pH 6.5), 0.3 mM NADH, 0.075-7.5 mM acetyl-CoA (for *V*_max_ and *K_M_* determination) as well as 0.0342 µg/µl ECH/HCDH and 0.0216 µg/µl Saci_1114, or 0.02052 µg/µl HCDH/ECH and 0,0104 µg/µlSaci_1121/1120, respectively. Furthermore, for Saci_1114 0.5 mM of respective acyl-CoAs (acetyl-CoA, butyryl-CoA, hexanoyl-CoA, octanoyl-CoA, lauroyl-CoA or palmitoyl-CoA) was added in addition to 2.5 mM acetyl-CoA to investigate the chain length specificity. For Saci_1121/Saci_1120 0.5 mM of respective acyl-CoAs (acetyl-CoA, butyryl-CoA, hexanoyl-CoA, octanoyl-CoA) were added to 1 mM of acetyl-CoA. The discontinuous assay contained in 400 µl total volume 100 mM MOPS-NaOH (pH 6.5), 1 mM acetyl-CoA, 0.3 mM NADH, 0.04275 µg/µl HCDH/ECH and 1 mM of the respective acyl-CoA (acetyl-CoA, butyryl-CoA, hexanoyl-CoA or octanoyl-CoA). After preincubation at 65°C for 2 min, 0.0405 µg/µl KT was added to start the reaction followed by further incubation for 5 min. Substrates and products were analyzed via HPLC.

The potential of Saci_1114 KT to utilize malonyl-CoA as extender unit was studied by incubation of 0.0054 µg/µl of Saci_1114 under the same conditions as described above with 0.2 mM NADH and 0.0342 µg/µlHCDH/ECH. As substrates either 2 mM of acetyl-CoA, 1 mM butyryl-CoA and 1 mM acetyl-CoA, or 1 mM butyryl-CoA and malonyl-CoA were used and absorption changes were followed at 340 nm.

*β Oxidation enzyme cascades for HPLC analysis* – All enzyme cascade (total volume 50 µl) studies were performed discontinuously in two steps. The first oxidation step by ACAD and ETF was done as described above at 65°C using DCPIP as electron acceptor and ran to completion. In the second step, 2 mM NAD^+^, 0.0144 µg/µl HCDH/ECH, 1.6 mM CoA and 0.054 µg/µl KT were added followed by incubation for 15 min. Substrate consumption as well as intermediate and product formation was monitored via HPLC.

*Acetoacyl-CoA reductase (ACR)* – ACR activity was determined at 35°C as AcAcCoA dependent oxidation of NADH/NADPH at 340 nm. The assay mixture (500 µl) contained 100 mM TRIS-HCl (pH 7), 0.3 mM AcAcCoA, 0.2 mM NADH/NADPH and 0.00806 µg/µl protein. The concentration of AcAcCoA or NADPH was varied from 0-0.5 mM or 0-0.2 mM, respectively, for *K_M_* determination while concentrations of the other components were kept constant. In the oxidative direction, ACR activity was assayed at 70°C using 3-HBCoA as substrate and 2 mM NAD^+^ or NADP^+^ as cosubstrate in 100 mM TRIS-HCl (pH 7) with 0.04032 µg/µl protein by following the increase in absorbance at 340 nm. Concentrations of 0-2 mM were used for *K_M_* measurement with the racemic mixture of 3(S/R)-HBCoA and 0-1 mM for stereoisomerically pure 3(R)-HBCoA in presence of 2 mM NADP^+^. The pH optimum was determined in the direction of 3-hydroxybutyryl-CoA formation at 35°C using 100 mM MES-NaOH (pH 5.5-6.5). The temperature optimum was determined in the direction of acetoacetyl-CoA formation using 100 mM TRIS-HCl (pH 7), at the respective temperature.

*MaoC-like 3(R)-hydroxyacyl-CoA dehydratase (MaoC-HCD)* – Both, the enoyl-CoA hydrating and 3(R)-hydroxyacyl-CoA dehydrating activity of reversible MaoC-HCD Saci_1085 was determined at 65°C in a discontinuous assay containing (total volume 50 µl) 50 mM MES-KOH (pH 6.5), 20 mM KCl, 0.0675 µg/µl protein, and the respective substrate in the indicated concentrations. After incubation for 2, 5, 10, and 30 min aliquots were taken, the reaction was stopped and substrate consumption and product formation was analyzed via HPLC (see below). *K_M_* values for 3(R)-hydroxybutyryl-CoA and crotonyl-CoA were determined by varying their concentration from 0-0.5 mM and from 0-1 mM, respectively. The chain length specificity for enoyl-CoA substrates was analysed by incubating 0.09 µg/µl protein with 0.4 mM crotonyl-CoA, hexenoyl-CoA or octenoyl-CoA. The substrate hexenoyl-CoA and octenoyl-CoA were enzymatically produced by ACAD, ETF and DCPIP as mentioned above. To study the stereospecificity 0.4 mM of stereoisomerically pure 3(S)-or 3(R)-hydroxybutyryl-CoA was used and the reaction was run for 30 min.

*Enoyl-CoA reductase (ECR)* – ECR activity was assayed photometrically at 70°C in a continuous assay as enoyl-CoA dependent oxidation of NAD(P)Hat 340 nm. The standard assay (500 µl) contained 100 mM HEPES-NaOH (pH 7.5), 10 mM KCl, 0.3 mM NADH or NADPH, 0.4 mM crotonyl-CoA and 0.024 µg/µl ECR. *K_M_* values were determined with 0-5 mM crotonyl-CoA and 0-0.04 mM NADPH, respectively. Butyryl-CoA oxidizing activity was disproved with 0.2 mM butyryl-CoA as substrate and 1 mM NAD^+^/NADP^+^ as cofactor under the same assay conditions (500 µl). The pH optimum for ECR was determined at 70°C in a pH range of 5.5-9.0 using a mixed buffer system of 50 mM HEPES, 50 mM TRIS and 50 mM MES adjusted with NaOH with 0.016 µg/µl protein. To analyse the substrate specificity of ECR, hexenoyl-CoA (C6) and octenoyl-CoA (C8) were enzymatically synthesized by ACAD (Saci_1123) from the saturated derivatives followed by the NADPH dependent enoyl-CoA reduction monitored at 340 nm. Both reactions were consecutively performed in the same cuvette at 70°C in a total volume of 500 µl. For enoyl-CoA production, the reaction mixture initially contained 100 mM HEPES-NaOH (pH 7.5), 0.3 mM of hexanoyl-CoA or octanoyl-CoA, 0.6 mM of FcPF_6_. 0.01508 µg/µl ACAD (Saci_1123) was added and the conversion of acyl-CoA to enoyl-CoA was run to completion (as judged at 300 nm). Then, the continuous assay was started by the addition of 0.2 mM NADPH, 0.02 µg/µl ECR and the decrease in absorbance was monitored at 340 nm. C10 and C16 2,3-enoyl-CoAs were chemically synthesized as described above and the continuous assay was used with 0.2 mM NADPH, 10 ug protein and 0.3 mM enoyl-CoA.

*Fatty acid synthesis enzyme cascades for HPLC analysis* –The conversion of 3(R)-hydroxyacyl-CoA via the corresponding enoyl-CoA into acyl-CoA was analyzed at 65°C in a discontinuous assay (total volume 50 µl), containing 50 mM MES-KOH (pH 6.5), 20 mM KCl, 0.0168 µg/µl MaoC-HCD Saci_1085, 0.0246 µg/µl ECR Saci_1115, 2 mM NADPH, and 0.4 mM 3(R)-HBCoA. After 30 min of incubation samples were analyzed for substrate consumption as well as intermediate and product formation via HPLC (see below). For the analyses of butyryl-CoA formation from AcAcCoA the same assay (total volume 50 µl) was used substituting 3(R)-HBCoA for 0.4 mM AcAcCoA and including 0.104 µg/µl ACR (concentration of the other components same as above). After 30 min or in regular intervals between 0 and 240 min (to analyze time dependence of the reaction(s)), samples were withdrawn and analyzed via HPLC. The conversion of 1 mM acetyl-CoA (instead 3(R)-HBCoA or AcAcCoA) to saturated acyl-CoA esters by the complete fatty acid synthesis enzyme cascade including 0.072 µg/µl µg KT/DUF35 Saci_1121/1120, 0.104 µg/µl ACR Saci_1104, 0.0168 µg/µl MaoC-HCD Saci_1085, and 0.0246 µg/µl ECR Saci_1115 in a total volume of 400 µl, was studied in the same buffer system. After 0h, 3h, 6h, and 21h aliquots were removed, reactions were stopped and samples were analyzed via HPLC as described below.

### HPLC analysis of the fatty acid metabolic intermediates from enzyme conversions

The discontinuous enzymatic conversions (see above) were stopped by addition of acetonitrile in a ratio sample/acetonitrile of 1:3 (v/v). Samples were incubated at-80°C for 20 min or at-20°C overnight. Precipitate proteins were removed via centrifugation at 4°C for 30 min. The supernatants were transferred to fresh tubes and the liquid was evaporated completely at 30°C using an Eppendorf^TM^ Concentrator Plus. The remaining solid was resuspended in ultrapure water and applied to the Thermo Scientific UltiMate 3000 HPLC system (Thermo Fisher Scientific US). CoA esters were separated via a reversed phase NUCLEODUR C18 Pyramid analytical column (MACHEREY-NAGEL GmbH & Co. KG, Germany) using two different HPLC programs: For shorter chain CoA esters (C ≤ 4), the column was pre-equilibrated with 96% of Buffer A (0.2 M ammonium acetate, pH 5) and 4% of Buffer B (ACN), CoA and CoA esters were eluted employing three linear gradients of ACN: (i) 0-20 min, 4-7% can, (2) 20-35 min, 7-30% ACN, and (3) 35-35.5 min, 30-4% followed by an isocratic flow with 4% Buffer B for 9.5 min. This HPLC program is herein referred to as the “4-30% ACN” program. For longer chain CoA esters (C ≥ 4), the column was pre-equilibrated with 99% of Buffer A and 1% of Buffer B, then acyl-CoA compounds were eluted with a linear gradient of 1 to 10% ACN for 5 min followed by an isocratic flow with 10% ACN for 16 min, then two linear gradients of 10 to 60% for 20 min and 60-1% for 0.5 min were applied. Afterwards, the system was run with 1% ACN for further 19.5 min. This second HPLC program for detecting longer chain CoA esters was referred to as “1-60% ACN” program. All HPLC analyses were carried out at 8°C and a flow rate of 1 ml/min. Data collection and processing was done by Thermo Scientific™ Dionex™ Chromeleon 7 Chromatography Data System (CDS) Software (Thermo Fisher Scientific US). The retention times of the relevant commercially available CoA derivatives were determined (Fig. S13) and are listed in Table S5. The identification of peaks was based on chromatography with standards and analysis of UV spectra of formed products.

### Construction of markerless ECR deletion mutants Δ*saci_1115*

To obtain the markerless deletions of *saci_1115*, the knock-out plasmid 1115 was constructed. Briefly, 500bp of the upstream and downstream region of the *saci_1115* gene were amplified by PCR and the respective PCR products were annealed via overlap extension PCR. The resulting products were then cloned into pSVA407 (for plasmids and strains, see Table S3). The resulting plasmids were then methylated by transformation in *E. coli* ER1821 to prevent plasmid degradation in the recipient strain. Methylated plasmid was then used to transform MW00G as described previously for MW001 (*31*). To prepare competent MW00G, a pre-culture was grown in Brock media supplemented with 0.1 % (w/v) NZ-amine and 0.2 % (w/v) dextrin and 20 µg/mL uracil to an OD_600_ of 0.5-0.7 and used to inoculate 50 mL Brock media supplemented with 0.1 % (w/v) NZ-amine and 10mM glycerol. The culture was harvested at an OD_600_ of 0.2-0.3 and transformation and mutant selection was performed as described previously (*31*).

### Analysis of Amino Acids by LC-MS

To confirm isotopic labeling of the *S. acidocaldarius* samples, known metabolites e.g. amino acids were analyzed. Therefore, the samples were extracted using a series of freezing and thawing cycles adapted from (*38*). Therefore 50 mg wet sample material was weight and placed in a 1.5 mL cup. The sample was resuspended in 500 µL prechilled MeOH at-80 °C. After vortexing for 2 min, sonication was performed for another 2 min. The sample was then frozen at-80 °C for 5 min, followed by thawing with vortexing for 2 min and repeated sonication for 2 min. The sample was centrifuged for 5 min at 1,400g and the MeOH was collected in a separated vial. The remaining sample was reextracted with 250 µL water and after repeating the vortexing and sonication steps, the sample was frozen again at-80 °C for 5 minutes. The sample was thawed while vortexing for 2 min. After centrifugation at 1,400g the supernatant was collected, combined with the MeOH phase and dried in a vacuum centrifuge. The sample was resuspended in 200 µL of ACN/water (90/10 v/v) and subjected to a 2 min sonication and 2 min vortexing. After centrifugation at 10,000 g for 2 min the supernatant was used for LC-MS analysis.

LC-MS measurements were conducted using an Agilent 1290 Infinity II LC system equipped with an Agilent AdvanceBio MS Spent Media column (2.1 × 150 mm) obtained from Agilent Technologies (Waldbronn, Germany). The mobile phases consisted of 10 mM ammonium acetate in water (pH 9) for phase A and 10 mM ammonium acetate in acetonitrile/water (9/1, v/v) (pH 9) for phase B. The gradient initiated at 90% B for 2 min, decreased linear to 40% B in 12 min, linear to 20% B at 13 min, maintained at 20% B until 16 min, and returned to 90% B for equilibration. The flow rate was set at 0.25 mL/min. Detection was carried out using the Agilent 6545 Q-ToF-MS from Agilent Technologies (Waldbronn, Germany. Ionization parameters were gas temperature 250 °C, 10 L/min drying gas, nebulizer at 40 psi, sheath gas temperature at 300 °C, sheath gas flow at 12 L/min, capillary voltage at 3000 V, nozzle voltage at 0 V, fragmentor voltage at 125 V, and skimmer voltage at 85 V. The Q-ToF-MS system operated in negative mode, with a mass range of 50-1000 m/z, acquiring spectra at 333 ms/spectra. For data analysis extracted ion chromatograms of known metabolites (e.g. amino acids) were generated for the [M-H]-ions for monoisotopic ^12^C and fully labeled ^13^C compounds (Fig. S5).

### Fatty acid analysis by GC-MS

Fatty acids were analyzed by GC-APCI-QqQ-MS (*11*). Extraction of the samples was performed using MTBE (*39*). Briefly, 50 mg wet sample material was weight and placed in a 1.5 mL glass vial and. The sample were resuspended in 100 µL internal standard solution containing decanoic acid (^2^H₂-FA 10:0), pentadecanoic acid (^2^H₂-FA 15:0), octadecanoic acid (^2^H₄-FA 18:0), 5,8,11,14-eicosatetraenoic acid (^2^H₈-FA 20:4 Δ5,8,11,14), and tetracosanoic acid (^2^H₄-FA 24:0) each at a concentration of 500 nM in MeOH. For hydrolysis of esterified fatty acids, 60 µL of 10 M potassium hydroxide were added to the samples. After homogenization in a cooled ultrasonic bath for 5 minutes, the samples were incubated at 60 °C for 30 minutes. The sample was acidified with 70 µL acetic acid. After a vortexing, 300 µL of methanol was added. The samples were homogenized in an ultrasonic bath for 5 min. Subsequently, 600 µL of MTBE was added, and the mixture was vortexed for an additional 5 min. Following this, 300 µL of water was introduced, and the combination was vortexed for 5 min. The upper phase was collected, and the aqueous phase underwent a second wash with 300 µL of MTBE. The resulting combined MTBE phases were then dried using a vacuum centrifuge at 45 °C.

The dried extracts were derivatized by the addition of 20 µL of 10% diisopropylethylamine in dichloromethane (1/9; v/v) and 20 µL of 10% 2,3,4,5,6-pentafluorobenzylbromid in dichloromethane (1/9; v/v). After incubation at 50 °C for 1 h the samples were dried in a vacuum centrifuge at 45 °C and redissolved in 100 µL MeOH for analysis.

The analyses were conducted using a Agilent 7890B GC with an autoinjector, utilizing a DB-5 column (30 m × 250 μm × 0.25 μm). The GC was couple to a 6495 triple Quad MS unsing a GC-APCI source (Agilent Technologies, Waldbronn, Germany). A 1 µL sample was injected into a split liner at 320 °C, with a split ratio of 1:10 and a septum purge flow of 1.5 mL/min. The column flow, operated with helium as the carrier gas, was set to 1.3 mL/min. The temperature gradient followed this profile: starting at 100 °C, with a linear increase of 15°/min to 160 °C, then a linear increase of 5°/min to 320 °C, which was maintained for 5 minutes. Detection was carried out in negative ionization and pseudo SRM mode with the m/z ratios of the [M-PFB]^−^ ions set as *m/z* for Q1 and Q3 with no fragmentation energy applied. To monitor the ^13^C labeled version of the fatty acids, the *m/z* ratios of the fully labeled compounds were used. In total 75 biologically relevant fatty acids were monitored.

### Mass determination by native MS analysis

KT/DUF35 (Saci_1121/1120) was rebuffered to 2.5 mM ammonium acetate buffer (NH_4_OAc; pH 6.8; Sigma-Aldrich, A2706; diluted in MS-grade water, Honeywell, 14263) using 10 kDa molecular weight cutoff spin-filter columns (UFC501096, Millipore). The protein concentration was quantified via microvolume spectroscopy (DeNovix, DS-11+) and adjusted to 1 µM. The protein sample was ionized using a TriVersa NanoMate nanoESI system (Advion), which was equipped with a 5 µm diameter nozzle spray chip (Advion, HD_A_384). The ESI spray was generated with 0.8 psi nitrogen backpressure and a positive nozzle chip voltage of 1.7 kV. MS spectra were recorded for a duration of two minutes in positive EMR mode on an Exactive Plus EMR Orbitrap mass spectrometer (ThermoFisher). For mass deconvolution, the resulting data was processed using UniDec software (v6.0.4). All settings that deviate from the standard UniDec settings are listed in Table S6. GraphPad (version 8.0.1) was used for data visualization.

### Phylogenetic analysis of fatty acid synthesis genes

For each target gene sequence the seed-ortholog sequence and corresponding non-supervised orthologous group (NOG) at the LUCA level (i.e., including all prokaryotic and eukaryotic proteins) were identified using the online eggnog-mapper web service (http://eggnog-mapper.embl.de/). The below described workflow was employed for all target NOGs with the SnakeMake v6.4.1 (*40*) workflow management system.

Member protein sequences of the target NOG were downloaded using the EggNOG v5 web API (*41*). Proteins were clustered at 80% sequence identity with CD-HIT v4.8.1 (*42*) and multiple sequence alignment (MSA) was performed using Clustal Omega v1.2.3 (*43*).

The generated MSA was used as a query for a profile search with the “diamond blastp —more-sensitive” program of DIAMOND (*44*) against a local copy of the NCBI’s NR database (*45*) v5 on April 9th, 2021. Significant hits (E< 1e-5) were clustered at 80% identity as above and recruited to the clustered target NOG. The clustered, extended NOG protein sequence dataset was aligned with Clustal Omega. Columns from the resulting MSA were removed if they contained over 20% gaps using trimAl v1.4.1 (*46*), and full sequences were removed if they did not cover over 20% of the trimmed alignment. An initial phylogenetic tree was inferred using FastTree 2 v2.1.11 (*47*). Members of the focal clade containing the specified seed-ortholog were extracted and MSA was performed using Clustal Omega on the full sequences. Trimming of the resulting MSA was performed as described above and a phylogenetic tree was inferred with FastTree as above.

To make phylogenetic analysis with complex models feasible, representative sequences were identified in the reconstructed phylogenetic tree that retain 80% of the phylogenetic diversity by applying Treemmer v0.3 (*48*). We adapted the Treemmer script to preferentially remove sequences recruited from NCBI nr and retain the original NOG member sequences.

We performed MSA in both the forward and reverse direction and merged the two alignments with T-Coffee v13.45.0 (*49*). MSA trimming was performed as above. A first round of maximum-likelihood (ML) phylogenetic reconstruction was performed using IQ-Tree v1.6.9 (*50*) with the best fit model identified by modelfinder (*51*) and 1,000 improved ultrafast bootstraps.

We ran TreeShrink v1.3.7 (*52*) with a quantile threshold of 0.01 and centroid rerooting with “--mode per-gene” to identify outlier long branches and RogueNaRok (*53*) to identify putative rogue sequences on the first round ML phylogenetic tree, respectively. After removing flagged outlier and rogue sequences, we performed and exhaustive MSA using MAFFT v7.453 (*54*) “--localpair--maxiterate 1000”, MAFFT “--geneafpair--maxiterate 1000”, and iteratively refined Clustal Omega “--iter 3” both for the forward and reverse direction that were subsequently merged with T-Coffee “-evaluate_mode=t_coffee_slow” and subsequently trimmed with trimAl “-gappyout”. We used the resulting alignments to perform phylogenetic reconstruction using more complex and realistic models of sequence evolution. We ran IQ-Tree with the flags “-m TESTNEW-mset LG,LG+C10,LG+C20,LG+C30,LG+C40,LG+C50,LG+C60,LG4M,LG4X-mrate,G4,R4,R5,R6” and calculated 1,000 improved ultrafast bootstraps. In all cases LG+C60+F+R6 was chosen as the best-fit model. Using this initial ML trees as guides we reconstructed the posterior mean site frequency (PMSF) approximation ML trees of the with the best-fit model with 100 non-parametric bootstraps. For three COGs (COG1028, COG1064, and COG2030), we repeated the steps described in this paragraph another time to remove rogue and outlier sequences. Trees were visualized with FigTree v1.4.4 (http://tree.bio.ed.ac.uk/software/figtree/).

### Syntenic gene blocks analysis

To identify putative syntenic gene blocks including *S. acidocaldarius* fatty acid synthesis related genes, we downloaded the predicted proteomes of 2339 representative archaeal genomes from the GTDB (*55*). We annotated genomes with emapper v97dad3b (*56*) using EggNOG db v5.0.1. We ran colinear syntenic blocks (CSB) finder (*57*) with “-q 2-s 3” parameters. Results were processed and visualized in R v4.2.1 (R Core Team 2018) with ggplot2 3.4.3 (*58*).

## Abbreviations

(ACAD): acyl-CoA dehydrogenase
(ETF): electron transfer flavoprotein
(HCDH/ECH): 3(S)-hydroxyacyl-CoA dehydrogenase/enoyl-CoA hydratase
(KT): β-ketothiolase or acetyl-CoA C acetyltransferase
(ACR): acetoacetyl(ketoacyl)-CoA reductase
(MaoC-HCD): MaoC-like 3(R)-hydroxyacyl-CoA dehydratase
(ECR): enoyl-CoA reductase
(SDR): short-chain dehydrogenases/reductases superfamily
(MDR): medium-chain dehydrogenases/reductases superfamily
(FAD): flavin adenine dinucleotide
(DCPIP): 2,6-dichlorophenolindophenol
(INT): iodonitrotetrazolium
(DTNB): 5,5’-dithiobis-(2-nitrobenzoic acid) or Ellman’s reagent
(F_c_PF6): ferrocenium hexafluorophosphate
(DTT): dithiothreitol
(ACN): acetonitrile

## Data availability

The original contributions presented in this study are included in the article/supplementary material, further inquiries can be directed to the corresponding author.

## Author contributions

Christian Schmerling: Formal analysis, Investigation, Validation, Visualization, Writing – Original Draft.

Xiaoxiao Zhou: Formal analysis, Investigation, Validation, Visualization, Writing – Original Draft.

Paul Eric Görs: Formal analysis, Investigation, Validation, Visualization

Stephan Köstlbacher: Formal analysis, Investigation, Validation, Visualization, Writing – Original Draft

Till Kessenbrock Formal analysis, Investigation, Validation, Visualization

David Podlesainski Formal analysis, Investigation, Validation, Visualization

David Sybers Formal analysis, Investigation, Validation

Kun Wang Formal analysis, Investigation, Validation

Ann-Christin Lindås: Supervision, Validation, Writing – Reviewing and Editing

Jacky Snoep: Formal analysis, Validation, Data curation, Investigation, Methodology, Visualization, Writing – Reviewing and Editing.

Eveline Peeters: Supervision, Validation, Writing – Reviewing and Editing

Markus Kaiser: Conceptualization, Methodology, Validation, Writing – Reviewing and Editing, Visualization, Supervision, Funding Acquisition.

Thijs J.G. Ettema: Conceptualization, Methodology, Validation, Writing – Reviewing and Editing, Visualization, Supervision, Funding Acquisition.

Sven Meckelmann Conceptualization, Methodology, Validation, Writing – Reviewing and Editing, Visualization, Supervision, Funding Acquisition.

Christopher Bräsen: Conceptualization, Methodology, Validation, Writing – Original Draft, Writing – Reviewing and Editing, Visualization, Supervision, Funding Acquisition.

Bettina Siebers: Conceptualization, Methodology, Validation, Writing – Original Draft, Writing – Reviewing and Editing, Visualization, Supervision, Project Administration, Funding Acquisition.

## Acknowledgments

This research was funded by the Volkswagen Foundation. BS, CB, MK, SM and TJGE wish to express their gratitude for the financial support received for the Lipid‖Divide-“Resolving the ‘lipid divide’ by unravelling the evolution and role of fatty acid metabolic pathways in Archaea” project within the "Life?"-A fresh scientific approach to the basic principles of life initiative (grant number 96725). EP and DS were supported by the Research Foundation Flanders (FWO-Vlaanderen) (grant number G021118). DS received a strategic basic research PhD fellowship from Research Foundation Flanders (FWO-Vlaanderen) (grant number G1S19717N).

