## Supplementary Information for "*De novo* synthesis of fatty acids in Archaea via an archaeal fatty acid synthase complex"

### Supplementary Text

#### Separation of fatty acid β oxidation and biosynthesis – general considerations

Both metabolic routes, fatty acid β oxidation and fatty acid biosynthesis in Bacteria and Eukarya, are tightly regulated on the transcriptional level but also on the protein level e.g. by feedback inhibition (*1, 2*). The synthesis machinery involves the ACP for fatty acid activation and transport between the involved enzymes (*3*) whereas β oxidation relies on CoA (*2, 4*). The fatty acid synthesis is usually NADPH dependent while β oxidation uses $NAD^+$ and FAD (*5*) (the $NAD^+$/$NADP^+$ specificity is less pronounced in Bacteria (*3*)). The Claisen condensation, the first reaction in the synthesis, uses malonyl-ACP as extender unit synthesized through ATP-dependent carboxylation of acetyl-CoA. During the ketoacyl-ACP synthase catalysed condensation the $CO_2$ is liberated again, and thus energetically driven by ATP hydrolysis which is regarded as irreversible (*6*). Conversely, the ketothiolase (KT) mediated thiolytic cleavage of ketoacyl-CoAs into acetyl-CoA and acyl-$CoA_{(Cn-2)}$ in β oxidation thermodynamically strongly favours the degradation direction ($\Delta G^{0'}$ ~-25.0 kJ $mol^{-1}$)(for $\Delta G^{0'}$ values see eQulilibrator (*7*)). The following reduction of ketoacyl- to the hydroxyacyl-thioester intermediate (ketoacyl-ACP reductase; $\Delta G^{0'}$ ~-13.8 kJ $mol^{-1}$) and the dehydration to the enoyl moiety (hydroxyacyl-ACP dehydratase; $\Delta G^{0'}$ ~+3.3 kJ $mol^{-1}$) in fatty acid synthesis are in principle reversible but specific for the (R)-hydroxyacyl intermediates. Instead, the enoyl-CoA hydratase (ECH) and hydroxyacyl-CoA dehydrogenase (HCDH) catalysing the corresponding reactions in β oxidation are (S)-stereoisomer specific (*5*). Finally, the interconversion of the unsaturated enoyl- to the saturated acyl-thioester in fatty acid synthesis catalysed by NAD(P)H-dependent enoyl-ACP/thioester reductases is highly exergonic (*3*) ($\Delta G^{0'}$ ~-56.2 kJ $mol^{-1}$). In β oxidation the acyl-CoA oxidation to the enoyl-CoA is carried out by FAD dependent acyl-CoA dehydrogenases (ACADs) channelling the electrons via the electron transfer flavoprotein (ETF) and the ETF: quinone oxidoreductase (EQOR) into the quinone pool of the respiratory chain (*4, 8*). This reaction is generally regarded as irreversible in the oxidative direction especially for mechanistic reasons (*9*) and also because the electrons are exergonically transported along a redox gradient finally to a terminal electron acceptor (e.g. oxygen) via the respiratory chain (acyl-CoA/Enoyl-CoA $E^{0'}$ -0.01 V → quinone ~+0.1 V → $O_2$/$H_2O$ +0.82 V).

#### β oxidation in *S. acidocaldarius*

Consistently, our analyses with the *S. acidocaldarius* ACAD Saci_1123 show that the reaction ran to completion (with both, ETF and the artificial electron acceptor ferrocenium) (Fig. S14) but could not operate in the reductive direction. This is in line with the mechanisms described for ACAD enzymes which - in its reduced state- preferentially bind the product enoyl-CoA and thus kinetically promote the oxidative half-reaction, i.e. the electron transfer from the ACAD flavin to the electron acceptor (*8*). Furthermore, although the Saci_0315 ETF could accept electrons from NADH (Fig. S7-S8), it could not convey the electrons to re-reduce ACAD further supporting that these enzymes cannot work in the reductive direction (Fig. S8). ETF likely transfers the electrons from acyl-CoA oxidation finally to the caldariellaquinone (CQ) of the respiratory chain in *S. acidocaldarius* (*10*) via an ETF: CQ oxidoreductase (Saci_0316 (etfC) and Saci_0317 (etfX)) forming an operon with the ETF encoding gene (*saci_0315*). A similar EtfABCX complex in *Pyrobaculum aerophilum* although with non-fused single α and β subunits was recently shown to bifurcate electrons from NADH to a quinone and ferredoxin (*11*). A second *etfABCX* gene cluster with split *etfA* and *B* genes is also present in *S. acidocaldarius* (*saci_0290-0293*).

The following conversions of enoyl- to the hydroxyacyl-CoA and further to the ketoacyl-CoA, are catalysed by a bifunctional fusion protein (Saci_1109) comprised of an N-terminal HCDH and a C-terminal ECH which represents the inverted domain architecture of bacterial homologues e.g. from *E. coli* (*4, 12*)(Fig. S10H). In general, in Archaea HCDH/ECH fusion enzymes nearly exclusively show this inverted domain organization and homologues have been characterized from *Metallosphaera sedula* and *Ferroglobus placidus* (*13, 14*). Conversely, from Bacteria so far only one homologue with this inverted domain order has been characterized from *Cupriavidus necator* (*15*). In accordance with the thermodynamics, the enoyl-CoA to hydroxyacyl-CoA hydration ($\Delta G^{0'}$ -3.3 kJ mol$^{-1}$) by Saci_1109 showed an 80% conversion but a further reaction to ketoacyl-CoA could not be observed in the presence of $NAD^+$ in line with the high $\Delta G^{0'}$ (+13.8 kJ mol$^{-1}$) (Fig. S15), unless the KT Saci_1114 was added (Fig. 2, Fig. S17). The thiolytic cleavage of acetoacetyl-CoA ($\Delta G^{0'}$ -25.0 kJ mol$^{-1}$) pulled the reaction sequence from crotonyl-CoA to two acetyl-CoA in agreement with the overall energetics of $\Delta G^{0'}$-11 kJ mol$^{-1}$. The acetoacetyl-CoA conversion by Saci_1114 ran to completion (Fig. S16). Also, with C6 and C8 enoyl-CoAs produced with ACAD, acetyl-CoA formation was also observed confirming the substrate specificity of the single enzymes for the whole cascade (Fig. S18). According to the energetics, the reversal of the Saci_1109/1114 cascade was only possible to a low extend with a surplus of acetyl-CoA and NADH (Fig. S19). Furthermore, both ECH and HCDH parts of Saci_1109 were specific for the (S) stereoisomer (Table S1) which is in accordance with other β oxidation ECHs and HCDHs, whereas the enzymes catalyzing these reaction in fatty acid synthesis are specific for the (R) steroeisomers. A complex formation of bifunctional enzyme and KT observed in the canonical β oxidation (*12, 16-18*) was not observed for the *S. acidocaldarius* enzymes probably due to the inverted domain organization (Fig. S20, Fig. S10H). Together, these results show a functional archaeal β oxidation in *S. acidocaldarius* for fatty acid degradation which however does likely not operate in the reductive direction for mechanistic and thermodynamic reasons.

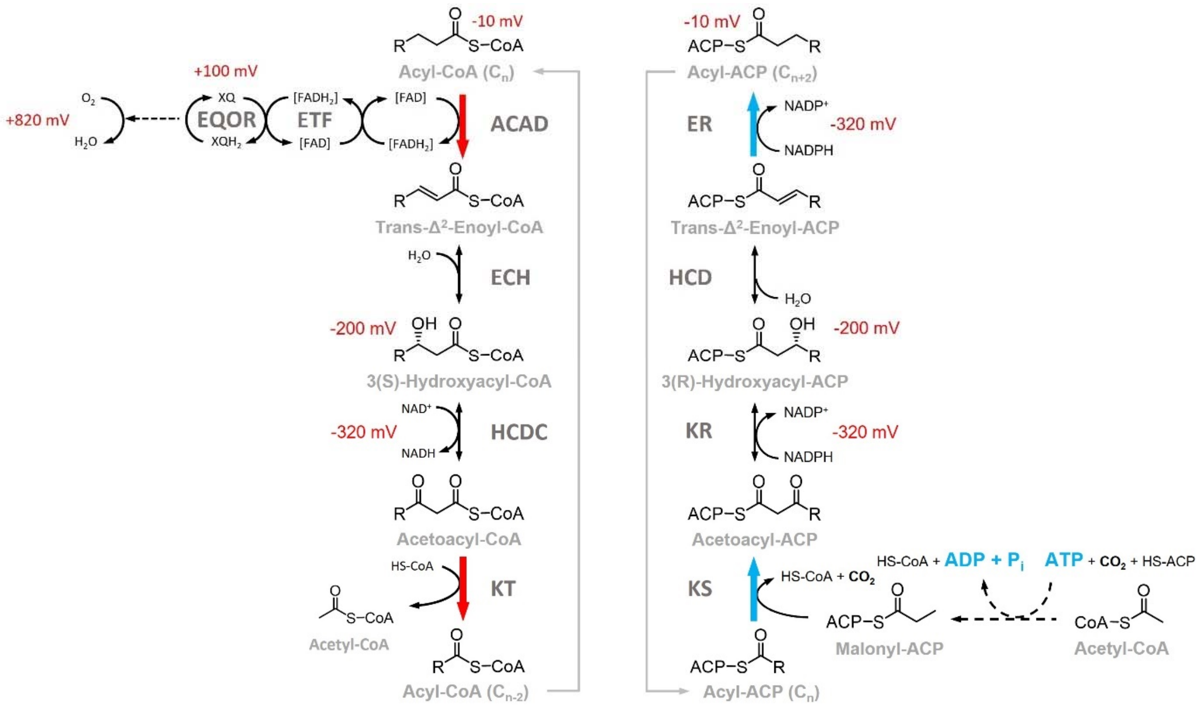


**Fig. S1. Comparative illustration of the fatty acid β oxidation (left panel) and fatty acid synthesis (right panel) as known from Bacteria and Eukarya** showing the similarities in the chemical conversions and the differences in acyl carriers, enzymes, stereospecificity, and electron carriers. The red and blue arrows indicate the main energetic (and mechanistic) driving forces in both processes (for details see supplementary text). For redox reactions the reduction potentials are given. ACAD, acyl-CoA dehydrogenase; ECH, enoyl-CoA dehydratase; HCDH, hydroxyacyl-CoA dehydrogenase; KT, ketothiolase; ETF, electron transfer flavoprotein; FAD, flavin adenine dinucleotide; $NAD(P)^+$, nicotinamide adenine dinucleotide (phosphate); EQOR, ETF:quinone oxidoreductase; KS, ketoacyl-ACP synthase; KR, ketoacyl-ACP reductase; HCD, hydroxyacyl-ACP dehydratase; ER, enoyl-ACP reductase; $P_i$, inorganic phosphate; ADP, adenosine diphosphate; ATP, adenosine triphosphate; ACP, acyl-carrier protein.

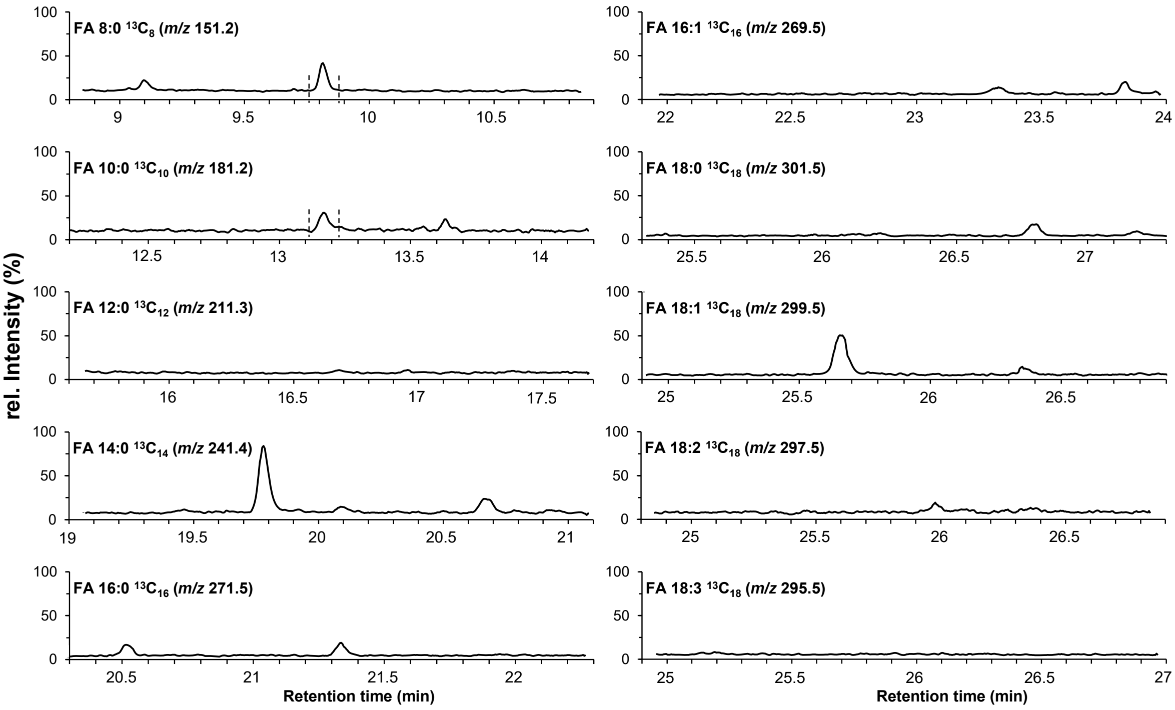


**Fig. S2. GC-MS detection of fully labelled $^{13}C$ fatty acid in $^{13}C$-glycerol grown cells of *S. acidocaldarius* MW00G.** Cells were grown for four passages on $^{13}C$-glycerol as carbon source to obtain fully $^{13}C$ labelled cells. Cells were isolated, hydrolysed before derivatization and analysis by GC-MS to characterize the total fatty acid content (for details see methods section). The chromatograms represent an overview of the raw data and display the individual mass-to-charge ratios (m/z) for each. Only for the $^{13}C$ labelled fatty acids FA 8:0 and FA 10:0 a peak with a signal-to-noise ratio of >3 at the expected retention times of 9.8 min and 13.2 min was observed (indicated by dashed lines). For fatty acids ranging from 12:0 to 18:0 including some unsaturated variants, no signal that matched the expected retention times or that significantly differs from the background of a blank sample was observed. All retention times were confirmed with $^{12}C$ standards of the corresponding fatty acid.

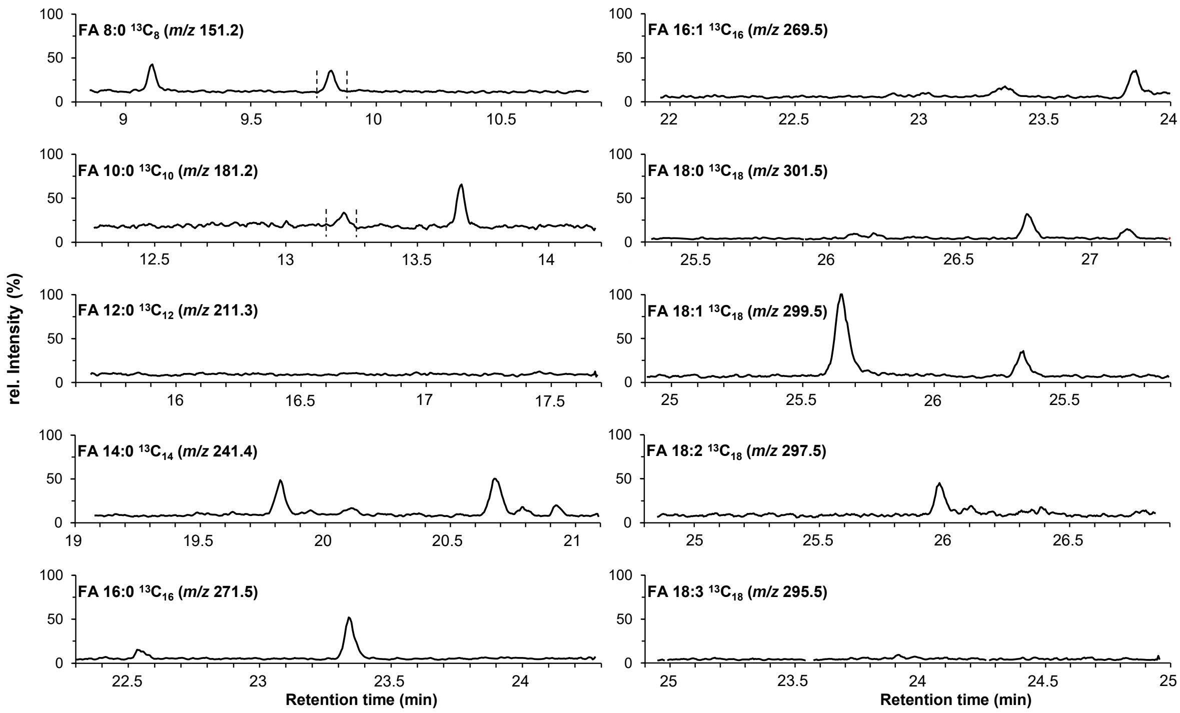


**Fig. S3. GC-MS detection of fully labelled $^{13}C$ fatty acid in $^{13}C$-glycerol grown cells of *S. acidocaldarius* wildtype MW00G Δsaci1115.** Cells were grown for four passages on $^{13}C$-glycerol as carbon source to obtain fully $^{13}C$ labelled cells. After cell isolation, the sample was hydrolyzed and derivatized to analyze the total fatty acid content (see methods for details). The chromatograms represent an overview of the raw data and display the individual mass-to-charge ratios (m/z) for each fully labelled $^{13}C$ fatty acid. Only for the $^{13}C$ labelled fatty acids FA 8:0 and FA 10:0 a peak with a signal-to-noise ratio of >3 at the expected retention times of 9.8 min and 13.2 min was observed (indicated by dashed lines). For fatty acids ranging from 12:0 to 18:0 including some unsaturated variants, no signal that matched the expected retention times or that significantly differs from the background of a blank sample was observed. All retention times were confirmed with $^{12}C$ standards of the corresponding fatty acid.

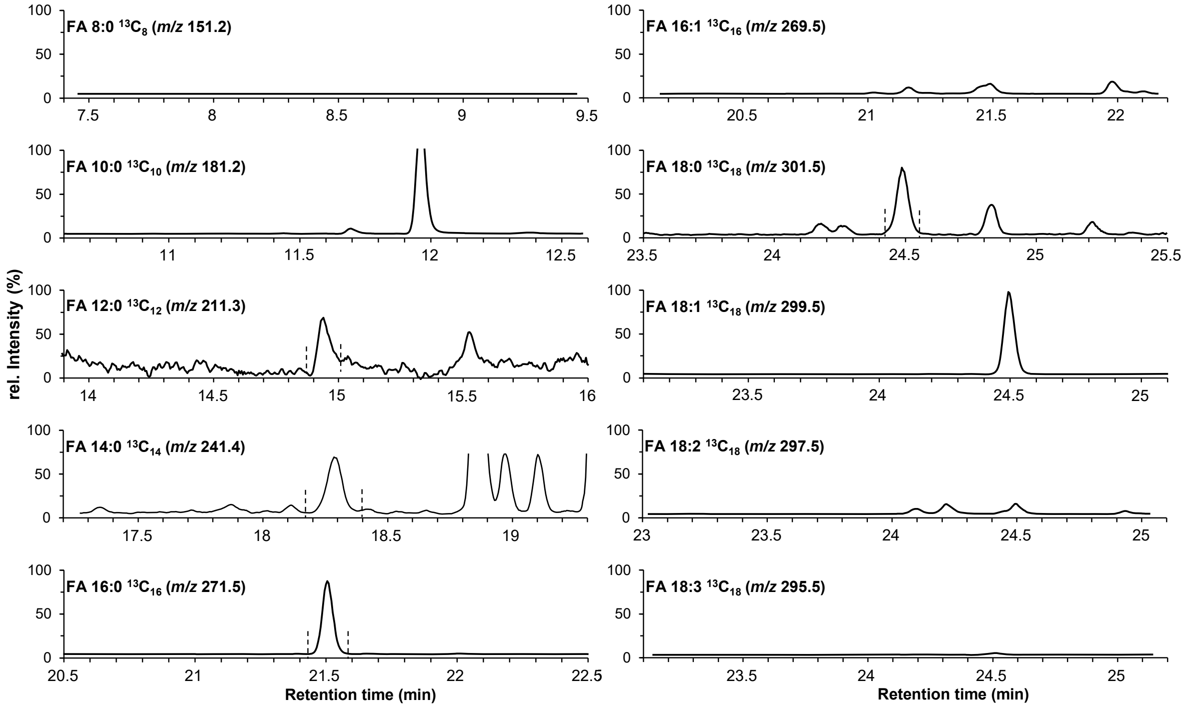


**Fig. S4. GC-MS detection of fully labelled $^{13}C$ fatty acid in $^{13}C$-glycerol grown cells from *Haloferax volcanii*.** Cells were grown for four passages on $^{13}C$-glycerol as sole carbon source to obtain fully $^{13}C$ labelled cells. After cell isolation, the sample was hydrolyzed and derivatized to analyze the total fatty acid content. The chromatograms represent an overview of the raw data and display the individual mass-to-charge ratios (m/z) for each fully labelled $^{13}C$ fatty acid. Signals with a signal-to-noise ratio of >3 were observed at the expected retention times for $^{13}C$ labelled fatty acids FA 12:0 at 15.0 min, for FA 14:0 at 18.3 min, for FA 16:0 at 21.5 min, and for FA 18:0 at 24.5 min (indicated by dashed lines). For FA 8:0, FA 10:0 and the monitored unsaturated fatty acids, no signal that matched the expected retention times or that significantly differs from the background of a blank sample was observed. All retention times were confirmed with $^{12}C$ standards of the corresponding fatty acid.

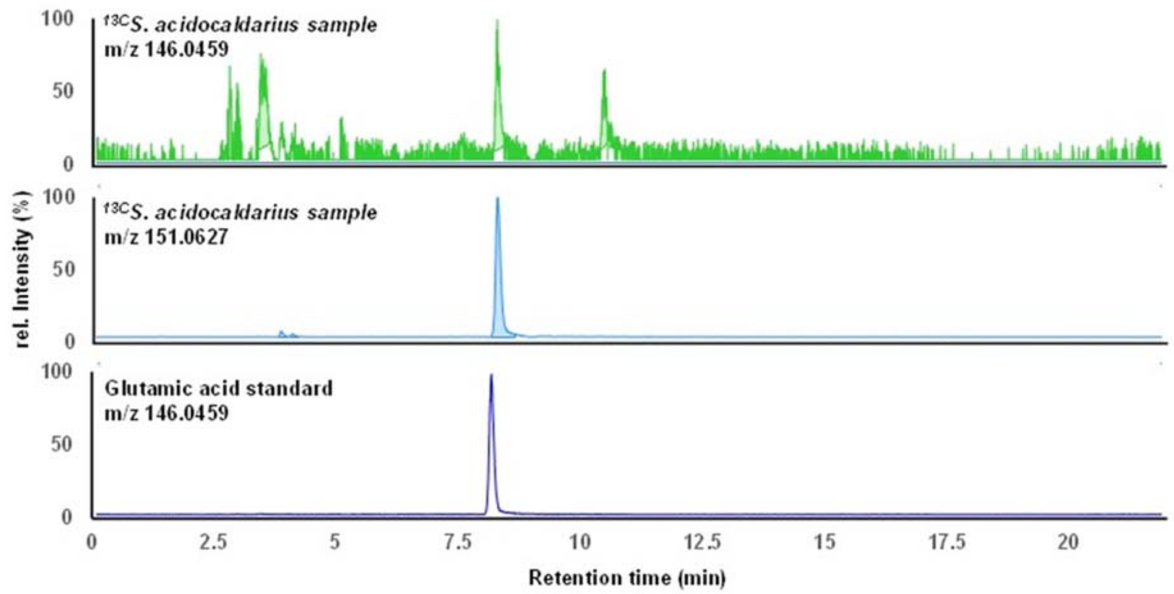


**Fig. S5. Extracted ion chromatograms (EICs) of the [M-H]$^-$ ion of glutamic acid obtained from LC-QTOF-MS analysis which confirm the incorporation of $^{13}C$ into metabolites of *S. acidocaldarius* wildtype MW00G.** Cells were grown for four passages on $^{13}C$-glycerol as sole carbon source. Top chromatogram shows the EIC of the mass-to-charge ratios (m/z) 146.0459 of glutamic acid for a sample grown on $^{13}C$-labled glycerol. Center chromatogram shows the EIC of the m/z 151.0627 for the same sample. This m/z corresponds to the [M-H]$^-$ ion of the fully labeled version glutamic acid. Bottom row shows the analysis of a glutamic acid standard to confirm the retention time of the compound. The results confirm a successful incorporation of $^{13}C$, other metabolites if present in the cells will be labelled as well.

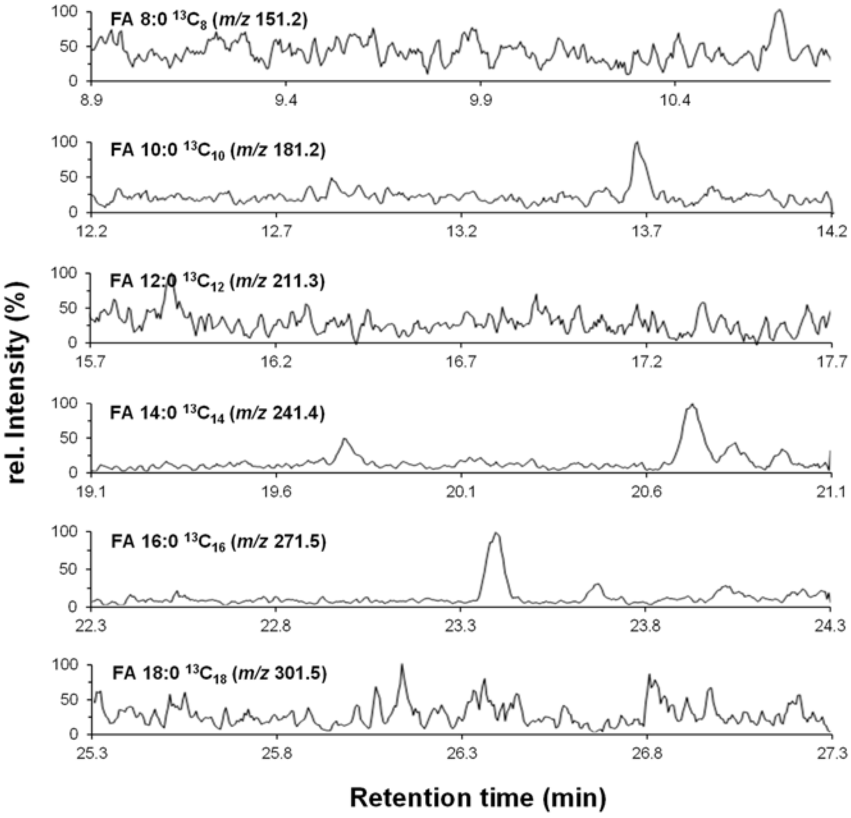


**Fig. S6. S6. GC-MS analysis of total fatty acid content in cells of *S. acidocaldarius* MW00G grown with $^{13}C$ D-xylose as sole carbon and energy source.** Cells were isolated, hydrolysed and derivatized as described in the methods section. The chromatograms represent an overview of the raw data and display the individual mass-to-charge ratios (m/z) for fully labelled $^{13}C$ fatty acids ranging from FA 8:0.to 18:0. No peaks were observed at the expected retention times of the $^{12}C$ variants of the corresponding fatty acids. In contrast to glycerol grown cells, the results show that upon growth on D-xylose no fatty acids are detectable in *S. acidocaldarius* wildtype MW00G.

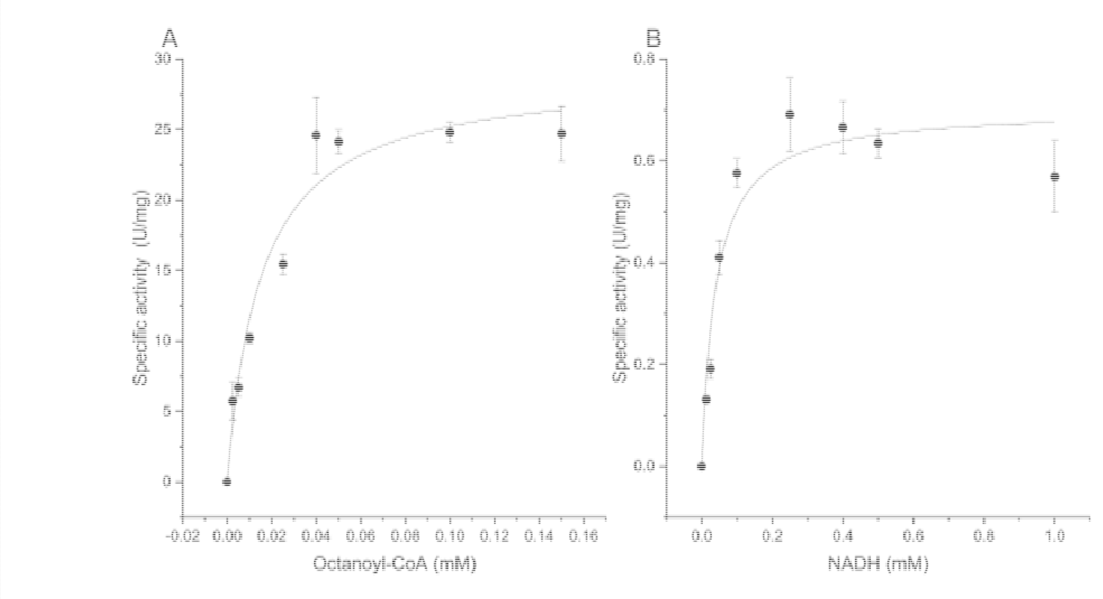


**Fig. S7. Investigation of kinetic properties of the recombinant acyl-CoA dehydrogenase (ACAD, Saci_1123) (A) and electron transfer flavoprotein (ETF, Saci_0315) (B) from *S. acidocaldarius*.** The kinetic properties of Saci_1123 were determined at 65°C in 50 mM HEPES-KOH (pH 6.5) with 20 mM KCl, 0.13 µg/µl ACAD, 1 mM $FcPF_6$ and 0-0.15 mM octanoyl-CoA (total volume 500 µl). Reduction of ferrocenium hexafluorophosphate ($FcPF_6$) was monitored at 300 nm (extinction coefficient 4.3 $mM^{-1}$ $cm^{-1}$). NADH-linked activity of ETF Saci_0315, was assayed in 50 mM HEPES-NaOH (pH 7.5) containing 100 mM NaCl, 0.2 mM iodonitrotetrazolium chloride (INT) and 0.015 µg/µl ETF protein in the presence of 0-1 mM NADH (total volume 500 µl). The activity was determined by monitoring the release of the red formazan at 500 nm (extinction coefficient 19.3 $mM^{-1}$ $cm^{-1}$). Independent measurements were performed in triplicate and error bars indicate the standard error of the mean (SEM).

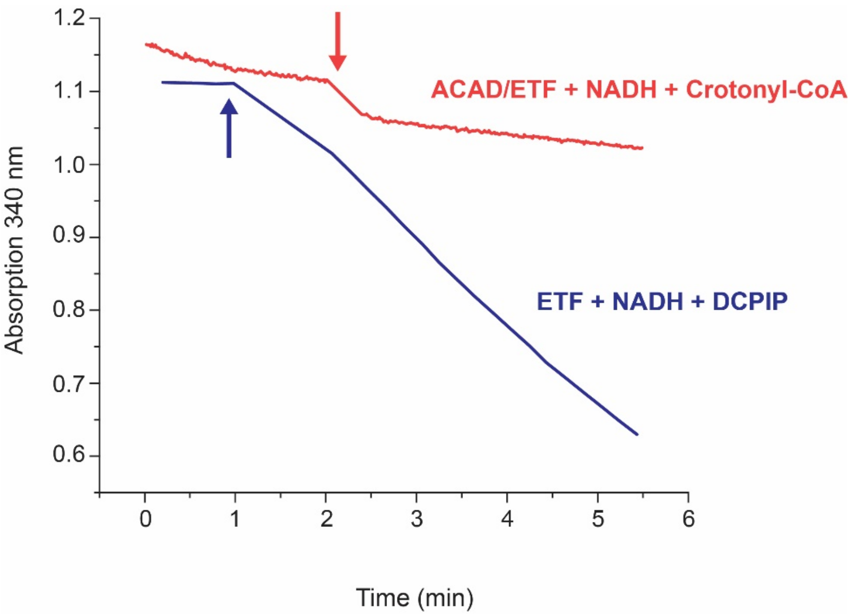


**Fig. S8. Electron transfer flavoprotein (ETF) Saci_0315 mediated electron transfer from NADH to 2,6-dichlorophenolindophenol (DCPIP) or via acyl-CoA dehydrogenase (ACAD, Saci_1123) to crotonyl-CoA**. Oxidation of NADH by ETF with DCPIP as electron acceptor (blue) was performed in 50 mM MES-KOH (pH 6.5) containing 20 mM KCl, and 0.0076 µg/µl ETF protein in the presence of 0.2 mM NADH and 0.2 mM DCPIP (total volume 500 µl). ETF mediated electron transfer from NADH via ACAD (0.0034 µg/µl) to crotonyl-CoA (0.4 mM) (red) was studied under the same conditions (without DCPIP). The data show that the ETF Saci_0315 accepts electrons from NADH and convey them to DCPIP but does not transfer the electrons to ACAD and further to crotonyl-CoA. Thus, the ACAD catalyzed reaction with ETF as electron carrier cannot be reverted with NADH as electron donor.

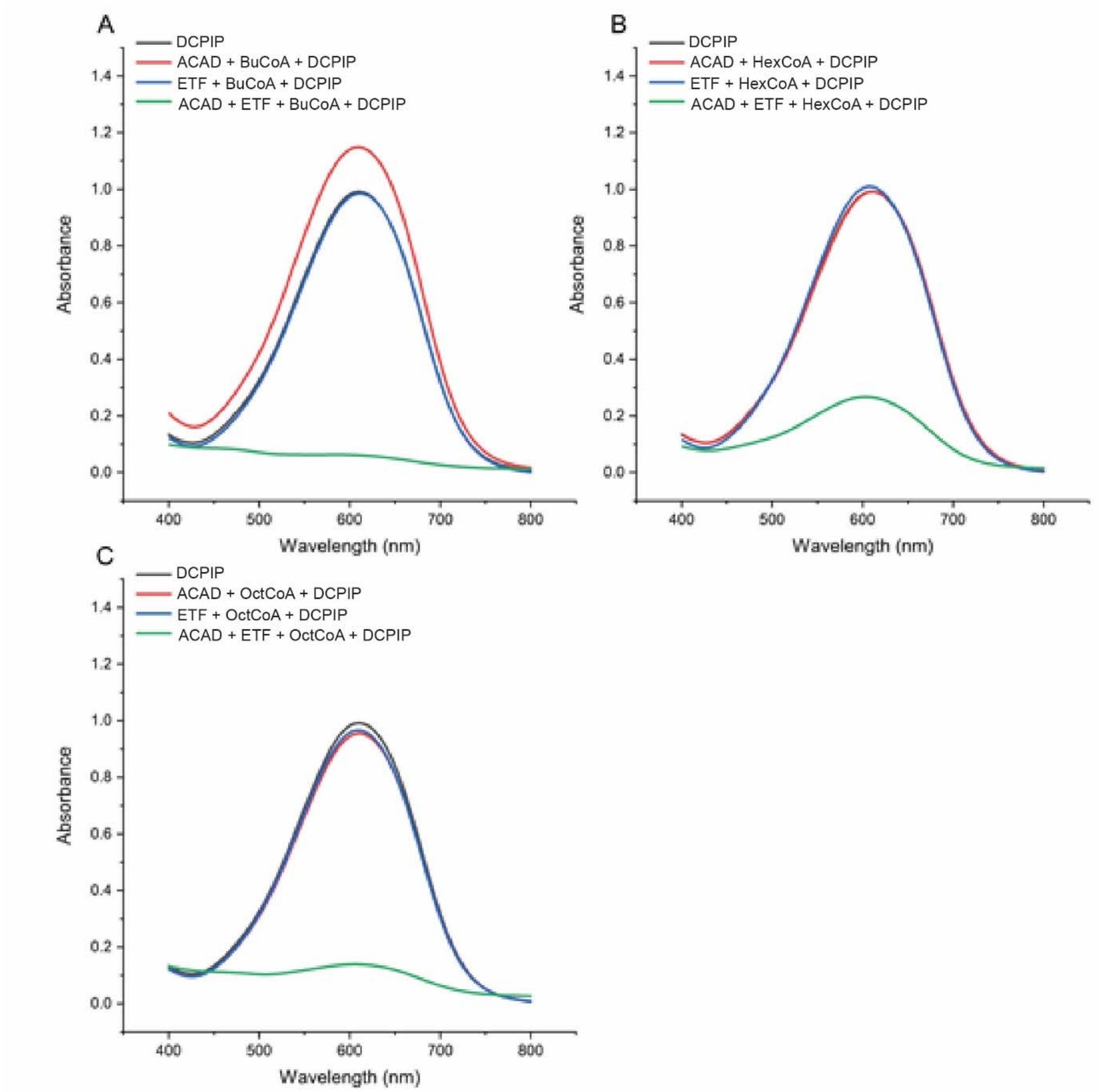


**Fig. S9. Investigation of electron transfer by acyl-CoA dehydrogenase (ACAD, Saci_1123) and electron transfer flavoprotein (ETF, Saci_0315).** The assay mixture (500 µl) contained 50 mM MES-KOH (pH 6.5), 20 mM KCl, 0.2 mM 2,6-dichlorophenolindophenol (DCPIP), 0.2 mM of different acyl-CoA (butyryl-CoA (BuCoA) (A), hexanoyl-CoA (HexCoA) (B) or octanoyl-CoA (OctCoA) (C)), 0.0034 µg/µl ACAD as well as 0.0076 µg/µl ETF and was incubated at 65°C for 5 min. Afterwards, the DCPIP spectrum in each sample was determined in the range of 400 to 800 nm. DCPIP exhibits the highest absorbance at 600 nm. Therefore, loss of absorbance at 600 nm indicated depletion of DCPIP due to the electron transfer from ACAD to DCPIP through ETF. The results indicate that all the employed acyl-CoAs require the presence of both ACDH and ETF for electron transport.

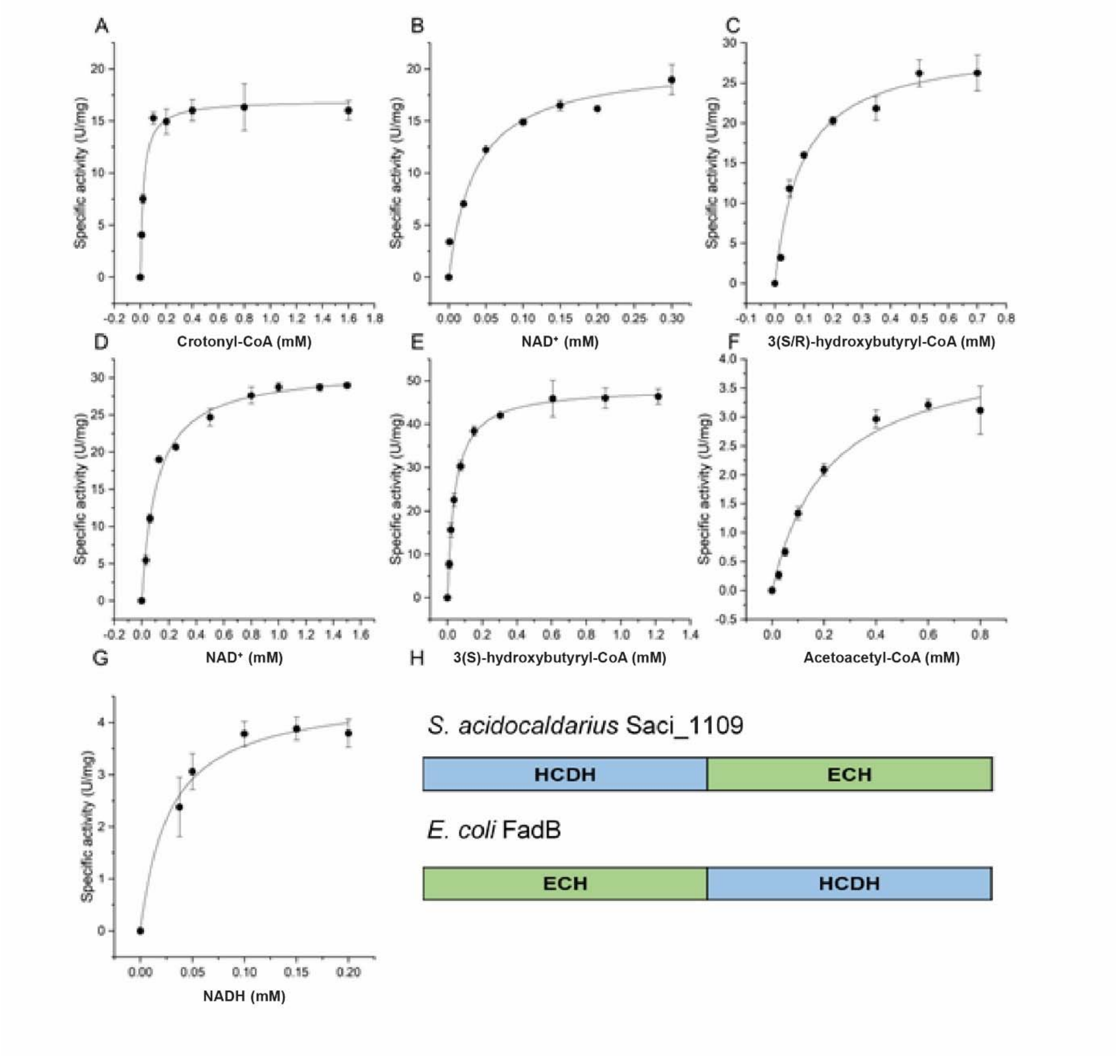


**Fig. S10. Investigation of kinetic properties of the recombinant 3(S)-hydroxyacyl-CoA dehydrogenase/enoyl-CoA hydratase (HCDH/ECH) fusion protein Saci_1109 from *S. acidocaldarius* and schematic illustration of the domain architecture.** The combined activity of the bifunctional HCDH/ECH Saci_1109 was determined in the oxidative direction at 75°C in 100 mM TRIS-HCl (pH 7) with 0.4 mM crotonyl-CoA, 0.2 mM $NAD^+$ and 0.00138 µg/µl protein (total volume 500 µl) and the formation of NADH was monitored at 340 nm. For the determination of 3(R)-hydroxybutyryl-CoA (3(R)-HBCoA) oxidizing partial reaction catalyzed by the HCDH domain a racemic mixture of 3(S/R)-HBCoA or the pure stereoisomer 3(S)-HBCoA was applied instead of crotonyl-CoA. The kinetic parameters for crotonyl-CoA (A), $NAD^+$ (B, with crotonyl-CoA), 3(S/R)-HBCoA (C), $NAD^+$ (D, with 3(S/R)HBCoA), and 3(S)-HBCoA (E) were determined. For detection of the activity in the reductive direction at 35°C, 0.6 mM acetoacetyl-CoA, 0.2 mM NADPH, and 0.0081 µg/µl protein was used. The kinetic parameters for acetoacetyl-CoA (F) and NADH (G) were determined. Independent measurements were performed in triplicate and error bars indicate the standard error of the mean (SEM). In (H) the domain organization of the Saci_1109 HCDH/ECH in comparison to the *E. coli* FadB protein is depicted.

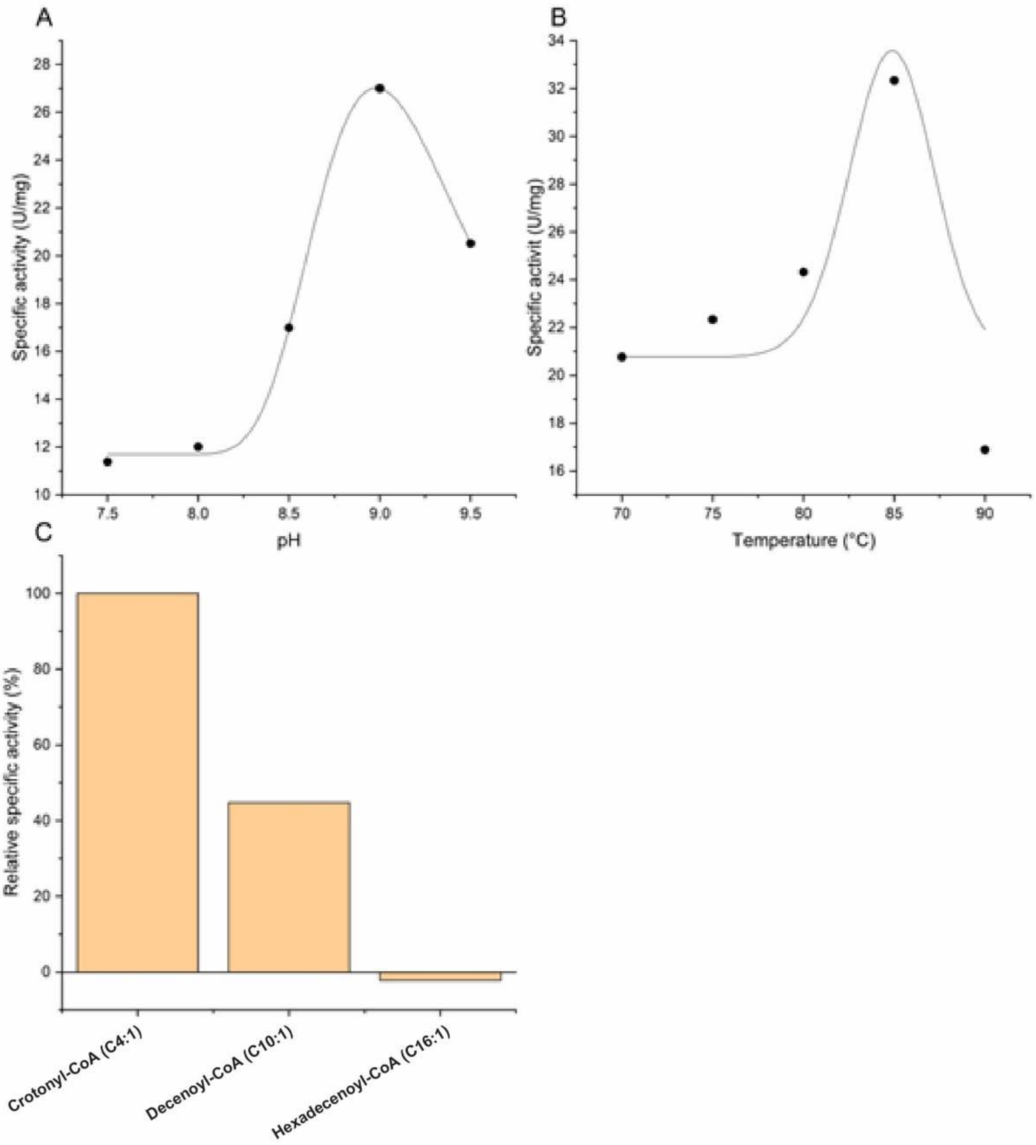


**Fig. S11. Investigation of the pH (A) and temperature (B) optimum and the substrate specificity (C) of the recombinant 3(S)-hydroxyacyl-CoA dehydrogenase/enoyl-CoA hydratase fusion protein (HCDH/ECH, Saci_1109) from *S. acidocaldarius*.** The optimal pH was determined in a mixed buffer of 50 mM MES, 50 mM HEPES and 50 mM TRIS at 70°C (total volume 500 µl). The assay contained 0.4 mM crotonyl-CoA, 0.2 mM $NAD^+$ and 0.00138 µg/µl Saci_1109. The temperature optimum was determined in 100 mM HEPES-NaOH (pH 8) utilizing the same assay. The substrate specificity was performed in presence of 0.3 mM of crotonyl-CoA, decenoyl-CoA and hexadecenoyl-CoA. The assay (400 µl) was done at 75°C, pH 7 with 0.2 mM $NAD^+$ and 0.0009 µg/µl protein.

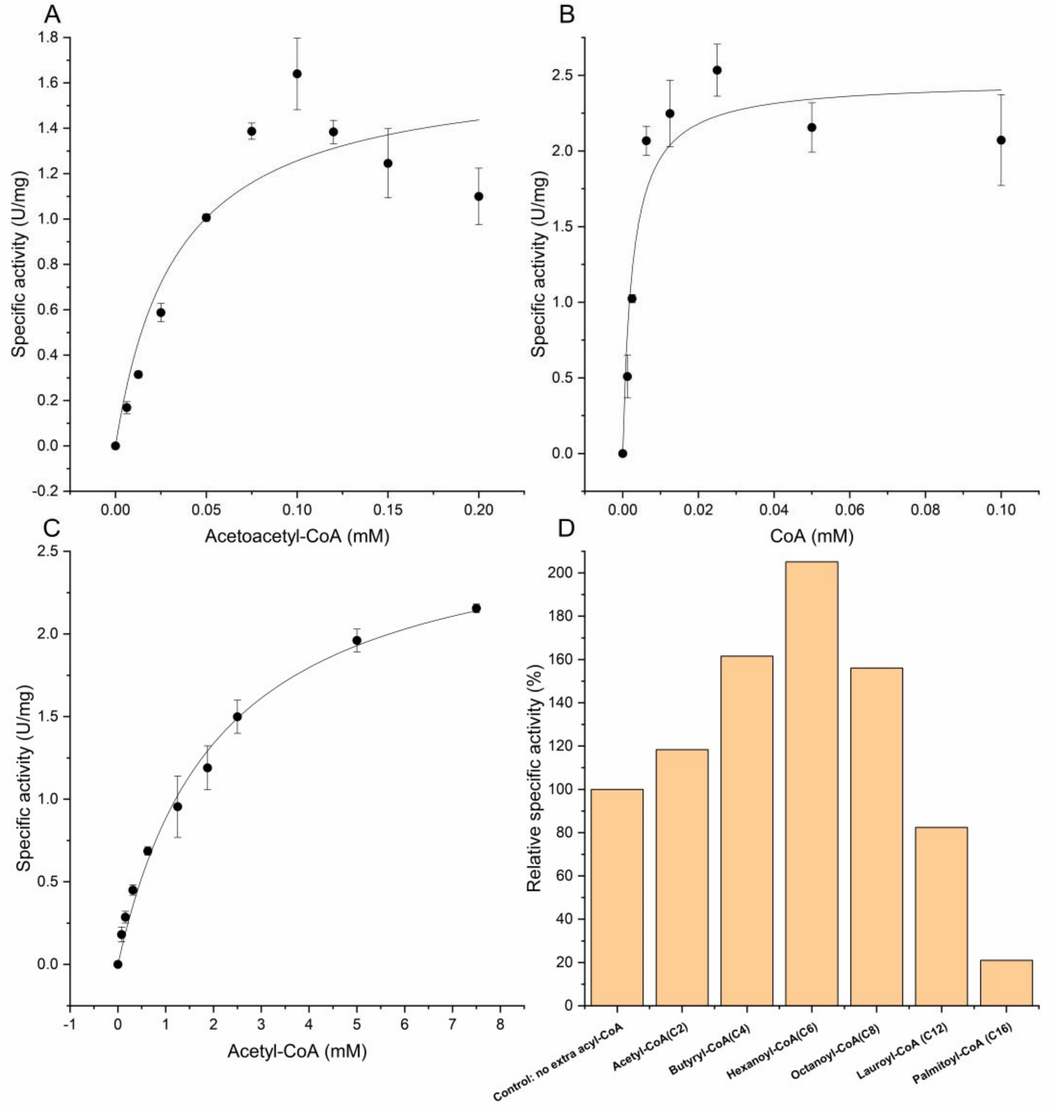


**Fig. S12. Investigation of kinetic properties and substrate specificity of the recombinant ketothiolase (KT, Saci_1114) from *S. acidocaldarius*.** The specific activity of KT was determined photometrically at room temperature (23°C), by monitoring the decrease of $Mg^{2+}$-acetoacetyl-CoA (AcAcCoA) chelation complex (extinction coefficient of 21.4 $mM^{-1}$ $cm^{-1}$) at 303 nm. The reaction mixture contained 100 mM TRIS-HCl (pH 8), 20 mM $MgCl_2$, 0.2 mM CoA, 0.1 mM AcAcCoA and 0.0054 µg/µl Saci_1114 (total volume 500 µl). To determine the kinetic properties variable concentrations of AcAcCoA (A) or Coenzyme A (CoA) (B) were employed. The activity in the direction of the Claisen condensation of two acetyl-CoAs was determined in a coupled assay with HCDH/ECH Saci_1109 as auxiliary enzyme by monitoring NADH oxidation at 340 nm. The enzyme assay (total volume 500 µl) was performed in 100 mM MOPS-NaOH (pH 6.5 at 75°C), 0.3 mM NADH, 0.0342 µg/µl Saci_1109, 0.0216 µg/µl Saci_1114 and 0-7.5 mM acetyl-CoA (C). Independent measurements were performed in triplicate and error bars indicate the standard error of the mean (SEM). The substrate specificity (D) was determined in presence of 2.5 mM acetyl-CoA and 0.5 mM of the indicated acyl-CoAs with different chain lengths, i.e. acetyl-CoA (C2), butyryl-CoA (C4), hexanoyl-CoA (C6), octanoyl-CoA (C8), lauroyl-CoA (C12) and palmitoyl-CoA (C16).

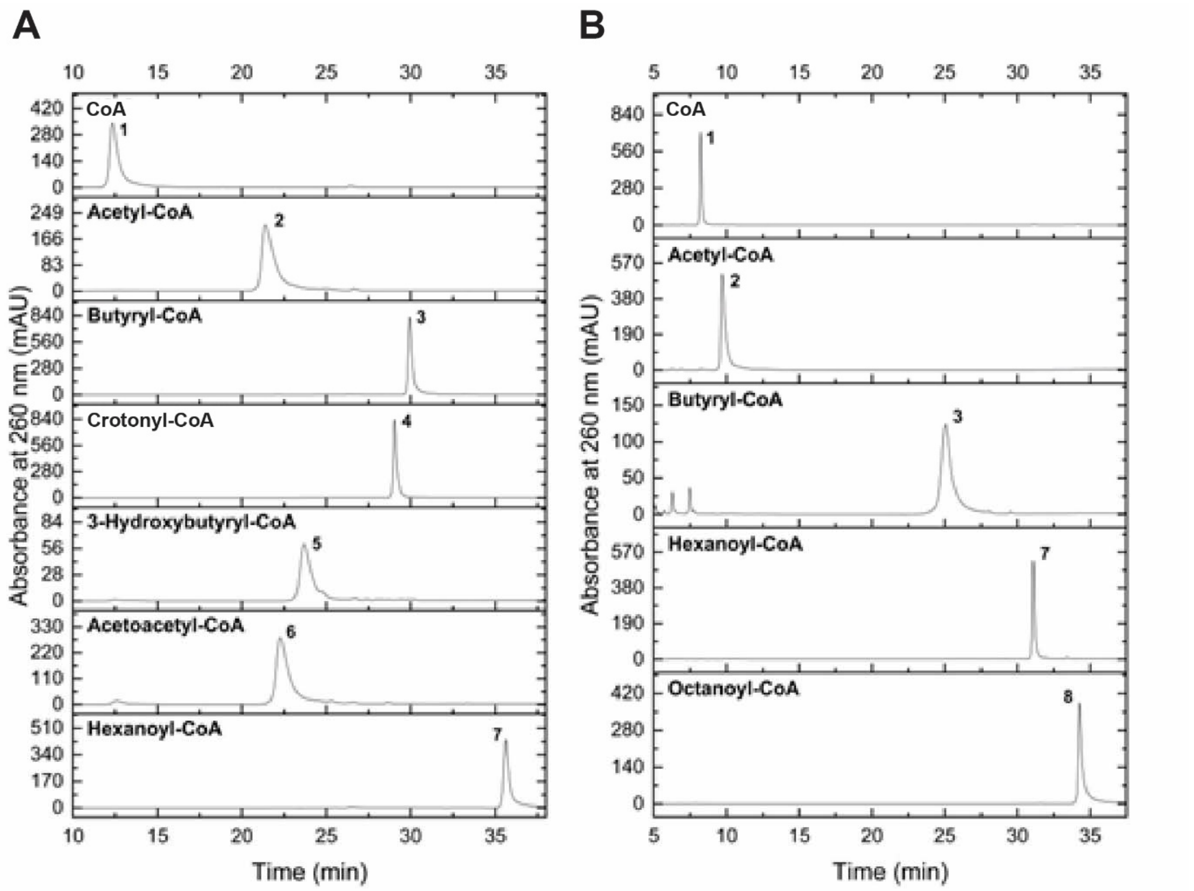


**Fig. S13. HPLC chromatograms of CoA ester standards involved in fatty acid metabolism.** Two distinct programs with different acetonitrile (ACN) concentration gradients were established for analyzing different chain lengths of CoA esters. The HPLC program “4-30% ACN program” (A) was used for shorter chain acyl-CoAs, and the “1-60% ACN program” (B) for longer chain CoA esters. The retention times representing the relevant CoA compounds are shown in Table S5. The peak numbers correspond to the following CoA compounds: 1, CoA (Coenzym A); 2, AcetylCoA; 3, Butyryl-CoA; 4, Crotonyl-CoA; 5, 3-Hydroxybutyryl-CoA; 6, Acetoacetyl-CoA; 7, Hexanoyl-CoA; 8, Octanoyl-CoA.

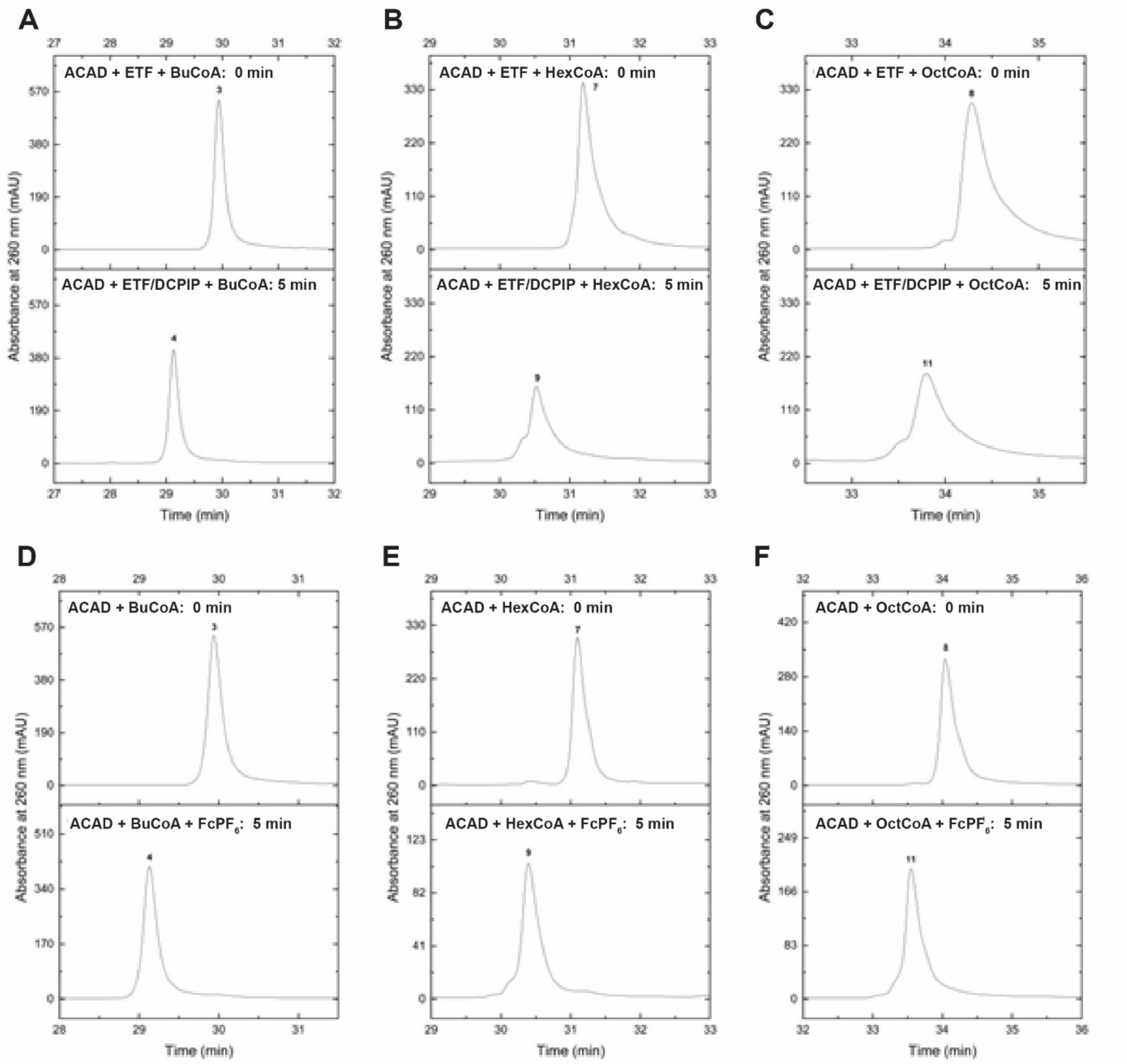


**Figure S14. HPLC analysis of acyl-CoA oxidation into enoyl-CoA by the recombinant acyl-CoA dehydrogenase (ACAD, Saci_1123) in β oxidation.** The oxidation of saturated acyl-CoAs to the corresponding enoyl-CoAs was analyzed in a discontinuous assay (total volume 50 µl) at 65°C. 0.02 µg/µl ACAD (Saci_1123) and 0.01 µg/µl electron transfer flavoprotein (ETF, Saci_0315) were incubated in 50 mM MES-KOH (pH 6.5) with 20 mM KCl, 0.4 mM 2,6-dichlorophenolindophenol (DCPIP) and 0.4 mM of acyl-CoAs (i.e. butyryl-CoA (BuCoA), hexanoyl-CoA (HexCoA) or octanoyl-CoA (OctCoA)) for 5 minutes (A, B & C). In addition, 0.8 mM ferrocenium hexafluorophosphate ($FcPF_6$) was used as artificial electron acceptor instead of ETF and DCPIP (D, E & F). Butyryl-CoA (BuCoA) conversion was analyzed via the 4-30% ACN HPLC program (A, D) and oxidation of hexanoyl-CoA (HexCoA) (B, E) or octanoyl-CoA (OctCoA) (C, F) by the 1-60% ACN HPLC program. The upper chromatogram in each panel shows the time point 0 min before reaction start by addition of the electron acceptor. As shown, all the tested acyl-CoAs (peak 3, 7 or 8) were fully converted into the corresponding enoyl-CoA products (peak 4, 9 or 11). For assignment of peak numbers to compounds and HPLC programs see Fig. S13.

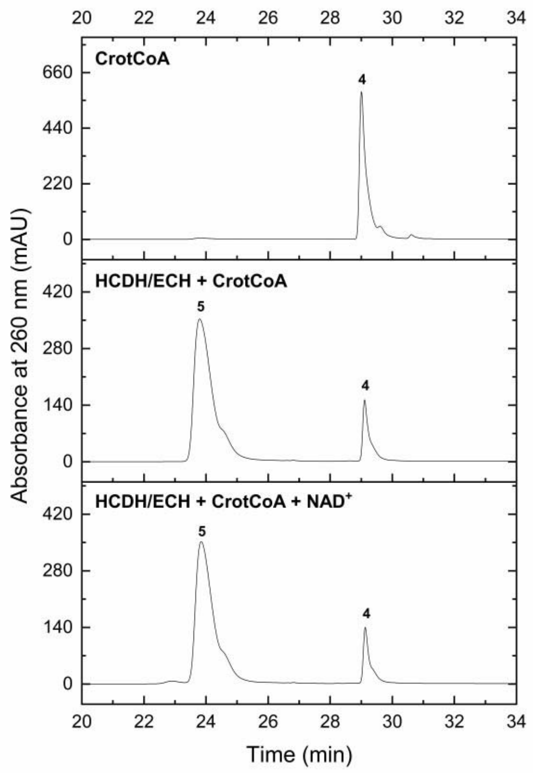


**Fig. S15. HPLC analysis of crotonyl-CoA (CrotCoA) conversion by the bifunctional 3(S)-hydroxyacyl-CoA dehydrogenase/enoyl-CoA hydratase (HCDH/ECH, Saci_1109) from *S. acidocaldarius* in β oxidation.** The discontinuous assay (total volume 50 µl) was performed in 50 mM HEPES-NaOH (pH 6.5) with 20 mM KCl, 0.4 mM crotonyl-CoA (CrotCoA) and 0.0144 µg/µl Saci_1109 in absence (middle) or presence of 2 mM $NAD^+$ (bottom). The control without enzyme and $NAD^+$ (i.e. before start of the reaction with enzyme) is shown in the upper chromatogram. The reaction mixture was incubated at 65°C for 15 min and the samples were then analyzed via the 1-60% ACN HPLC program. The formation of 3-hydroxybutyryl-CoA (peak 5) from crotonyl-CoA (peak 4) was detected (middle). However, further production of acetoacetyl-CoA was not observed after addition of $NAD^+$. For assignment of peak numbers to compounds and HPLC programs see Fig. S13.

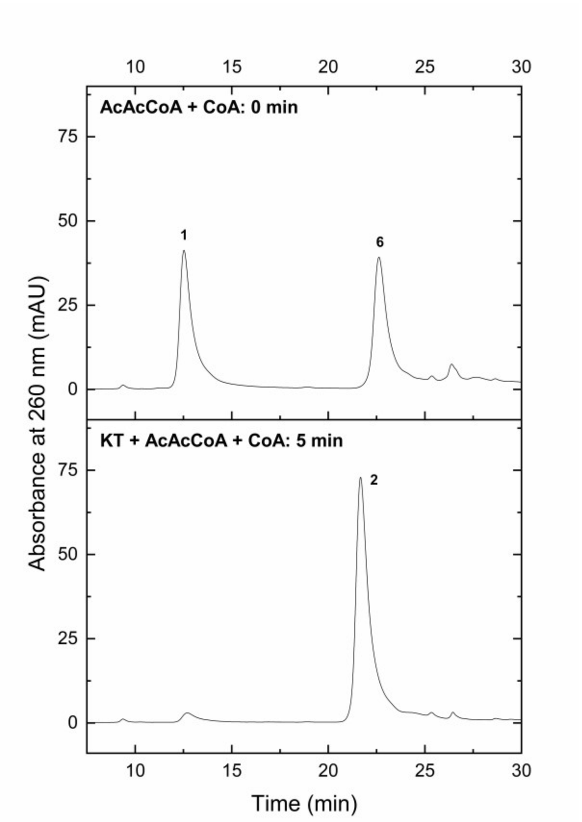


**Figure S16. HPLC analysis of cleavage of acetoacetyl-CoA (AcAcCoA) by the recombinant β-ketothiolase (KT, Saci_1114) in β oxidation.** To analyze the last thiolytic step in β oxidation, the assay (total volume 100 µl) was performed in 50 mM MES-KOH (pH 6.5) with 0.1 mM Coenzyme A (CoA) and 0.1 mM AcAcCoA. The reaction mixture was preincubated at 23°C for 2 min and the reaction was then initiated by addition of 0.027 µg/µl of KT followed by incubation for 5 min. Samples before addition of KT (0 min, upper chromatogram) and after 5 min of incubation with KT (bottom chromatogram), were investigated by the 4-30% ACN HPLC program. As shown, AcAcCoA (peak 6) and CoA (peak 1) were completely converted to acetyl-CoA (peak 2). For assignment of peak numbers to compounds and HPLC programs see Fig. S13.

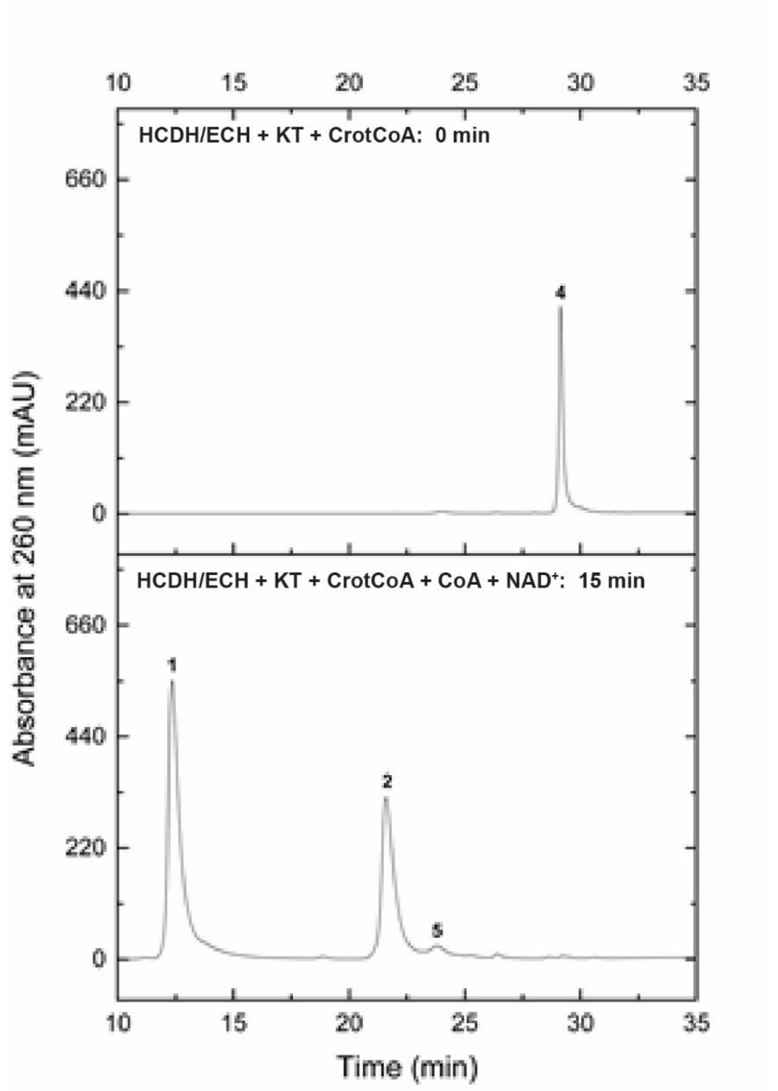


**Fig. S17. HPLC analysis of the last three steps in β oxidation catalyzed by the recombinant bifunctional 3(S)-hydroxyacyl-CoA dehydrogenase/enoyl-CoA hydratase (HCDH/ECH, Saci_1109) and β-ketothiolase (KT, Saci_1114).** The assay (total volume 50 µl) was performed in 50 mM MES-NaOH (pH 6.5) at 65°C, containing 20 mM KCl, 0.4 mM crotonyl-CoA (CrotCoA), 2 mM $NAD^+$, 0.0144 µg/µl HCDH/ECH, 1.6 mM CoA and 0.054 µg/µl KT. The reaction was started by addition of $NAD^+$ and CoA, incubated for 15 min and analyzed by HPLC using the 4-30% ACN program. After 15 min the initial substrate crotonyl-CoA (peak 4, upper chromatogram, 0 min) was completely converted to the end product acetyl-CoA (peak 2) by HCDH/ECH and KT in presence of CoA (peak 1). In addition, a limited amount of the intermediate 3-hydroxybutyryl-CoA (peak 5) was detected. In the control before addition of $NAD^+$ and CoA (upper chromatogram) no crotonyl-CoA conversion was observed. For assignment of peak numbers to compounds and HPLC programs see Fig. S13.

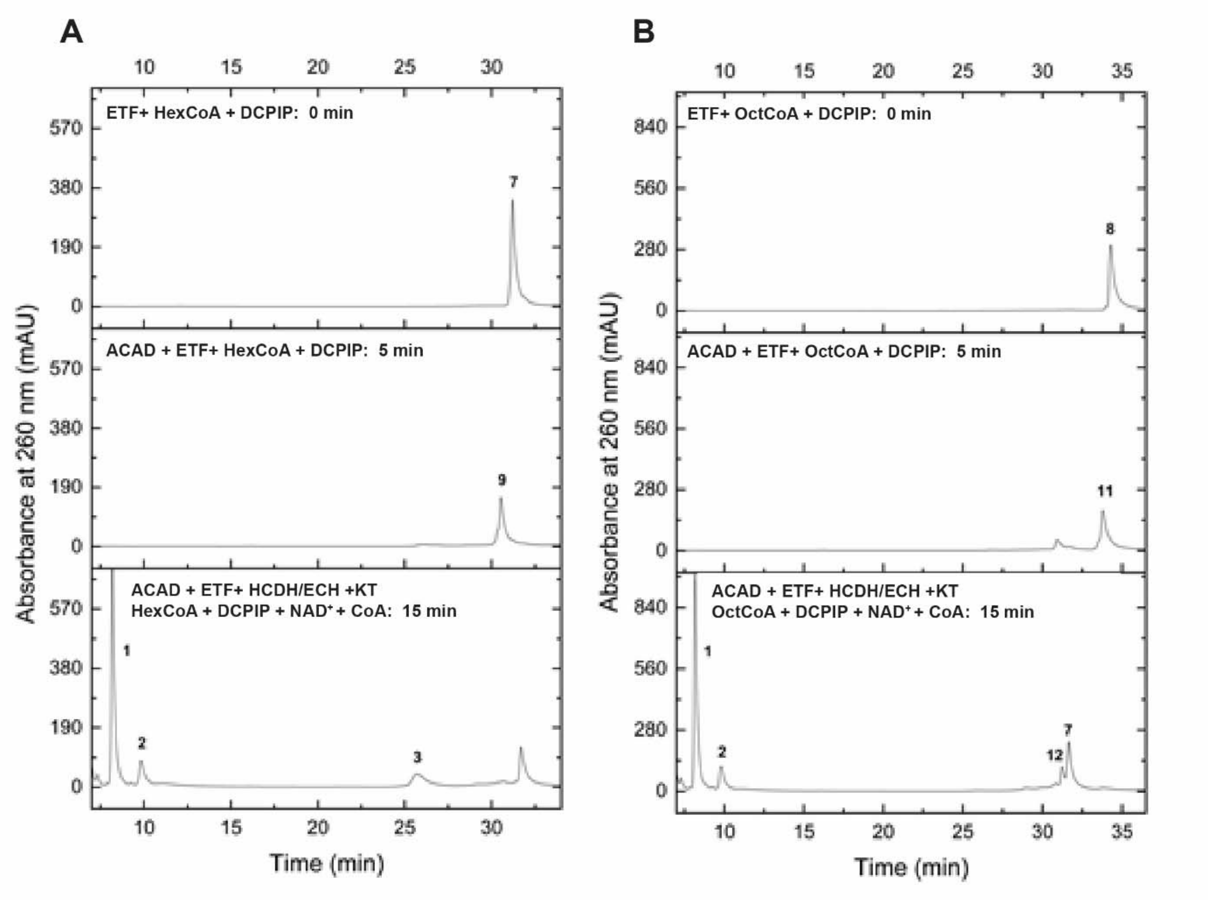


**Figure S18. HPLC analysis of the complete β oxidation enzyme cascades for degradation of hexanoyl-CoA (HexCoA) (A) and octanoyl-CoA (OctCoA) (B).** The enzyme cascade reactions (total volume 50 µl) were carried out in two steps. The first oxidation step by 0.02 µg/µl acyl-CoA dehydrogenase (ACAD, saci_1123) and 0.01 µg/µl electron transfer flavoprotein (ETF, saci_0315) was done in 50 mM MES-KOH (pH 6.5 at 65°C) with 20 mM KCl, 0.4 mM 2,6-dichlorophenolindophenol (DCPIP) and 0.4 mM of acyl-CoA (i.e. hexanoyl-CoA (HexCoA) (A) or octanoyl-CoA (OctCoA) (B)). The reaction was run for 5 min. In the second step, 0.2 mM $NAD^+$, 0.0144 µg/µl bifunctional 3(S)-hydroxyacyl-CoA dehydrogenase/enoyl-CoA hydratase (HCDH/ECH, Saci_1109), 1.6 mM CoA and 0.054 µg/µl β-ketothiolase (KT, Saci_1114) were added to the reaction mixture and incubated for 15 min. The β oxidation metabolites were detected by the 1-60% ACN HPLC program. The control before reaction start with ACAD is shown in the upper chromatograms. After the first step (addition of ACAD), acyl-CoAs (peak 7 or 8) were completely oxidized to the corresponding enoyl-CoAs, i.e. hexenoyl-CoA ((A), peak 9, middle panel) and octenoyl-CoA ((B), peak 11, middle panel) by ACAD and ETF in presence of the artificial electron acceptor DCPIP. Finally, after addition of HCDH/ECH, KT, CoA (peak 1) and $NAD^+$ (not shown), hexenoyl-CoA could be fully converted to acetyl-CoA (peak 2) and butyryl-CoA (peak 3) ((A), bottom) whereas the octenoyl-CoA was completely oxidized to acetyl-CoA and hexanoyl-CoA (peak 7) as well as small amounts of 3-hydroxyoctanoyl-CoA (peak 12) ((B), bottom panel). For assignment of peak numbers to compounds and HPLC programs see Fig. S13.

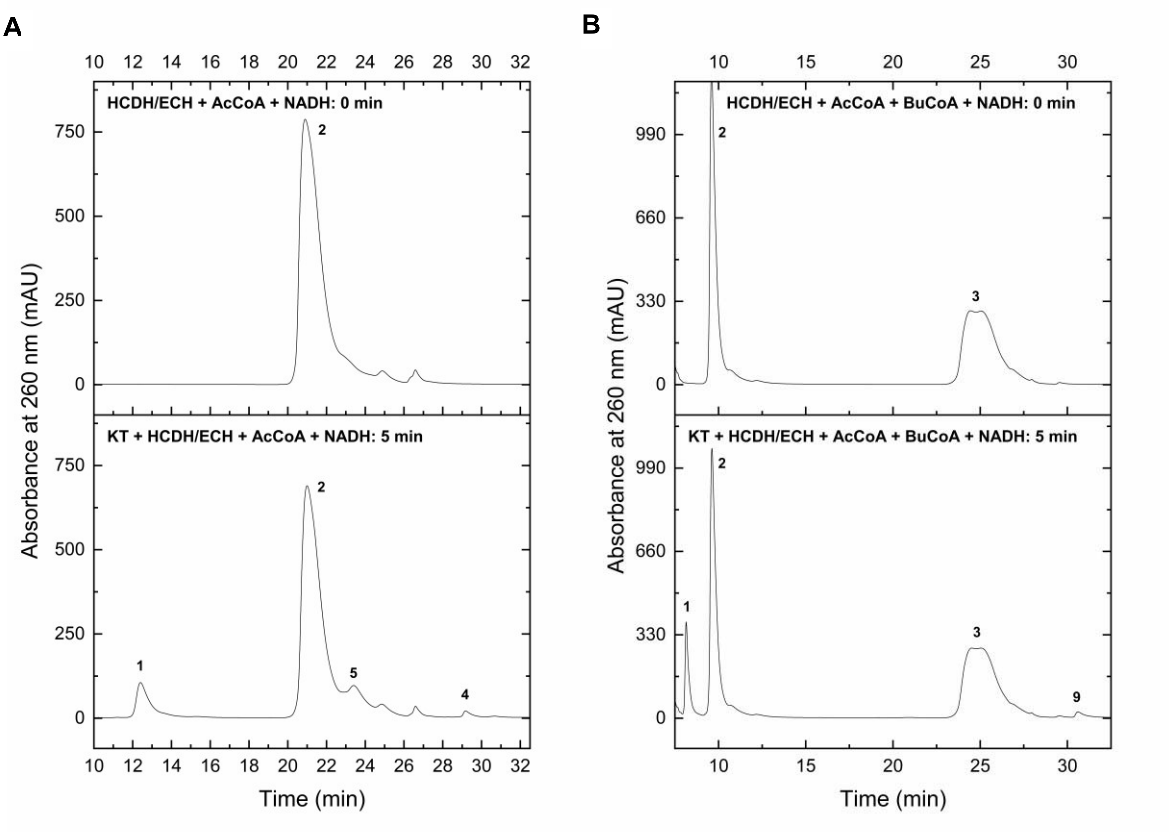


**Fig. S19. HPLC analysis of the acetyl-CoA (AcCoA) (A) and butyryl-CoA (BuCoA) (B) conversion in the reductive direction catalyzed by recombinant β-ketothiolase (KT, Saci_1114) and bifunctional 3(S)-hydroxyacyl-CoA dehydrogenase/enoyl-CoA hydratase (HCDH/ECH, Saci_1109).** The enzyme assay (total volume 400 µl) was performed in 50 mM HEPES-NaOH (pH 6.5), 2 mM acetyl-CoA (AcCoA), 0.3 mM NADH and 0.04275 µg/µl HCDH/ECH (Saci_1109) at 65°C. After preincubation for 2 min, the reaction was started by addition of 0.0405 µg/µl KT (Saci_1114) and run for 5 min. Controls before addition of KT (0 min) are shown in the upper chromatograms in (A) and (B). After incubation, samples were analyzed by the 4-30% ACN (A) and 1-60% ACN (B) HPLC program. In both measurements, release of free CoA (peak 1) by KT was observed. During acetyl-CoA conversion (A), a small amount of crotonyl-CoA (peak 4) and 3-hydroxybutyryl-CoA (peak 5) was formed. Even less hexenoyl-CoA (peak 9) was synthesized from butyryl-CoA (peak 3) and acetyl-CoA (peak 2) by KT and HCDH/ECH (B). This is in accordance with the thermodynamics of the reaction sequence ($\Delta G^{0'}$+14 kJ $mol^{-1}$) (see "Supplementary text"). For assignment of peak numbers to compounds and HPLC programs see Fig. S6.

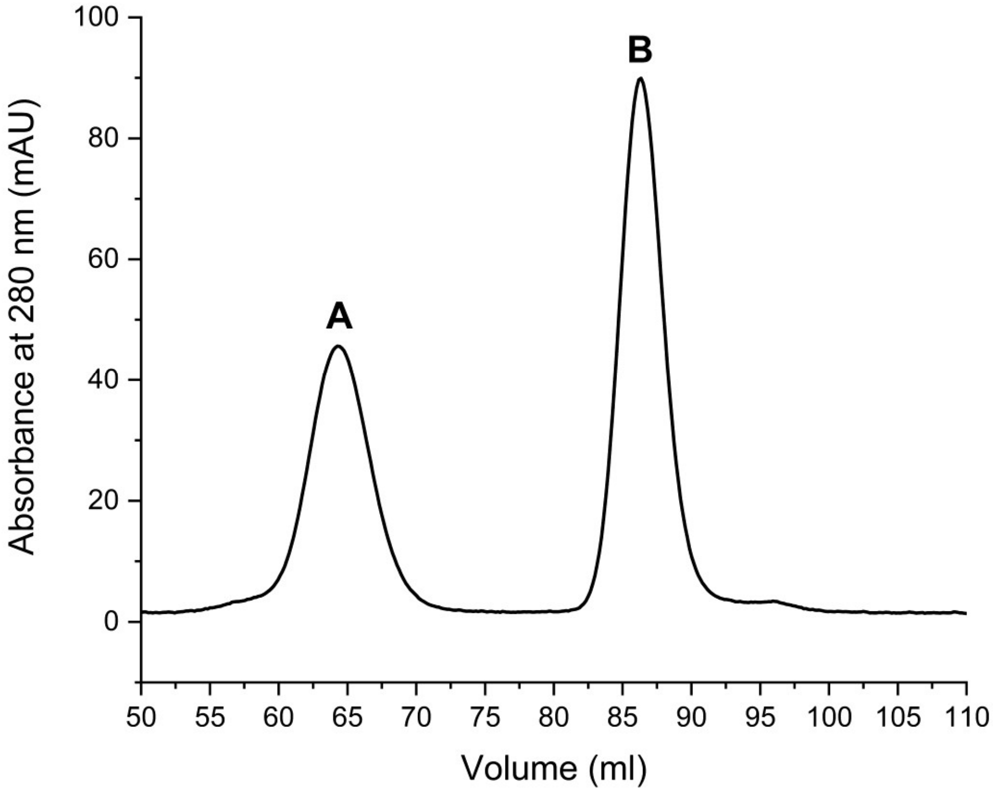


**Fig. S20. Complex formation of the recombinant bifunctional 3(S)-hydroxyacyl-CoA dehydrogenase/enoyl-CoA hydratase (HCDH/ECH, Saci_1109) and β-ketothiolase (KT, Saci_1114) from *S. acidocaldarius* analyzed by size exclusion chromatography.** 1090 µg of purified recombinant ECH/HCDH and 644 µg of purified recombinant KT, were mixed and incubated on ice for 4 hours. Then, the protein solution was applied onto a Superdex 200 prep grad HiLoad 16/60 gel filtration column (GE Healthcare Life Sciences, Freiburg, Germany) equilibrated in 50 mM HEPES-NaOH (pH 7.2) and 300 mM NaCl and separated. Two distinct peaks representing either HCDH/ECH (A, 64.33 ml) or KT (B, 86.29 ml) were obtained indicating no complex formation between these two proteins under the chosen experimental conditions.

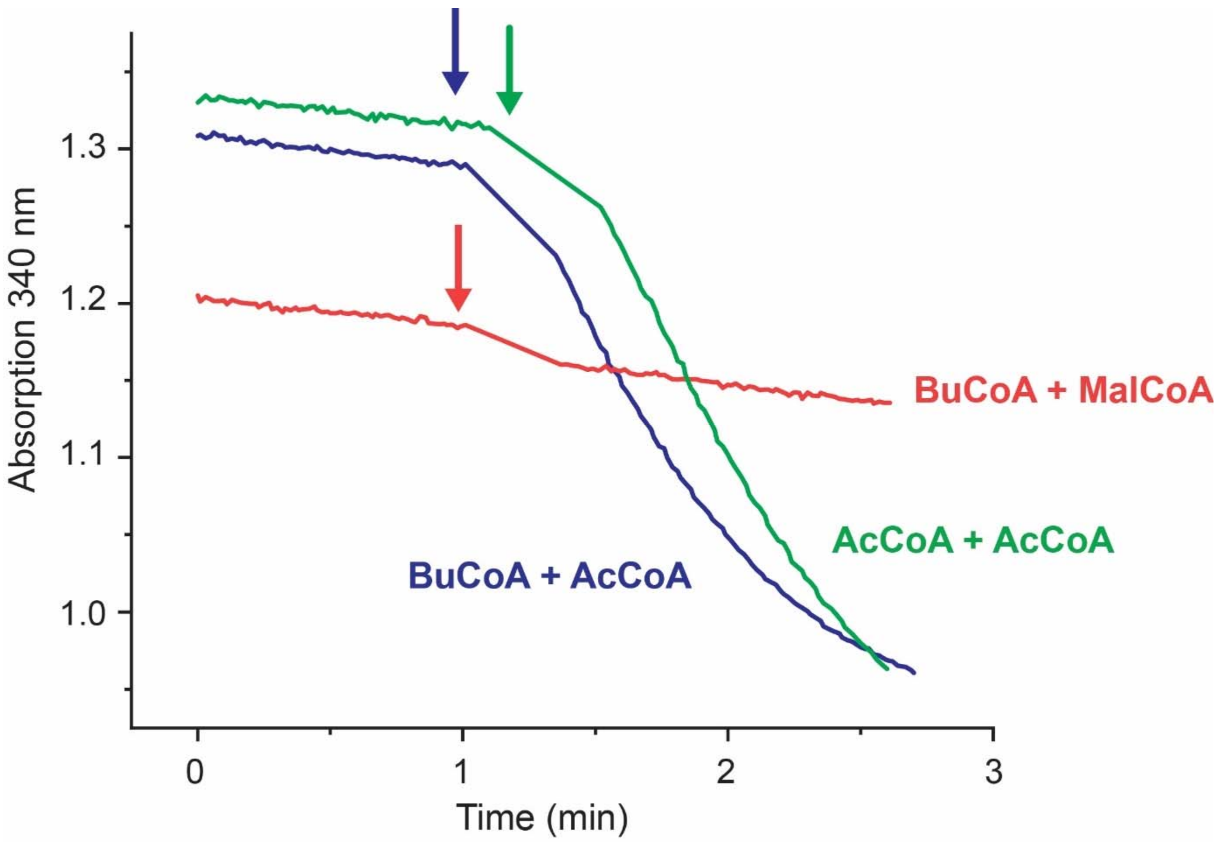


**Fig. S21. The recombinant β-ketothiolase (KT, Saci_1114) does not utilize malonyl-CoA as extender unit.** This was studied by incubation of 0.0054 µg/µl of Saci_1114 under the same conditions as described for the KT activity measurement (see methods section) by coupling acetoacetyl-CoA (AcAcCoA) and ketohexanoylyl-CoA formation to NADH (0.2 mM) oxidation via 0.0342 µg/µl bifunctional 3(S)-hydroxyacyl-CoA dehydrogenase/enoyl-CoA hydratase (HCDH/ECH, Saci_1109). As substrates either 2 mM of acetyl-CoA (AcCoA, green), 1 mM butyryl-CoA (BuCoA) + 1 mM AcCoA (blue), or 1 mM BuCoA + 1 mM malonyl-CoA (MalCoA) (red) were used and absorption changes over time were followed at 340 nm. The NADH oxidation with AcCoA (green) and BuCoA+AcCoA (blue) demonstrates ketoacyl-CoA formation and thus Ac-CoA as extender unit, whereas with Mal-CoA no ketoacyl-CoA is formed indicating that Mal-CoA cannot be utilized as extender unit by KT.

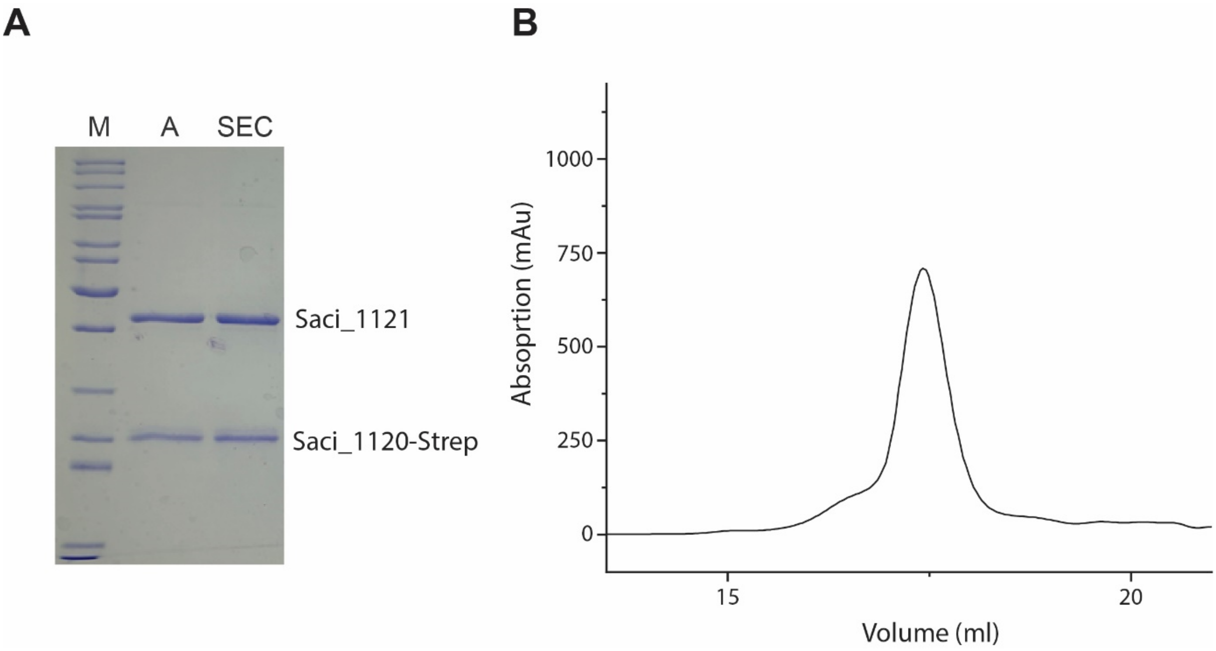


**Fig. S22. The DUF35 domain protein (DUF35, Saci_1120) and the β-ketothiolase (KT, Saci_1121) form a complex after co-expression in *S. acidocaldarius***. The genes encoding Saci_1121 and Saci_1120 form an operon in *S. acidocaldarius*. Both genes were co-expressed in *S. acidocaldarius* with a C-terminal twin-strep-tag introduced to the DUF35 domain protein. Protein purification was performed via strep-tactin affinity chromatography (AC) and size exclusion chromatography (SEC). In both purification steps DUF35 and KT co-eluted as shown by SDS-PAGE (A) and by one single peak in the chromatogram after SEC Abbreviation: M: Marker (PageRuler™ Unstained Protein Ladder, Thermo Fisher Scientific, USA).

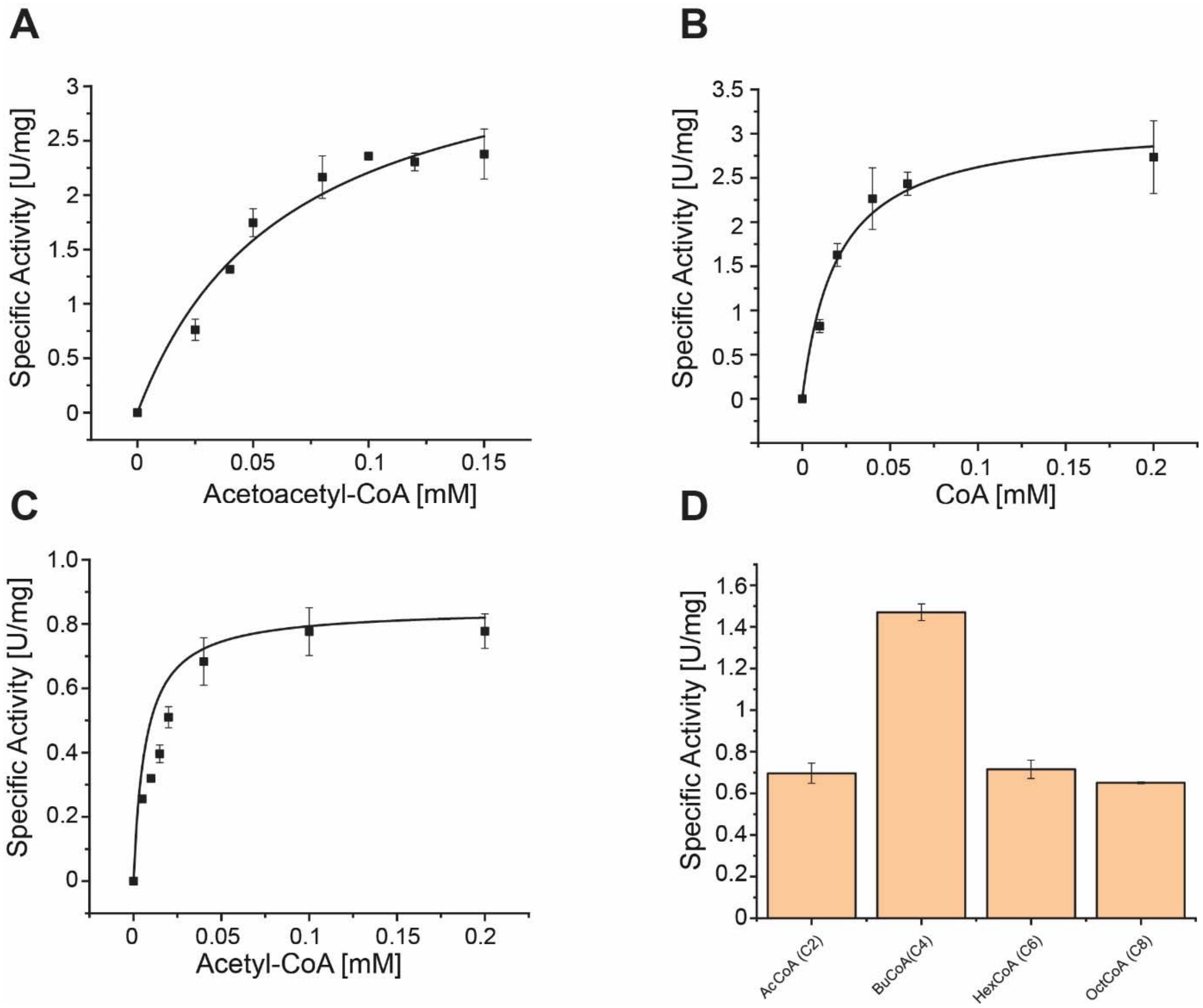


**Fig. S23. Investigation of kinetic properties and substrate specificity of the recombinant β-ketothiolase Saci_1121/DUF35 domain protein Saci_1120 (KT/DUF35 (Saci_1121/1120)) complex from *S. acidocaldarius*.** The specific activity of the Saci_1121/1120 KT/DUF35 subcomplex was photometrically tested at room temperature (23°C) and pH 8.0 by monitoring the decrease of $Mg^{2+}$-acetoacetyl-CoA (AcAcCoA) chelation complex (extinction coefficient of 21.4 $mM^{-1}$ $cm^{-1}$) at 303 nm. The reaction mixture (total volume 500 µl) contained 100 mM TRIS-HCl (pH 8), 20 mM $MgCl_2$, and 0.0036 µg/µl Saci_1121/Saci_1120. To determine kinetic properties, variable concentrations of AcAcCoA (0-0.2 mM) with 0.2 mM CoA (A) or CoA (0-0.1 mM) with 0.1 mM AcAcCoA (B) were employed. The activity in the direction of the Claisen condensation of two acetyl-CoA was determined by coupling the AcAcCoA formation to NADH oxidation via 3(S)-hydroxyacyl-CoA dehydrogenase/enoyl-CoA hydratase (HCDH/ECH, Saci_1109) as auxiliary enzyme at 340 nm. The enzyme assay (total volume 500 µl) included 100 mM MOPS-NaOH (pH 6.5 at 75°C), 0.2 mM NADH, 0.02052 µg/µl Saci_1109, 0.0104 µg/µl Saci_1121/Saci_1120 and 0-0.2 mM acetyl-CoA for $K_M$ measurement (C). The independent measurements were performed in triplicate and error bars indicate the standard error of the mean (SEM). The substrate specificity (D) was determined by including 1 mM acetyl-CoA and 0.5 mM of extra acyl-CoA with different chain lengths (acetyl-CoA (AcCoA, C2), butyryl-CoA (BuCoA, C4), hexanoyl-CoA (HexCoA, C6), octanoyl-CoA (OctCoA, C8).

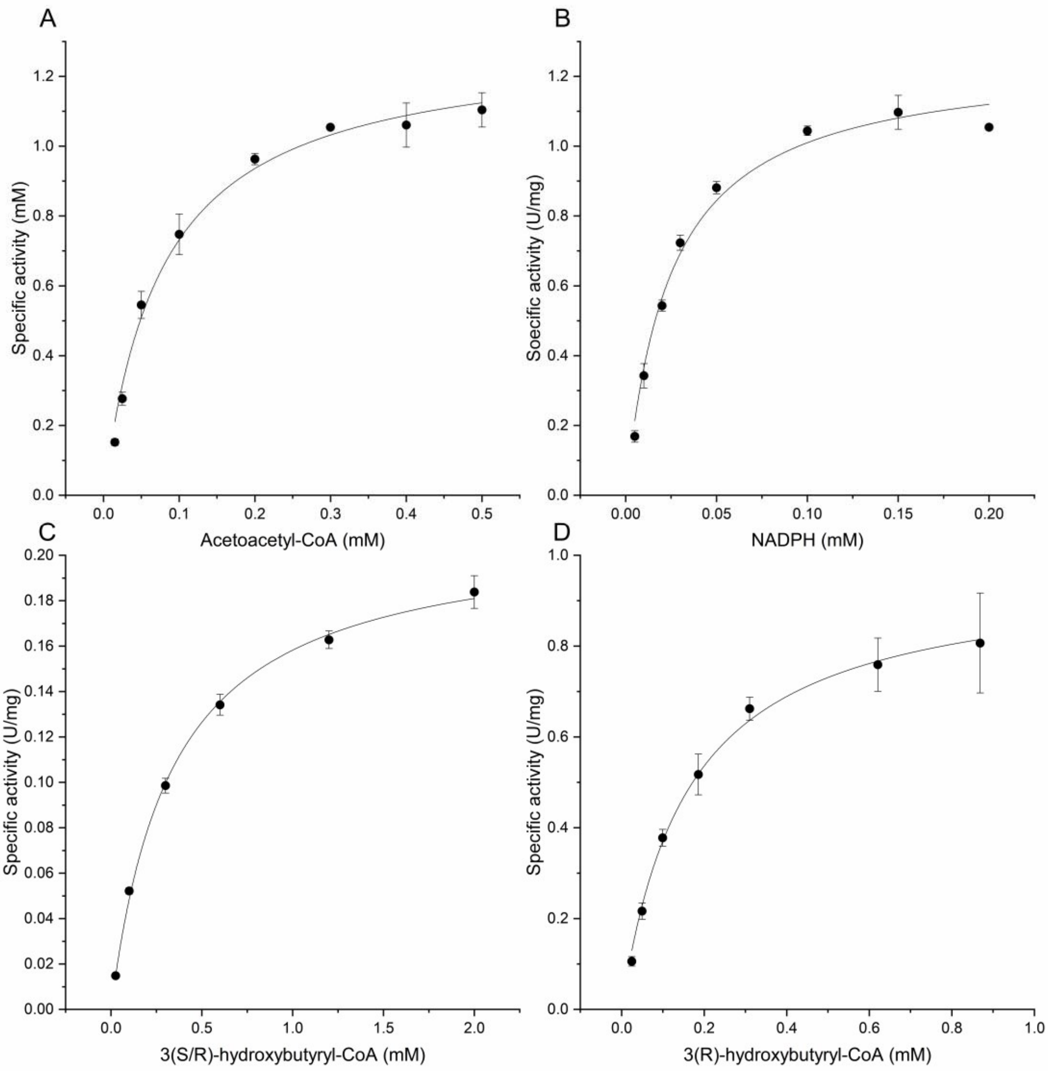


**Figure S24. Investigation of kinetic properties of the recombinant acetoacetyl(ketoacyl)-CoA reductase (ACR, Saci_1104) from *S. acidocaldarius*.** The ACR activity (total volume 500 µl) was determined in 100 mM TRIS-HCl (pH 7) in the reductive direction with 0.3 mM acetoacetyl-CoA (AcAcCoA), 0.2 mM NADPH and 0.00806 µg/µl protein at 35°C (340 nm). With NADH no activity was observed. The kinetic parameters were determined for AcAcCoA (A) and NADPH (B), respectively. In the oxidative direction, the activity was determined at 70°C using the commercially available, racemic mixture of 3(S/R)-hydroxybutyryl-CoA (HBCoA) or the pure stereoisomer 3(R)-HBCoA as substrate in presence of 2 mM $NADP^+$ (no activity observed with $NAD^+$) and 0.04032 µg/µl purified protein. The kinetic parameters for 3-HBCoA reduction were determined in presence of 0-2 mM 3(S/R)-HBCoA (C) or 0-1 mM 3(R)-HBCoA (D). Independent measurements were performed in triplicate and error bars indicate the standard error of the mean (SEM).

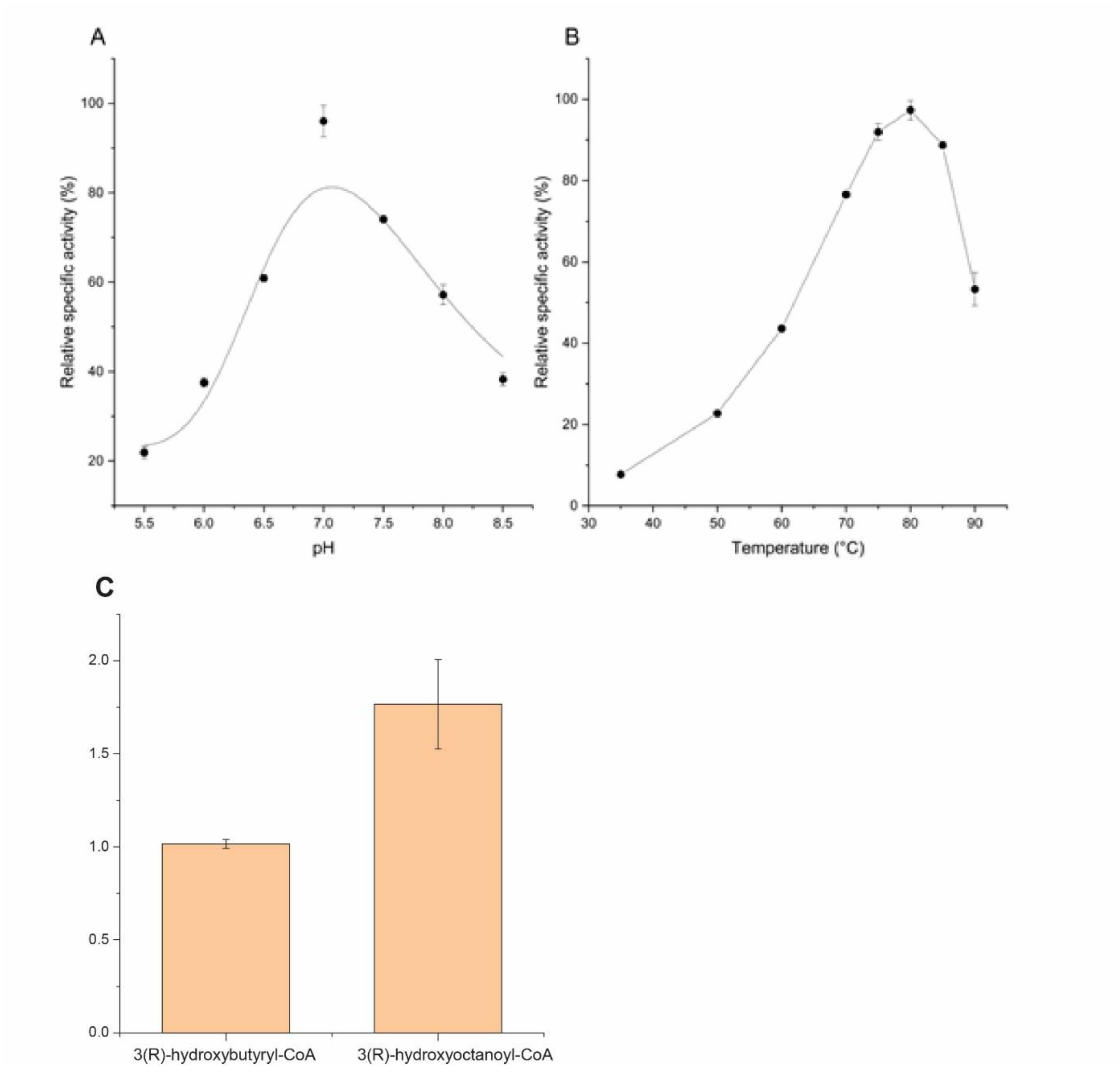


**Fig. S25. Investigation of the pH (A) and temperature (B) optimum and substrate specificity (C) of the recombinant acetoacetyl(ketoacyl)-CoA reductase (ACR, Saci_1104) from *S. acidocaldarius*.** The pH optimum was determined in the direction of 3-hydroxybutyryl-CoA (3-HBCoA) formation at 35°C using 100 mM MES-NaOH (pH 5.5-6.5) or 100 mM TRIS-HCl (pH 7.0-8.5) in presence of 0.3 mM acetoacetyl-CoA (AcAcCoA), 0.2 mM NADPH and 0.00806 µg/µl protein. The temperature optimum was determined in the direction of acetoacetyl-CoA formation using 100 mM TRIS-HCl (pH 7, at the respective temperature) in presence of 2 mM $NADP^+$, 0.3 mM 3(S/R)-hydroxybutyryl-CoA and 0.04032 µg/µl protein in a temperature range between 35-90°C. Independent measurements were performed in triplicate and error bars indicate the standard error of the mean (SEM).

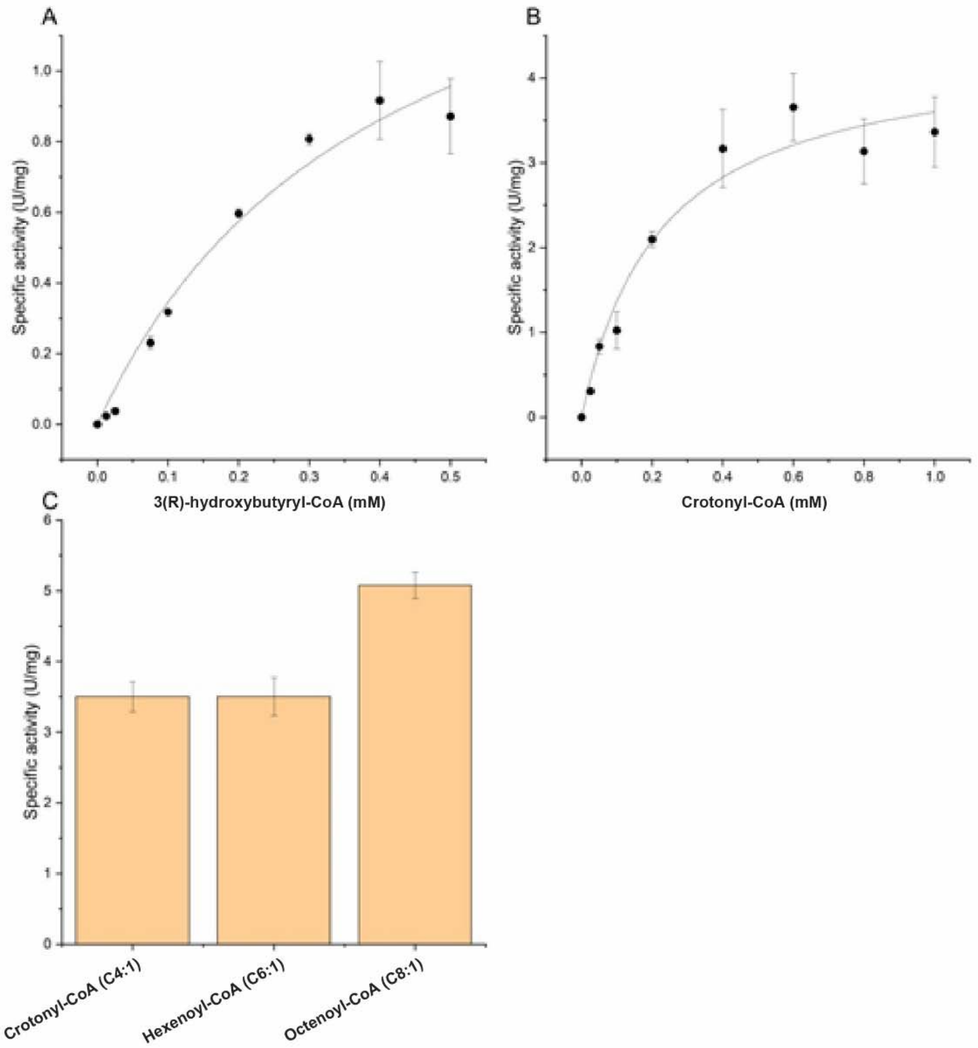


**Fig. S25. Investigation of kinetic properties (A, B) and substrate specificity (C) of the recombinant MaoC-like 3(R)-hydroxyacyl-CoA dehydratase (MaoC-HCD, Saci_1085) from *S. acidocaldarius*.** The activity of MaoC-HCD was tested at 65°C using a discontinuous assay (50 µl) in 50 mM MES-KOH (pH 6.5), 20 mM KCl and 0.0675 µg/µl protein. The kinetic parameters for 3(R)-hydroxybutyryl-CoA (0-0.5 mM (A)) and crotonyl-CoA (0-1 mM (B)) were determined. The chain length specificity was determined in presence of 0.09 µg/µl protein and 0.4 mM of the respective enoyl-CoA, crotonyl-CoA (C4), hexenoyl-CoA (C6) or octenoyl-CoA (C8). After incubation for 2, 5, 10, and 30 min, aliquots were withdrawn from the assay mixtures, and the reaction was stopped by addition of acetonitrile in a ratio of 1:3 (v/v) followed by freezing. The formation of the respective products was analysed via HPLC (C4 compounds 4-30% ACN HPLC program, compounds >C4, 1-60% ACN program) and the specific activities were calculated (se methods section). Independent measurements were performed in triplicate and error bars indicate the standard error of the mean (SEM).

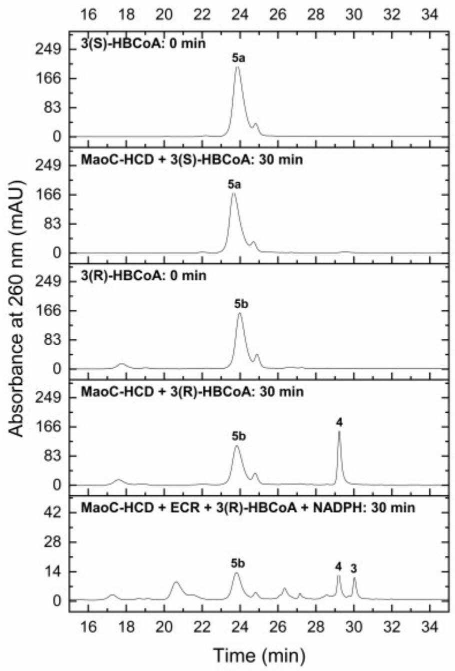


**Fig. S26. HPLC analysis of the stereospecificity of the recombinant MaoC-like 3(R)-hydroxyacyl-CoA dehydratase (MaoC-HCD, Saci_1085) (upper four chromatogramms) and the conversion of 3(R)-hydroxybutyryl-CoA (HBCoA) to butyryl-CoA by MaoC-HCD and enoyl-CoA reductase (ECR, Saci_1115) (bottom chromatogram).** To study the stereospecificity, 0.4 mM of pure stereoisomers 3(S)- or 3(R)-HBCoA were incubated with 0.0675 µg/µl Saci_1085 in 50 mM MES-KOH (pH 6.5) and 20 mM KCl at 65°C for 30 min followed by HPLC analyses of the samples. 3(S)-HBCoA (peak 5a) was not converted (first and second chromatogram from top) while 3(R)-HBCoA (peak 5b) was converted to crotonyl-CoA (peak 4) by MaoC-HCD (third and fourth chromatogram from top). Chromatogram 1 and 3 (from top) represent the reference state before addition of enzyme (0 min). -- The two step butyryl-CoA formation from 3(R)-HBCoA via crotonyl-CoA was carried out by MaoC-HCD (Saci_1085) and ECR (Saci_1115) (bottom chromatogram). The assay was performed at 70°C in 50 mM MES-KOH (pH 6.5), 20 mM KCl, 2 mM NADPH with 0.0246 µg/µl ECR and 0.0168 µg/µl MaoC-HCD, and was incubated for 30 min. Crotonyl-CoA (peak 4) and butyryl-CoA (peak 3) formation from 3(R)-HBCoA (peak 5b), respectively, was observed (bottom chromatogram). All assay samples were analyzed using the 4-30% ACN HPLC program. For assignment of peak numbers to compounds and HPLC programs see Fig. S13.

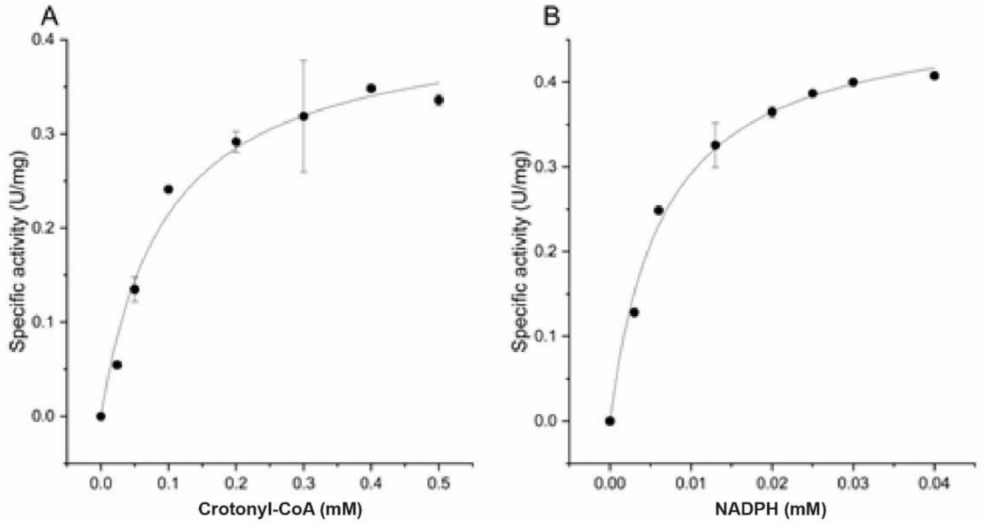


**Fig. S27. Investigation of kinetic parameters of the recombinant enoyl-CoA reductase (ECR, Saci_1115) from *S. acidocaldarius*.** The ECR activity (total volume 500 µl) was determined photometrically at 70°C (340 nm) with 0.024 µg/µl of protein in 100 mM HEPES-NaOH (pH 7.5), 10 mM KCl, 0.3 mM NADPH and 0.4 mM crotonyl-CoA. The kinetic parameters were determined in the presence of 0-0.5 mM crotonyl-CoA (A) and 0-0.4 mM NADPH (B). Independent measurements were performed in duplicate and error bars indicate the standard error of the mean (SEM).

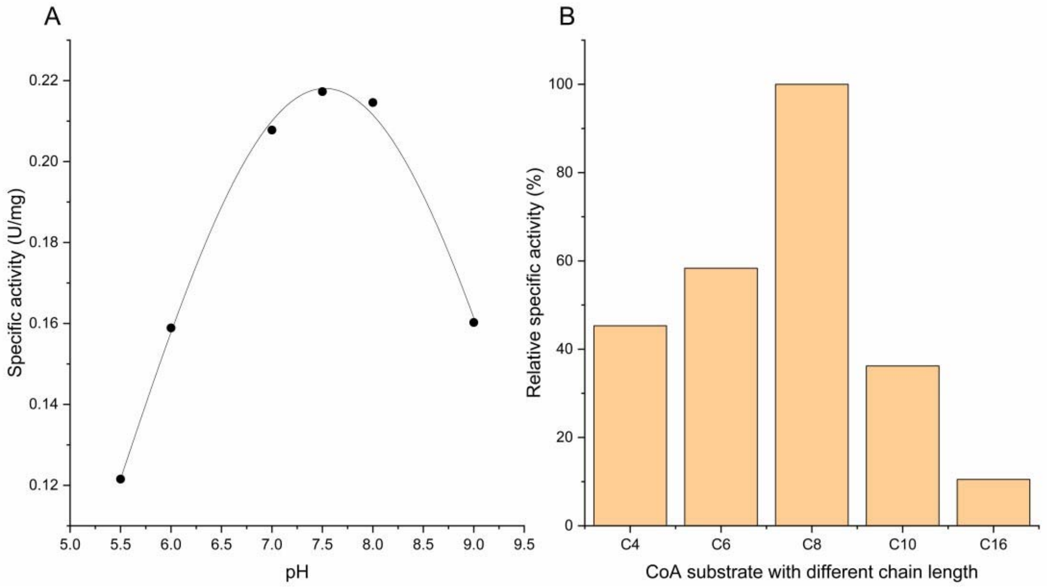


**Fig. S28. Investigation of the pH optimum (A) and substrate specificity (B) of the recombinant enoyl-CoA reductase (ECR, Saci_1115) from *S. acidocaldarius*.** The pH optimum for ECR was determined at 70°C in a pH range of 5.5-9.0 using a mixed buffer system of 0.5 M HEPES, 0.5 M TRIS and 0.5 M MES in presence of 10 mM KCl, 0.4 mM crotonyl-CoA, 0.3 mM NADPH and 0.016 µg/µl enzyme (total volume 500 µl). The substrate specificity of ECR towards different chain length enoyl-CoAs, i.e. crotonyl-CoA (C4), hexenoyl-CoA (C6), octenoyl-CoA (C8), decenoyl-CoA (C10) or hexadecenoyl-CoA (C16), was determined in 100 mM HEPES-NaOH (pH 7.5, 70°C) with 0.3 mM of the respective enoyl-CoA, 0.2 mM NADPH and 0.02 µg/µl ECR.

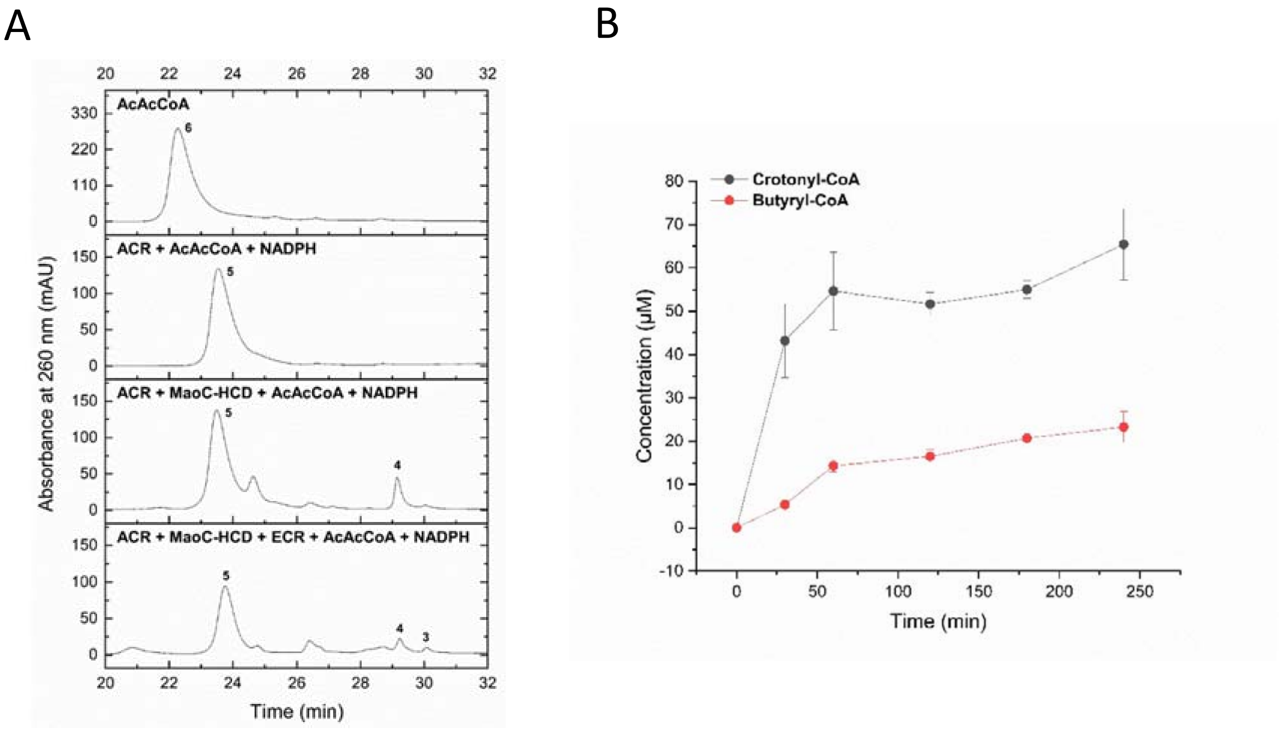


**Figure S29. Acetoacetyl-CoA conversion to butyryl-CoA by the recombinant acetoacetyl(ketoacyl)-CoA reductase (ACR, Saci_1104), MaoC-like 3(R)-hydroxyacyl-CoA dehydratase (MaoC-HCD, Saci_1085), and enoyl-CoA reductase (ECR, Saci_1115).** In **(A)** the consecutive build-up of the three-step enzyme cascade is shown: All conversions (total volume 20 µl were carried out discontinuously in 50 mM MES-KOH (pH 6.5) with 20 mM KCl, for 30 min. After incubation samples were analyzed for substrate and intermediate/product formation via HPLC. First chromatogram from top: Reference state containing 0.4 mM acetoacetyl-CoA (AcAcCoA, peak 6) before addition of enzyme(s) and 2 mM NADPH; second chromatogram from top: After incubation with 0.052 µg/µl ACR and 2 mM NADPH at 65°C, AcAcCoA is completely converted to 3(R)-hydroxybutyryl-CoA (HBCoA); third chromatogram from top: Incubation of AcAcCoA in the presence of 0.104 µg/µl ACR, 2 mM NADPH, and 0.0168 µg/µl MaoC-HCD at 65°C resulted in the formation of 3(R)-HBCoA (peak 5) as main product but also lower amounts of crotonyl-CoA (peak 4) which coincides well with the thermodynamics of both reactions (see supplementary text); bottom chromatogram: After incubation of AcAcCoA with 0.104 µg/µl ACR, 2 mM NADPH, and 0.0168 µg/µl MaoC-HCD, and 0.0246 µg/µl ECR at 65°C, butyryl-CoA (peak 3) was detected at least in low amounts in addition to 3(R)-HBCoA and crotonyl-CoA demonstrating the functional three enzyme cascade *in vitro*. In (B) the time-dependent conversion of acetoacetyl-CoA to crotonyl-CoA and further to butyryl-CoA via ACR, MaoC-HCD and ECR was monitored over 240 min under same conditions described above. At the time point indicated aliquots were withdrawn, analyzed via HPLC, and crotonyl-CoA and butyryl-CoA formation was quantified via peak areas. Approximately 70 µM crotonyl-CoA (black) and 20 µM butyryl-CoA (red) were produced within the observed time frame. The low amounts of butyryl-CoA may be explained by the low activity of the ECR (see main text). For assignment of peak numbers to compounds and HPLC programs see Fig. S13.

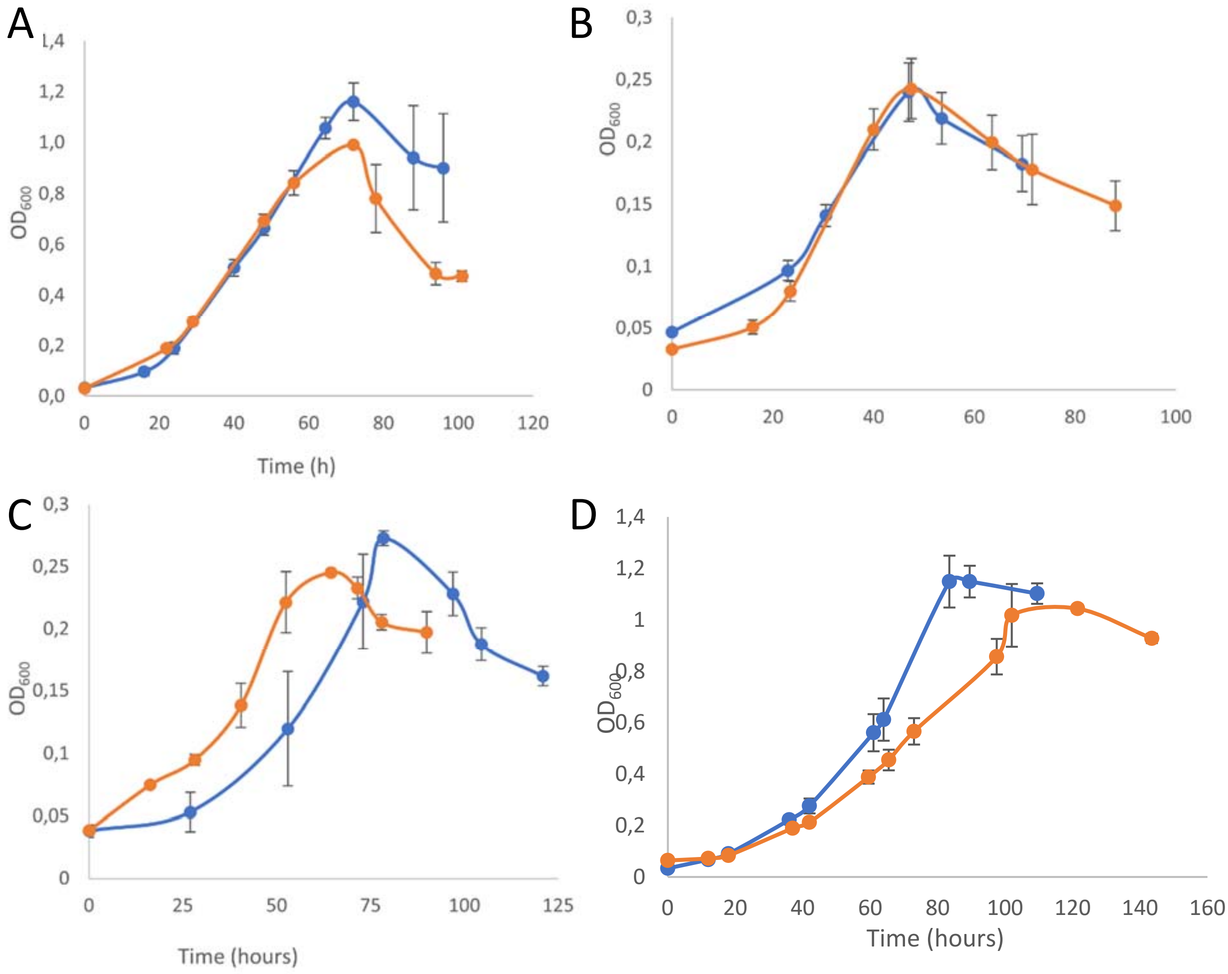


**Figure S30. Comparative growth studies of the S*. acidocaldarius* MW00G parental strain and the enoyl-CoA reductase (ECR) deletion strain Δ1115** on basal Brock medium with 0.2% (w/v) D-xylose (A), 2 mM butyrate (B), and 2 mM hexanoate (C), and 10 mM glycerol (D) as sole carbon and energy source. Growth was quantified as increase in OD at 600 nm over time. Parental strain *S. acidocaldarius* MW00G is depicted in blue and the Δ1115 deletion strain in orange.

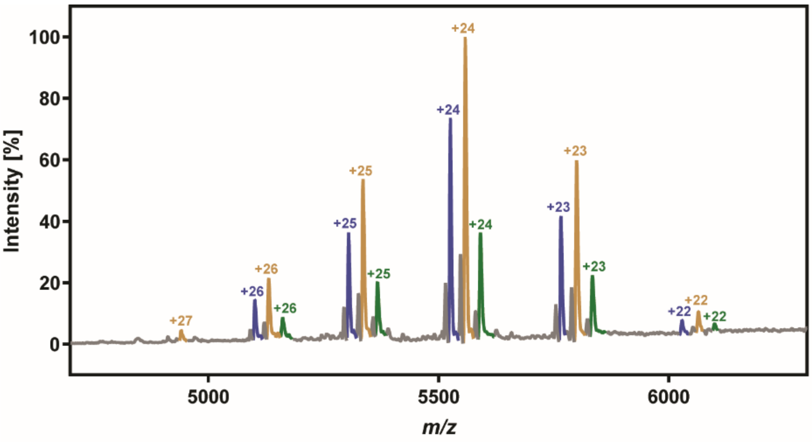


| # | Mass [Da] | Int. [%] |
|---|---|---|
| 1 | 132,603 ± 5 | 32.7 ± 0.5 |
| 2 | 133,384 ± 6 | 52.6 ± 1.4 |
| 3 | 134,174 ± 4 | 15.7 ± 1.1 |

**Figure S31. Determination of the molecular mass of the recombinant β-ketothiolase/DUF35 domain protein (KT/DUF35; Saci_1121/1120) complex via nMS.** A representative averaged MS spectrum of 1000 scans of the KT/DUF35 complex (1 µM in 2.5 mM $NH_4OAc$) was color-coded to indicate the peaks that originated from the same mass species and the respective deconvoluted charge states are presented at the top of the peaks. The mean deconvoluted protein masses of three technical replicates, calculated by UniDec, are provided in the right table along with their standard error and relative intensity.

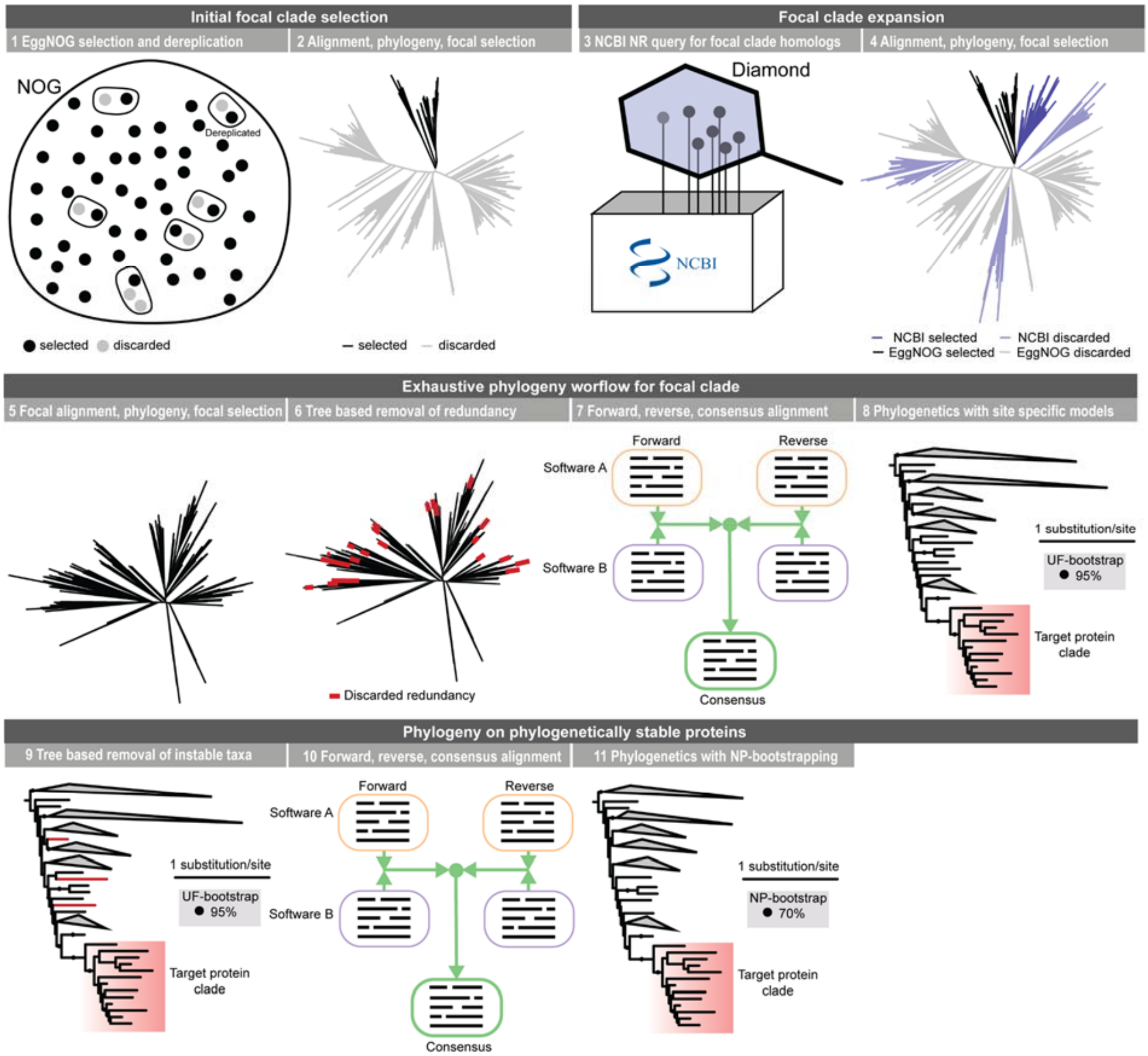


**Figure S32. Phylogenetic workflow to study fatty acid synthesis protein evolutionary history.**

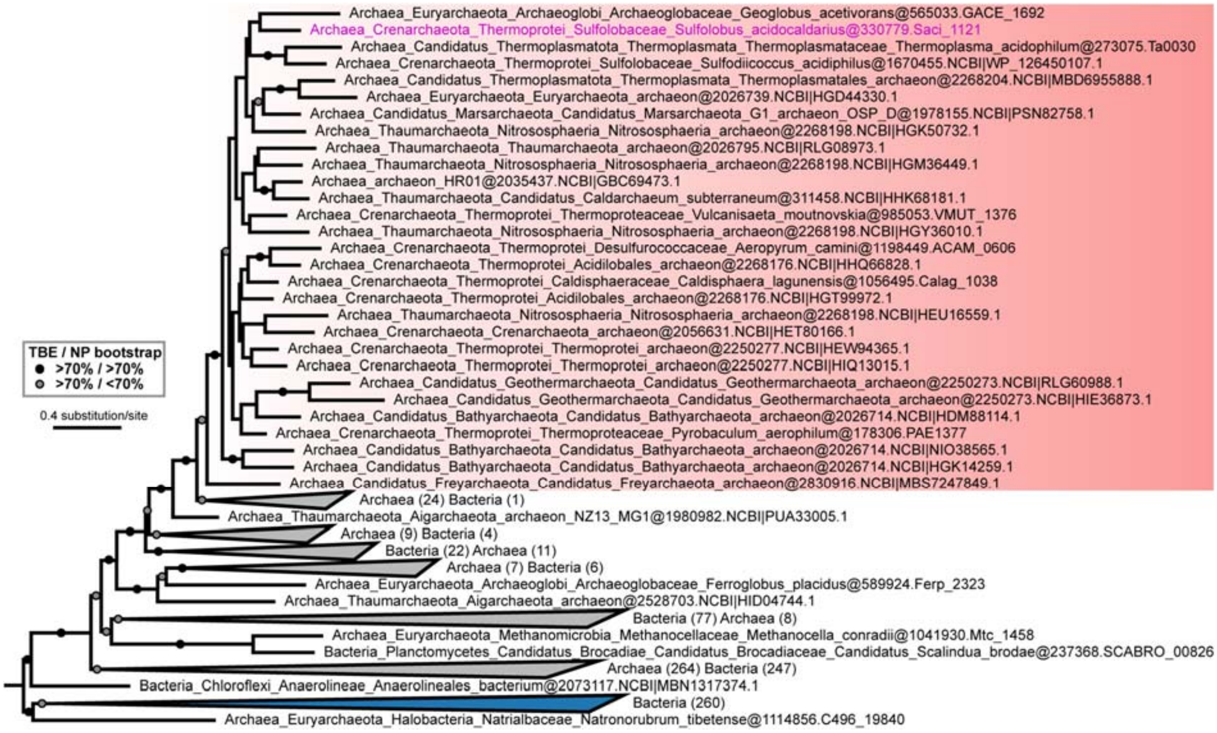


**Figure S33. Maximum-likelihood (ML) tree of the ketothiolase Saci_1121 (COG0183)** under the LG+C60+R6+F+PMSF model of evolution with 100 NP bootstraps and TBE support. Based on 22,646 initial protein sequences.

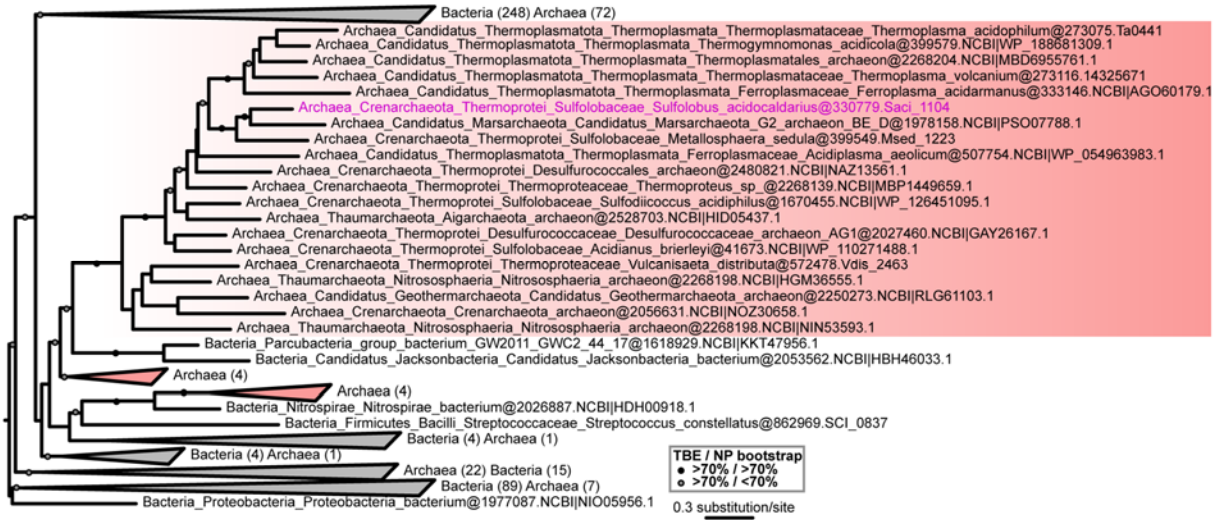


**Figure S34. ML tree of the Acetoacyl-CoA reductase Saci_1104 (COG1028)** under the LG+C60+R6+F+PMSF model of evolution with 100 NP bootstraps and TBE support. Based on 121,803 initial protein sequences.

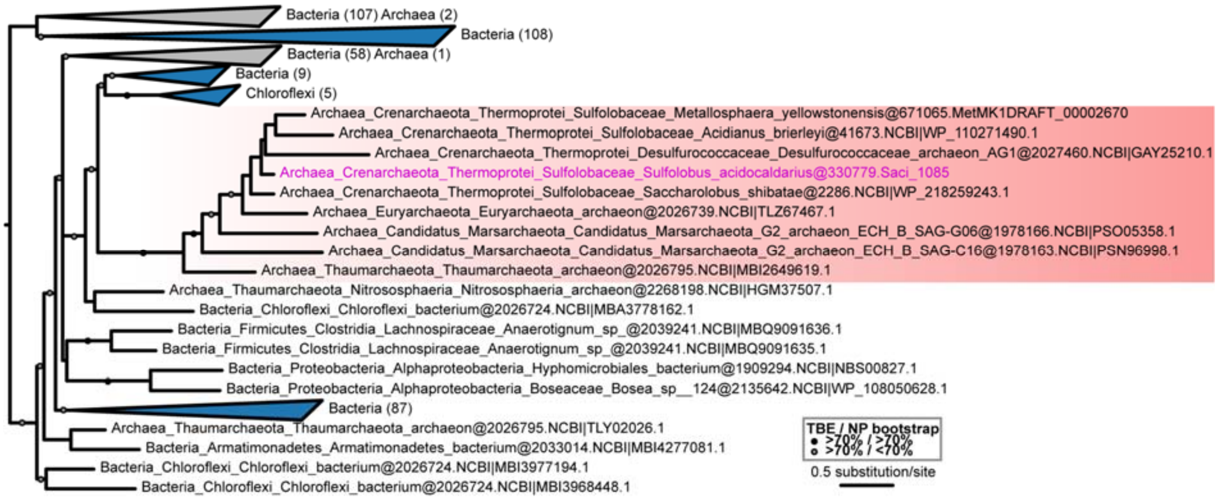


**Figure S35. ML tree of the (R)-hydroxyacyl-CoA dehydratase Saci_1085 (COG2030)** under the LG+C60+R6+F+PMSF model of evolution with 100 NP bootstraps and TBE support. Based on 12,511 initial protein sequences.

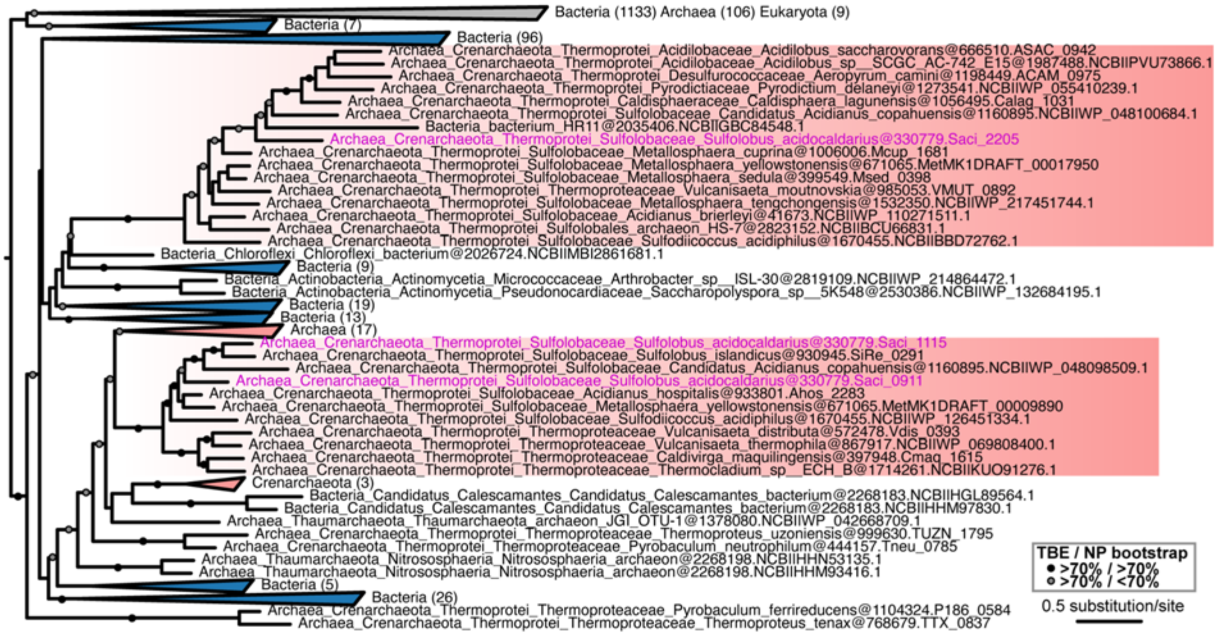


**Figure S36. ML tree of the Enoyl-CoA reductase Saci_1115 (COG1064)** under the LG+C60+R6+F+PMSF model of evolution with 100 NP bootstraps and TBE support. Based on 8,512 initial protein sequences.

**Table S1. Molecular and kinetic parameters of the β oxidation enzymes in *S. acidocaldarius*.**

| ORF/Enzyme | Molecular mass (kDa) | Temp | Substrate/Cofactor | Km (mM) | Vmax (U $mg^{-1}$ ) | kcat ($S^{-1}$ ) |
|---|---|---|---|---|---|---|
| **Saci_1123 Acyl-CoA dehydrogenase (ACAD)** | 44 (subunit) 170 (native) homotetramer | 65°C | Butyryl-CoA (C4) | NM | 4.5 | 3.29 |
| | | | Hexanoyl-CoA (C6) | NM | 8.24 | 6.02 |
| | | | Octanoyl-CoA (C8) | 0.0151 ± 0.004 | 29.046 ± 2.048 | 21.234 |
| | | | Palmitoyl-CoA (C16) | NM | ND | ND |
| **Saci_0315 Electron transfer flavoprotein (ETF)** | 68 (subunit) 67 (native) monomer | 65°C | NADH | 0.039 ± 0.012 | 0.703 ± 0.046 | 0.784 |
| **Saci_1109 Enoyl-CoA hydratase/3-hydroxyacylCoA dehydrogenase (ECH/HCDH)** | 73 (subunit) 512 (native) homoheptamer | 75°C | Aryloyl-CoA (C3:1) | NM | [a] 0.073 | 0.088 |
| | | | Crotonyl-CoA (C4:1) | 0.024 ± 0.004 | 16.974 ± 0.479 | 20.56 |
| | | | $NAD^+$ | 0.036 ± 0.009 | | |
| | | | $NADP^+$ | NM | ND | ND |
| | | | Decenoyl-CoA (C10:1) | NM | [a] 7.6 | 9.21 |
| | | | Hexadecenoyl-CoA (C16:1) | NM | ND | ND |
| | | | 3(S/R)-Hydroxybutyryl-CoA C4) | 0.092 ± 0.016 | 29.658 ± 1.446 | 35.923 |
| | | | NAD+ | 0.11 ± 0.012 | | |
| | | | NADP+ | NM | ND | ND |
| | | | 3(S)-Hydroxybutyryl-CoA (C4) | 0.043 ± 0.002 | 48.439 ± 0.398 | 58.672 |
| | | | 3(R)-Hydroxybutyryl-CoA (C4) | NM | ND | ND |
| | | 35°C | Acetoacetyl-CoA (C4) | 0.076 ± 0,008 | 6.969 ± 0.191 | 8.441 |
| | | | NADH | 0.028 ± 0.007 | | |
| | | | NADPH | *0.094 ± 0.22 | | |
| **Saci_1114 β-Ketothiolase (KT)** | 88 (subunit) 88 (native) homodimer | 23°C | Acetoacetyl-CoA (C4) | 0.033 ± 0.01 | 1.676 ± 0.25 | 1.2 |
| | | 75°C | CoA | 0.00338 | 2.53375 | 1.8 |
| | | | Acetyl-CoA (C2) | 2.097 ± 0.263 | 2.739 ± 0.139 | 1.959 |

* The kinetic parameters were calculated in regardless of inhibition effect caused by higher concentration of the substrate.
Numbers after ± represent standard error (SE).
ND: not detectable.
NM: not measured
[a] The values of these specific activities were estimated according to the experimental measurements.

**Table S2. Molecular and kinetic parameters of the enzymes involved in the new potential fatty acid synthesis pathway in *S. acidocaldarius*.**

| ORF/Enzyme | Molecular mass (kDa) | Temp | Substrate/Cofactor | Km (mM) | Vmax (U mg$^{-1}$) | kcat (S$^{-1}$) |
|---|---|---|---|---|---|---|
| **Saci_1104 Acetoacetyl (ketoacyl)-CoA reductase (ACR)** | 27 (subunit) 84 (native) homotrimer | 35°C | Acetoacetyl-CoA (C4) | 0.077 ± 0.009 | 1.3 ± 0.042 | 0.586 |
| | | | NADPH | 0.024 ± 0.003 | | |
| | | 70°C | 3(S/R)-Hydroxybutyryl-CoA (C4) | 0.336 ± 0.02 | 0.211 ± 0.004 | 0.095 |
| | | | 3(S)-Hydroxybutyryl-CoA (C4) | NM | ND | ND |
| | | | 3(R)-Hydroxybutyryl-CoA (C4) | 0.16 ± 0.012 | 0.965 ± 0.024 | 0.435 |
| | | | 3(R)-Hydroxyoctanoyl-CoA (C4) | NM | 1.77 | 0.80 |
| **Saci_1085 MaoC like 3(R)-hydroxacyl-CoA dehydratase (MaoC-HCD)** | 19 (subunit) 122 (native) homohexamer | 65°C | 3(S)-Hydroxybutyryl-CoA (C4) | NM | ND | ND |
| | | | 3(R)-Hydroxybutyryl-CoA (C4) | 0.399 ± 0.127 | 1.72 ± 0.301 | 0.549 |
| | | | Crotonyl-CoA (C4:1) | 0.222 ± 0.064 | 4.398 ± 0.414 | 1.404 |
| | | | Hexenoyl-CoA (C6:1) | NM | 3.5 | 1.12 |
| | | | Octenoyl-CoA (C8:1) | NM | 5.1 | 1.63 |
| **Saci_1115 Enoyl-CoA reductase (ECR)** | 36 (subunit) 69 (native) homodimer | 70°C | Aryloyl-CoA (C3:1) | NM | ND | ND |
| | | | Crotonyl-CoA (C4:1) | 0.096 ± 0.018 | 0.422 ± 0.024 | 0.256 |
| | | | NADPH | 0.007 ± 0.00076 | | |
| | | | NADH | NM | 0.077 | 0.047 |
| | | | [b] Hexenoyl-CoA (C6:1) | NM | [a] 0.54 | 0.33 |
| | | | [b] Octenoyl-CoA (C8:1) | NM | [a] 0.93 | 0.56 |
| | | | Decenoyl-CoA (C10:1) | NM | [a] 0.337 | 0.2 |
| | | | Hexadecenoyl-CoA (C16:1) | NM | [a] 0.098 | 0.059 |
| **Saci_1121/1120 Ketothiolase/ DUF35 (KT/DUF35)** | 43 (subunit Saci_1121) 20 (subunit Saci_1120) 133 (native) heterotetramer | 23°C | Acetoacetyl-CoA (C4) | 0.065 ± 0.0187 | 3.642 ± 0.453 | 4.024 |
| | | | CoA | 0.020 ± 0.0041 | 3.138 ± 0.200 | 3.467 |
| | | 70°C | Acetyl-CoA (C2) | 0.014 ± 0.002 | 0.848 ± 0.026 | 0.937 |
| | | | Butyryl-CoA (C4) | NM | 1.471 ± 0.040 | 1.625 |

Numbers after ± represent standard error (SE).
ND: not detectable.
NM: not measured

**Table S3. Strains, oligonucleotides and plasmids used in this study.**

| **Strains** | **Function** | **Source/Reference** |
|---|---|---|
| *E. coli* DH5 *α* | Plasmid construction | Hanahan, USA |
| *E. coli* Rosetta (DE3) | Heterologous gene expression | Stratagene, USA |
| *S. acidocaldarius* MW001 | Homologous gene expression | **(*19*)** |
| *S. acidocaldarius* MW00G | Glycerol adaption strain, Fatty acid detection | **(*20*)** |
| *S. acidocaldarius* MW00G Δ*saci_1115* | Glycerol adaption strain Δ*saci_1115,* Fatty acid detection | This work |
| *Haloferax volcanii H26* | Fatty acid detection | **(*21*)** |
| **Primers** | Sequences (5´→3´) | |
| *saci_1123*_fw_*Hind*III | ATAAAGCTTATGGTTTTGCCTTTTAAAAC | |
| *saci_1123*_rv_*Hind*III | ATAAAGCTTTTACATTTTTATGCCAAATAA | |
| *saci_0315*_fw | TAGCAGCCGGATCCTCGAGCAGCCCCTTCTTTTTAATTAAC | |
| *saci_0315*_rv | CCTGGTGCCGCGCGGCAGCCATATGGCAGAGCTTAAAATTGTCG | |
| *saci_1109*_fw_ *Bam*HI | TATGGATCCATGAAAGTAGAAGATATTAAGAAA | |
| *saci_1109*_rv_ *Bam*HI | GATGGATCCTTATTCTCCTTTGAACTGTG | |
| *saci_1104*_fw_*Nde*I | GCTCGCCATATGTACTCTCTTAAAGAC | |
| *saci_1104*_rv_*Bam*HI | CTAGCTGGATCCTTAAGCAATTCCT | |
| *saci_1115*_fw_*Nde*I | CCTACGCATATGATGAAAGCTGTAATTCTTC | |
| *saci_1115*_rv_*Bam*HI | CGAGCTGGATCCTTATGGCTTTATAAGAATTTTAC | |
| *saci_1085*_fw_*Nco*I | GCCCGCCATGGGGTCAGAGCAGGGTCC | |
| *saci_1085*_rv_*Xho*I | CGGGCCTCGAGTCATTGTGGTTTGTCAGTAC | |
| *saci_1121/saci_1120*_fw_*Nco*I | GAGAGCCATGGGAGAAACGTGGCAATTG | |
| *saci_1121/saci_1120*_rv_*Bam*HI | GAGAGGGATCCGACTACATTGAAGACATAG | |
| *saci_1115*_ol_up_fw_*Bam*HI | TATGGATCCTGTGGATGCCACAAAGTGG | |
| *saci_1115*_ol_up_rv | CTCACTATTAGTTCAATTAAATATAAA**C**AAACAAAAATAATTAAAATTTTG | |
| *saci_1115*_ol_down_fw | GTTTATATTTAATTGAACTAATAGTGA**G**GAAAAATAGAAAAACTTC | |
| *saci_1115*_ol_down_rv_*Sal*I | TAAGTCGACTCAATTACCTCCAGCATTTAG | |
| **Oligonucleotides for sequencing** | **Sequences (5´→3´)** | |
| T7-promoter | TAATACGACTCACTATAGGG | |
| T7-terminator | GCTAGTTATTGCTCAGCGG | |
| FXara-fw | CAGCGTTTATAACGTTTAACATG | |
| FXsulf-rv | CCATTTAATAGTTTGTATGGTCTACCC | |
| **Plasmids** | **Genotype or description** | **Source/Reference** |
| pET15b | *E. coli* expression plasmid carrying an N-terminal his-tag for cloning/expression of *saci_1104, saci_1115 and saci_0315* | Novagen, USA |
| pET28b | *E. coli* expression plasmid carrying both an N-terminal and a C-terminal his-tag for cloning/expression of *saci_1114* | Novagen, USA |
| pET45b | *E. coli* expression plasmid carrying an N-terminal his-tag for cloning/expression of *saci_1123, saci_1109* | Novagen, USA |
| pSVA407 | Gene targeting plasmid, pGEM-T Easy backbone, *pyrEF* cassette of *S. solfataricus* | (*19*) |
| pBS-araFX-UTR-NtSS | *S. acidocaldarius* expression plasmid carrying a N-terminal twin-strep-tag for cloning/expression of *saci_1085* | (*22*) |
| pBS-araFX-UTR-CtSS | *S. acidocaldarius* expression plasmid carrying a C-terminal twin-strep-tag for cloning/expression of *saci_1121/saci_1120* | (*22*) |

**Table S4. Expression conditions used for the selected genes from *S. acidocaldarius*.**

| Enzymes | ORFs | Overexpression conditions |
|---|---|---|
| Acyl-CoA dehydrogenase (ACAD) | *saci_1123* | 0.4 mM IPTG for induction; 20°C overnight |
| Electron transfer flavoprotein (ETF) | *saci_0315* | 0.4 mM IPTG for induction; 20°C overnight |
| Enoy-CoA hydratase/3-hydroxyacyl-CoA dehydrogenase (ECH/HCDH) | *saci_1109* | 1 mM IPTG for induction; 30°C overnight |
| β-Ketothiolase/Acetyl-CoA acetyltransferase (KT/ACAT) | *saci_1114* | 1 mM IPTG for induction; 22°C overnight |
| Acetoacetyl(ketoacyl)-CoA reductase (ACR) | *saci_1104* | 1 mM IPTG for induction; 37°C 4 hours |
| Enoyl-CoA reductase (ECR) | *saci_1115* | 1 mM IPTG for induction; 16°C overnight |
| MaoC-like 3(R)-hydroxyacyl-CoA dehydratase (MaoC-HCD) | *saci_1085* | 0.3% D-xylose for induction; 75°C 48 hours |
| β-Ketothiolase/Acetyl-CoA acetyltransferase (KT/ACAT)<br>DUF35 | *saci_1121/*<br>*saci_1120* | 0.3% D-xylose for induction; 75°C 48 hours |

**Table S5. Retention times of the corresponding CoA ester compounds in different HPLC running programs.** The relevant HPLC chromatographs are shown in Fig. S13 (ND: not detected or not detectable).

| Peak no. | Compound | Retention time (min) | |
|---|---|---|---|
| | | in 4-30% ACN program | in 1-60% ACN program |
| 1 | CoA | 12.5 | 8.2 |
| 2 | Acetyl-CoA | 21.5 | 9.7 |
| 3 | Butyryl-CoA | 29.9 | 25.5 |
| 4 | Crotonyl-CoA (C4:1) | 29 | ND |
| 5 | 3-Hydroxybutyryl-CoA | 23.5 | ND |
| 5a | 3-(S)-Hydroxybutyryl-CoA | 23.5 | ND |
| 5b | 3-(R)-Hydroxybutyryl-CoA | 23.5 | ND |
| 6 | Acetoacetyl-CoA | 22.5 | ND |
| 7 | Hexanoyl-CoA | 35.5 | 31 |
| 8 | Octanoyl-CoA | ND | 34 |
| 9 | Hexenoyl-CoA (C6:1) | ND | 30.3 |
| 10 | 3-Hydroxyhexanoyl-CoA | ND | 24.5 |
| 11 | Octenoyl-CoA (C8:1) | ND | 33.9 |
| 12 | 3-Hydroxyoctanoyl-CoA | ND | 30.6 |

**Table 6. Software settings used for UniDec analysis of MS data.**

| **Data Processing** | |
|---|---|
| **m/z range** | 4,000-7,000 |
| Background Subtraction | Yes |
| **UniDec Parameters** | |
| Mass Range [Da] | 10,000-200,000 |
| Sample Mass Every [Da] | 1 |
| Point Smooth Width | 1-3 |
| **Peak Selection** | |
| Peak Detection Threshold | 0.1 |
| Peak Normalization | Total |
| **Additional Filtering** | |
| Filter Peak Scores [% DScore] | 35 |
